# *In vivo* detection of protein-protein interactions with single molecule resolution

**DOI:** 10.1101/2024.09.06.611644

**Authors:** Marilyne Davi, Daniel Ladant

## Abstract

Protein-protein interactions are central in all biological processes. Methods capable of detecting interactions within living, intact cells have been particularly useful to identify and characterize protein interaction networks. We describe here an exquisitely sensitive regulatory circuit that can detect in bacteria, protein-protein interaction with single molecule sensitivity. This approach involves the interaction-mediated reconstitution of a cyclic AMP signaling cascade in *Escherichia coli* taking advantage of the high catalytic activity of the adenylate cyclase (AC) from *Bordetella pertussis* upon activation by its natural activator, calmodulin (CaM). We show that a single complex of interacting hybrid proteins per cell is enough to confer a selectable trait to the host. This <u>e</u>xquisitely <u>s</u>ensitive <u>a</u>denylate <u>c</u>yclase <u>h</u>ybrid (ESACH) system allows for direct *in vivo* selection of ligands exhibiting high affinity for given targets or for studying interactions involving toxic proteins. The extreme sensitivity of the AC/CaM/cAMP signaling cascade may thus be harnessed to interrogate biological processes with single molecule resolution in live bacteria and could be exploited to design novel synthetic regulatory networks operating at, or even below, the theoretical threshold limit of one molecule per cell.

## INTRODUCTION

Most cellular processes are driven by molecular interactions and the study of protein interaction networks represents an essential step to understand biological systems. Methods capable of detecting protein-protein interactions (PPI) within living, intact cells have been particularly useful to identify and characterize the “interactome”, a cornerstone of systems biology. Most of these methods are based on the co-expression, in the same cell, of two (or more) hybrid proteins that, upon interaction, produce a phenotypic and/or selective trait. This approach was pioneered by Fields and Song, who developed the yeast two-hybrid system (or interaction trap), in which the putative interaction partners are co-expressed as fusions with two moieties of a transcriptional activator ^1^. Association of the two hybrid proteins restores the activity of the split transcriptional activator and leads to the expression of specific reporter genes. Similar approaches have later been designed in which split enzymes or fluorescent/luminescent proteins are used as reporter systems to select or image *in vivo* protein-protein interactions in various hosts, such as yeast, mammalian or bacterial cells ^2^ ^3 4 5 6 7 8 9 10 11 12 13 14 15 16 17 18 19 20^.

We previously elaborated a bacterial two-hybrid system, BACTH (Bacterial Adenylate Cyclase Two-Hybrid) that is based on the interaction-mediated reconstitution of a cyclic AMP (cAMP) signaling cascade in *Escherichia coli* ^21^. In this system, the proteins of interest are genetically fused to two complementary fragments, T25 and T18, from the catalytic domain of *Bordetella pertussis* adenylate cyclase (AC), and co-expressed in an *E. coli cya* strain (i.e. deficient in its endogenous adenylate cyclase). Association of the two hybrid proteins results in functional complementation between the separately inactive T25 and T18 fragments leading to cAMP synthesis. In *E. coli*, cAMP binds to the catabolite activator protein (CAP or CRP) and triggers the transcriptional activation of catabolic operons, such as lactose or maltose, thus yielding a characteristic phenotype. This system has been extensively used to reveal a wide variety of interactions between bacterial, eukaryotic, or viral proteins, occurring at various subcellular locations, e.g. cytosol, membrane or DNA level ^22^ ^23^ ^24^ ^25^.

Here, we describe a novel architecture for the AC-based two-hybrid screen that permits *in vivo* detection of PPI with an extreme sensitivity. This approach takes advantage of the high catalytic potency of *B. pertussis* AC upon activation by its natural activator, the eukaryotic calcium-sensor, calmodulin (CaM) ^26^. With this new design, we found that, on average, less than one complex of interacting hybrid proteins per cell is enough to confer a selectable trait to the host. This exquisitely sensitive adenylate cyclase hybrid (ESACH) system should be particularly appropriate for direct *in vivo* selection of ligands exhibiting high affinity for given targets or for studying PPI involving toxic proteins. The ultimate sensitivity provided by the AC/CaM/cAMP signaling cascade that can reliably detect one active enzyme complex per bacterium, could also be exploited to engineer novel synthetic regulatory networks operating at, or even below, the theoretical threshold limit of one molecule per cell.

## RESULTS

### Design of an exquisitely sensitive adenylate cyclase hybrid (ESACH) system

Our previously described AC-based two-hybrid system^21^ relies on the interaction-mediated reconstitution of enzymatic activity from two complementary fragments of *B. pertussis* AC. In the absence of its activator calmodulin (CaM), AC exhibits a k_cat_ of about 1-2 s^-1^, and therefore few hundreds of active hybrid protein complexes per bacteria are required to produce enough cAMP to confer a Cya*^+^* phenotype to the *E. coli Δcya* host cells. Upon binding to CaM, AC is activated more than a 1000 fold ^26^ to reach a turnover number of about 2000 s^-1^. We reasoned that by taking advantage of the full catalytic potency of AC upon activation by CaM, one should drastically increase the sensitivity of this genetic assay, so that a single active AC/CaM complex per cell might potentially be sufficient to confer a selectable trait to the bacterial host.

In the novel architecture explored here, named ESACH for <u>E</u>xquisitely <u>S</u>ensitive <u>A</u>denylate <u>C</u>yclase <u>H</u>ybrid, the two proteins of interest are separately fused to AC and CaM and co-expressed in an *E. coli Δcya* strain (Fig. 1). To render the AC activation dependent upon the association of the hybrid proteins, AC was engineered to reduce its naturally very high affinity for CaM (the CaM concentration required for half-maximal activation, K_1/2_, is about 0.1-0.2 nM in the presence of calcium ^27^) by introducing appropriate mutations, so that when both the modified AC and CaM are expressed alone at low level in a *E. coli Δcya* strain, they cannot spontaneously associate. Among the various modifications known to decrease CaM affinity, we chose two, ACM1 and ACM2, that result from two-amino acid insertions within the T18 moiety of AC: Leu-Gln between residues 247 and 248 in ACM1, and Cys-Ser between residues 335 and 336 in ACM2 ^27^. These insertions were previously shown to decrease CaM affinity by more than 3,000 and 50-100 fold, respectively ^27^.These two-codon insertion mutations are expected to be less prone to reversion toward a wild-type, high-affinity phenotype than a single point mutation replacing a critical residue involved in CaM-binding ^28^. As a model system of interacting proteins, we used the antigen-binding fragment from camelidae heavy chain antibodies, or “nanobodies”, V_H_H # 3G9A and V_H_H # 3K1K, here called V_9A_ and V_1K_, respectively, that interact with high affinity (K_D_ ≈ 0.5 nM) with the green fluorescent protein (GFP) as reported by Kirchhofer et al ^29^.

**Fig. 1:**
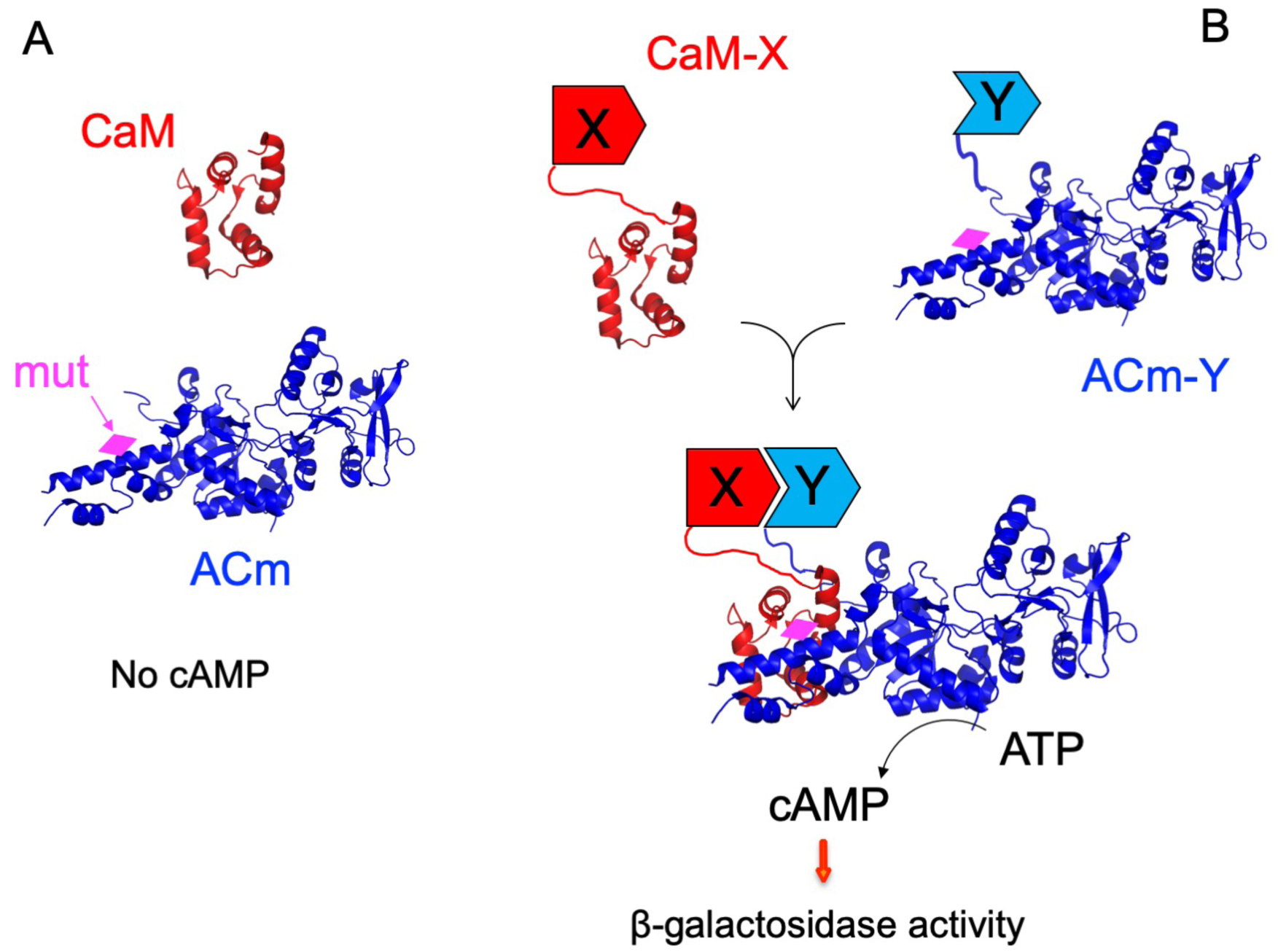
Principle of the exquisitely sensitive adenylate cyclase hybrid (ESACH) system. (A) The catalytic domain of *B. pertussis* adenylate cyclase (blue) is modified (ACm) in its CaM-binding domain (magenta diamond) to decrease its affinity for CaM (red). When expressed at low level in *E. coli ΔcyaA*, CaM cannot activate ACm and there is no cAMP synthesis. (B) When CaM and ACm are fused to two interacting proteins, X and Y, they are brought into close proximity and CaM can activate ACm to produce cAMP (right). Cyclic AMP then binds to the catabolite gene activator protein, CAP, and the cAMP/CAP complex can stimulate the transcription of the catabolite genes, such as the lactose operon or the maltose regulon. The figure was generated from the crystal structure of the catalytic domain of CyaA complexed with the Cterminal domain of CaM (accession number 1zot) ^68^ and visualized in PyMOL (Schrodinger LLC, USA).

### *In vivo* detection of active hybrid AC/CaM complexes

Different expression systems were explored in order to express AC in *E. coli* at the minimal possible level yet enough to confer a selectable Cya*+* phenotype to an *E. coli Δcya* strain. Among them, we selected an expression vector (pAC0) derived from the low-copy plasmid pACYC184, in which all transcriptional and translational control sequences upstream of the AC open reading frame (residues 1 to 399 from *B. pertussis* CyaA) were deleted (Fig. 2, Table S1 and Annex 1). Expression of the wild-type AC as a fusion with GFP from this vector (pAC0-Gfp) was able to restore a Cya*+* phenotype -as detected by the formation of blue colonies on LB X-gal plates (Fig. S1), β-galactosidase assays in liquid cultures or measurements of cAMP on total bacterial extract (Table 1) -to the *E. coli Δcya* strain DHM1, provided the host cells also harbored a compatible plasmid expressing CaM (pTCam, a pUC derived vector with a CaM gene under control of a thermosensitive λ promoter ^30^).

**Fig. 2:**
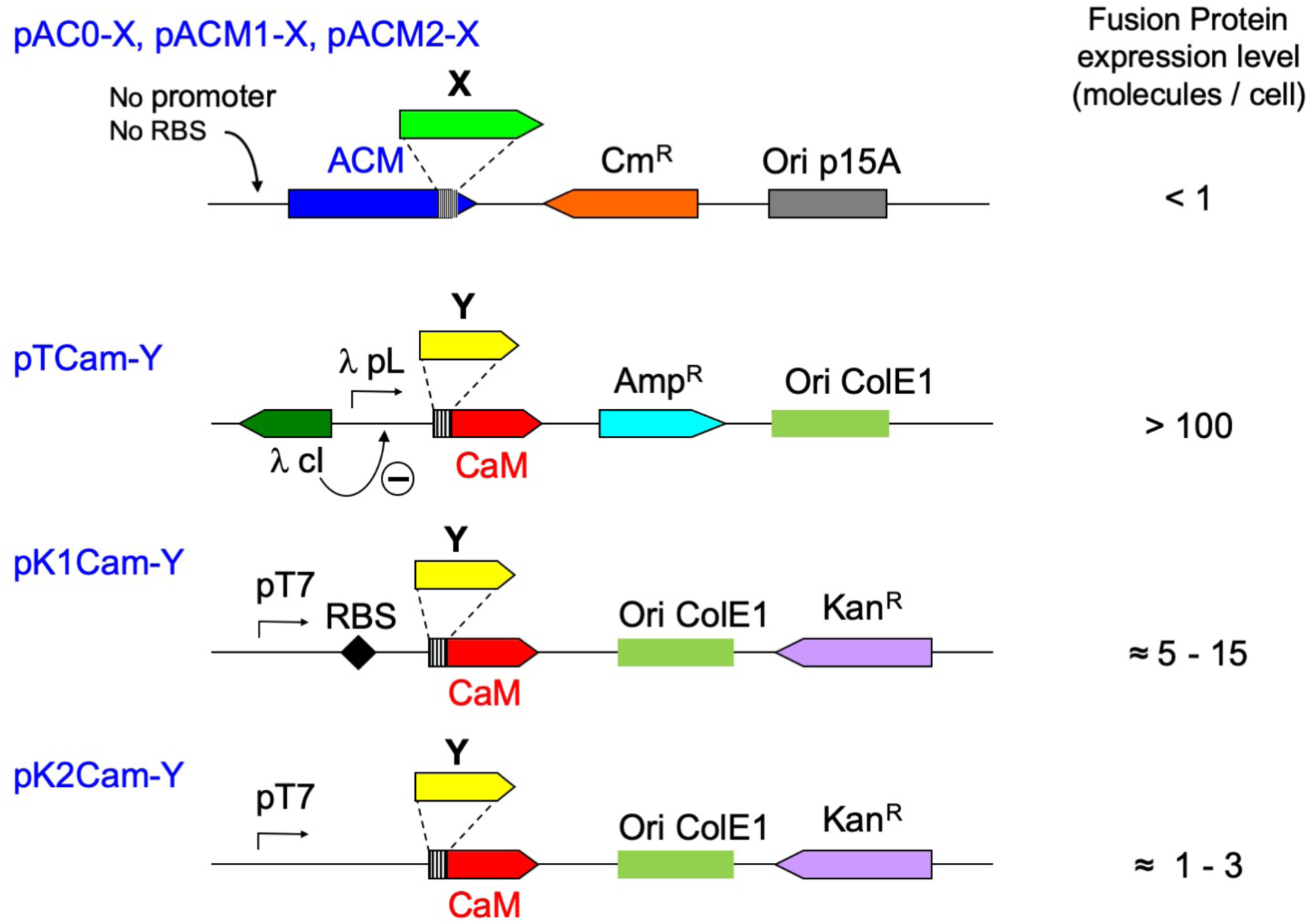
Schematic representation of ESACH plasmids. The colored boxes represent the ORFs of different genes, with the arrows indicating the direction of transcription/translation. The hatched boxes correspond to the multicloning site sequences (MCS) fused to the Cter of ACM or N-ter of CaM. X and Y are the protein of interest fused to ACM and CaM. The origins of replication of the plasmids are indicated by shaded boxes. λcI corresponds to the thermosensitive repressor cI^857^ that strongly represses the λ promoter pL at low temperature (30°C or below), pT7 to the T7 promoter (note that DHM1 does not express any T7 polymerase) and RBS to the ribosome binding site. For each plasmid, the relative expression level of the ACM or CaM fusion proteins, expressed as number of molecules per bacterial cell, and estimated by western blot analysis, is given on the right.

**Table 1:**
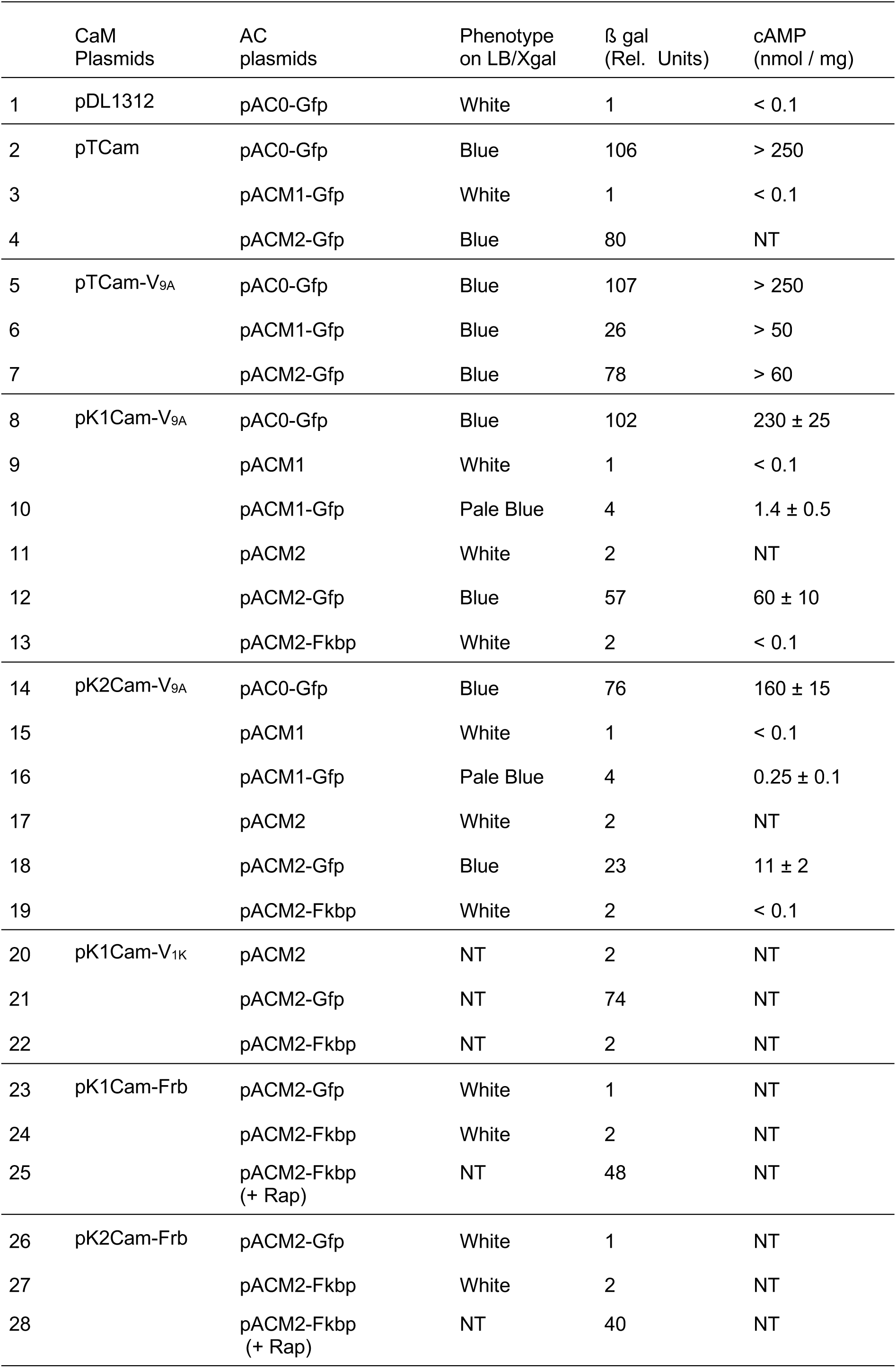
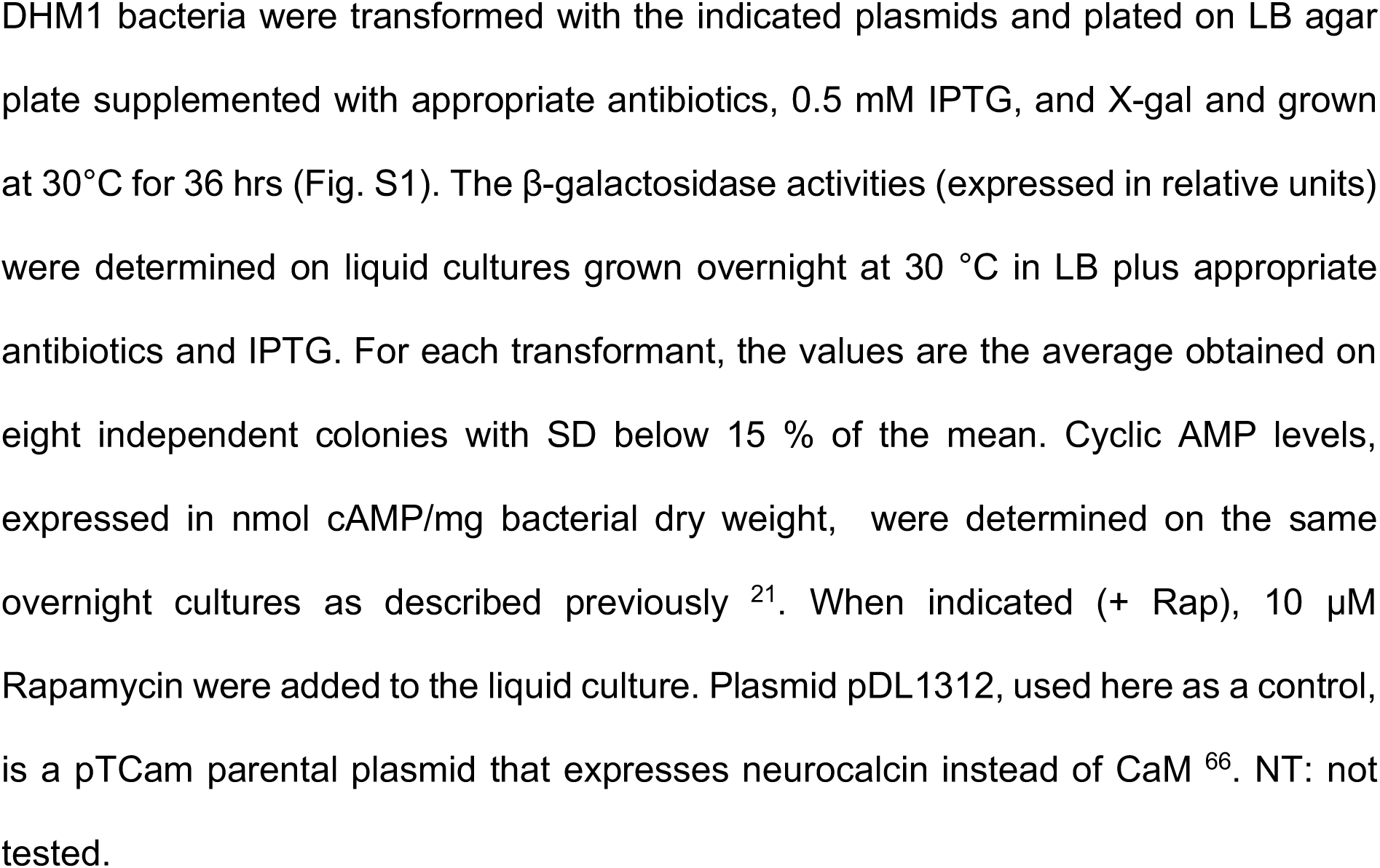
*In vivo* complementation between AC and CaM fusions.

When the ACM1 variant, similarly fused to GFP (encoded by plasmid pACM1-Gfp) was co-expressed with CaM in DHM1, it failed to restore a Cya^+^ phenotype. However, when the pACM1-Gfp plasmid was co-transformed with a pDLTCaM41 derivative, pTCam-V_9A_ (Fig. 2), that expresses CaM as a fusion with the V_9A_ nanobody, the co-transformants exhibited a clear Cya^+^ phenotype, although the cAMP and β-galactosidase expression levels were much lower than in cells co-expressing CaM and the wild-type AC-GFP (Table 1). We concluded that CaM-V_9A_ but not CaM alone, could activate *in vivo* the ACM1-GFP variant as a result of the specific interaction between the V_9A_ and the GFP moieties.

The second AC variant, ACM2, similarly expressed as a GFP fusion (from plasmid pACM2-GFP) also conferred a robust Cya^+^ phenotype to DHM1 when co-transformed with pTCam-V_9A,_ as anticipated, but also and unexpectedly, when co-transformed with pTCaM (Table 1). We hypothesized that the CaM protein level -due to the residual transcription from the cI^857^-repressed λ promoter (Fig. 2) - was high enough to spontaneously activate the ACM2 variant that has a higher affinity for CaM than ACM1 (^27^ and Fig. 3). We therefore tested alternative expression systems in order to reduce the level of expression of CaM fusions. We constructed two plasmids, pK1Cam-V_9A_ and pK2Cam-V_9A_, in which the CaM-V_9A_ fusion was expressed under the control of a T7 promoter with or without an RBS sequence, respectively (Fig. 2, Table S1 and Annex 1). As the DHM1 cells have no T7 polymerase, the CaM fusion was expected to be expressed at very low level. These plasmids, when co-transformed with pAC0-GFP or with pACM2-GFP, conferred a robust Cya^+^ phenotype to DHM1 cells although the cAMP and β-galactosidase levels were lower with the latter as compared to the former (Table 1). Only a barely detectable Cya^+^ phenotype was noticed when they were co-transformed with pACM1-Gfp, likely because of the lower specific activity of the ACM1 variant as compared to wild-type AC or ACM2 (^27^ and see Fig.3). More importantly, a Cya^-^ phenotype was obtained upon co-transformation of pK1Cam-V_9A_ or pK2Cam-V_9A_, with pACM2 that expresses ACM2 alone (Table 1). Hence, the expression levels of the CaM-V_9A_ fusion obtained with these new plasmids now appeared to be appropriate to allow the selective activation of the ACM2-GFP fusion but not of ACM2 alone. We similarly tested another camelidae V_H_H (V_H_H # 3K1K), here noted V_1K 29_, that also exhibits a high affinity for GFP: the CaM-V_1K_ fusion was also found to selectively activate *in vivo* the ACM2-GFP fusion but not ACM2 (Table 1).

**Fig. 3:**
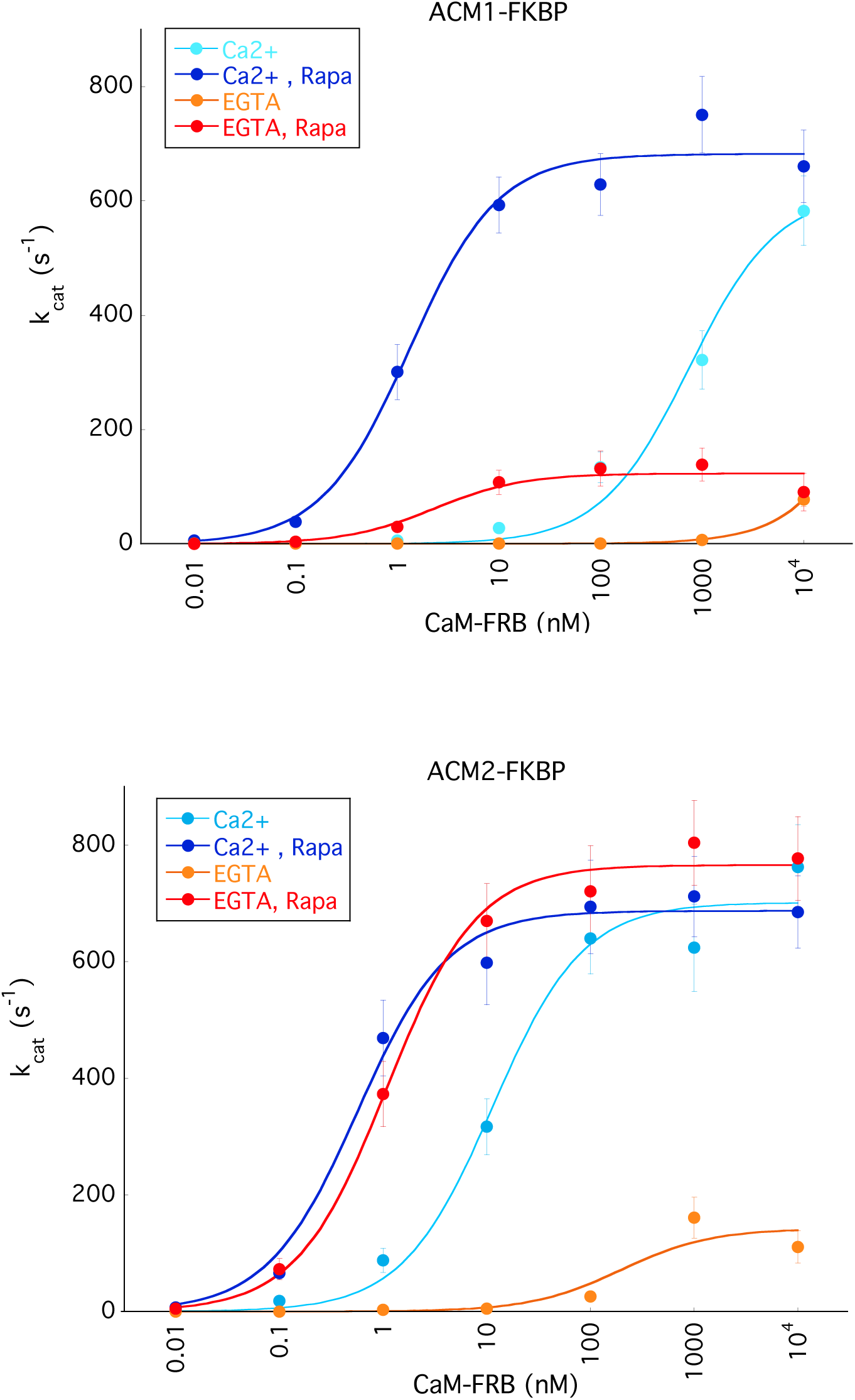
*In vitro* assays of the rapamycin-induced activation of ACm-FKBP by CaM-FRB. Purified ACM1-FKBP (upper panel) and ACM2-FKBP (lower panel) (both at 0.5 nM) were pre-incubated for 10 min at 30°C with the indicated concentrations of CaM-FRB in a reaction medium containing either 0.1 mM calcium (Ca^2+^) or 0.1 mM calcium and 1 mM EGTA (EGTA, i.e. no free calcium) and in the presence of 5 μM Rapamycin (Rapa) when indicated. Reactions were initiated by adding 2 mM ATP and carried out for 20 min. Cyclic AMP was then measured spectrophotometrically as described in Material and Methods. The AC activities (expressed in k_cat_ = mol of cAMP produced per mol of enzyme per sec) at the indicated CaM-FRB concentrations were fitted to equation: A = (A_Max_ x [CaM]) / (K_D_ + [CaM]) with A_Max_ = maximal activities at saturating CaM-FRB and K_D_ = the CaM-FRB concentration at half-maximal activation. The deduced A_Max_ and K_D_ values are listed in Table S2 (Supplementary Material)

The ESACH system was further characterized by analyzing the rapamycin-induced interaction between the FK506-binding protein, FKBP and the FKBP-rapamycin binding domain, FRB, that binds with high affinity to FKBP in the presence of rapamycin ^31^ . Plasmids expressing ACM2 fused to FKBP, pACM2-Fkbp, and CaM fused to FRB, pK1Cam-Frb and pK2Cam-Frb (Table S1 and Annex 1), were constructed and tested in complementation assays in DHM1 as above. As shown in Table 1, the CaM-FRB fusion, produced by the pK1Cam-Frb or pK2Cam-Frb plasmids, was able to specifically activate *in vivo* the ACM2-FKBP only when the cells were grown in the presence of rapamycin. Furthermore, the CaM-FRB fusion did not activate the ACM2-GFP hybrid and conversely the CaM-V_9A_ or CaM-V_1K_ fusions did not activate the ACM2-FKBP hybrid (Table 1). Altogether, these data indicate that the CaM fusions produced by pK1Cam or pK2Cam plasmids could activate *in vivo* the ACM2 hybrids in a highly selective manner, dictated by the specific association between the protein modules appended to CaM and ACM2.

### *In vitro* assays of the interaction-mediated activation of modified ACs

To corroborate the above results at the molecular level, we characterized *in vitro* the interaction-mediated activation of modified AC-fusions by cognate CaM-fusions. The ACM1 and ACM2-FKBP fusions as well as the CaM-FRB fusion were overexpressed in *E. coli* and purified to near homogeneity (see Material and Methods and Fig. S2). The activity of the AC fusions in the presence of increasing concentrations of CaM-FRB was then assayed *in vitro* ^32^ either in the presence or in the absence of calcium or rapamycin. As shown in Fig. 3, in the presence of calcium and absence of rapamycin (ie when FKBP and FRB do not associate), the CaM-FRB concentrations at half-maximal activation (≈ K_D_) were found to be ∼ 750 nM for ACM1-FKBP and ∼ 10-15 nM for ACM2-FKBP (Table S2), in agreement with earlier measurements of the activation of ACM1 or ACM2 by native CaM ^27^. Upon addition of rapamycin, both AC fusions were half-maximally activated at about 1-2 nM CaM-FRB, a concentration close to the FKBP/rapamycin/FRB equilibrium constant ^31^. Most notably, in the absence of calcium, that is, in conditions prevailing in the bacterial cytosol, and without rapapmycin, ACM1-FKBP was barely activated by CaM-FRB, while ACM2-FKBP was partially activated at CaM-FRB concentrations above 100 nM. This can explain why a Cya^+^ phenotype was observed when DHM1 was co-transformed with pACM2-GFP and pTCam as this plasmid likely expressed high CaM levels in bacteria. In the absence of calcium and in the presence of rapamycin ACM2-FKBP was fully activated by CaM-FRB, with a half-stimulating concentration of about 1 nM. Interestingly, in the same conditions, ACM1-FKBP was also half-maximally activated at about 2.5 nM CaM-FRB, but its maximal activity reached only about 15% of that found in the presence of calcium. We hypothesize that in the presence of rapamycin, ACM1-FKBP/rapamycin is fully saturated by CaM-FRB at concentrations above 10 nM, but, in this complex, the Ca^2+^-free CaM moiety can only transiently associate (e.g. about 15 % of the time) with the ACM1 moiety to produce an active enzyme, as a result of the strongly destabilizing mutation introduced in ACM1. All together, these experiments directly corroborated the stringent interaction-mediated activation of the ACM fusions observed in bacteria. They also explained the lower efficiency of complementation observed *in vivo* with the ACM1 variant as compared with ACM2.

### Detection of interaction between membrane associated proteins

We then explored the capacity of the ESACH system to detect interactions between membrane-associated and/or periplasmic proteins. For this, we constructed plasmids (Fig. 4A, Table S1 and Annex 1) expressing ACM2 or CaM-V_9A_ fused to a trans-membrane (TM) segment (the first TM from the *E. coli* OppB oligopeptide transporter subunit ^24^) followed by the leucine zipper dimerization domain of GAL4 (Zip). The resulting fusions, ACM2-TM-zip and CaM-V_9A_-TM-zip, should insert into the membrane with their Zip motif localized in the periplasm (see Fig. 4B). As controls, ACM2 and CaM-V_9A_ were also expressed as fusions to the leucine zipper motif without the TM domain (*i.e.* localized in the cytosolic compartment). As shown in Table 2, the membrane-associated ACM2-TM-Zip and CaM-V_9A_-TM-Zip hybrids efficiently associated through their leucine zipper motifs localized in the periplasm. The interaction signal was similar to that detected between the hybrid proteins ACM2-Zip and CaM-V_9A_-Zip that have a cytosolic leucine zipper (Fig. 4B). Importantly, ACM2-TM-Zip did not interact with CaM-V_9A_-Zip nor ACM2-Zip with CaM-V_9A_-TM-Zip (Table 2). This indicates that the leucine zipper motif of the TM-Zip constructs was properly addressed to the periplasm by the OppB TM domain. In addition, both the membrane-anchored CaM-V_9A_-TM-Zip and the cytosolic CaM-V_9A_-Zip could efficiently interact *via* their cytoplasmic-localized V_9A_ module (Fig. 4B), with ACM2-GFP (but not with ACM2-FKBP as expected) (Table 2). These data thus indicate that the ESACH system can efficiently report interactions involving integral membrane proteins or interactions between an integral membrane protein and a cytosolic protein. In addition, they show that CaM can be dually tagged at both its N and C termini to yield fusions that can specifically activate two distinct ACM-fusions.

**Fig. 4:**
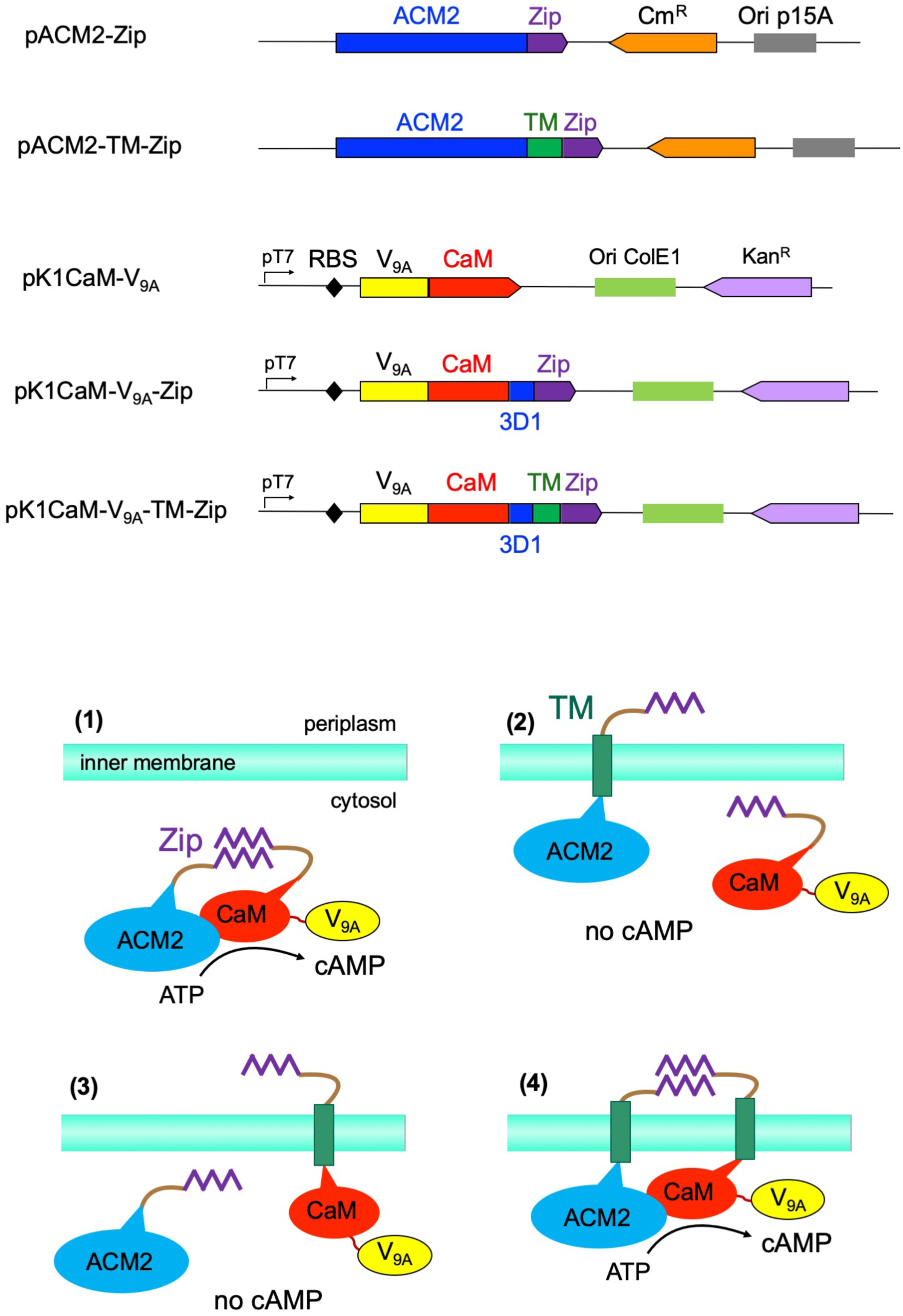
Detection of interactions between membrane-associated proteins with the ESACH system. Upper panel: Schematic representation of plasmids expressing cytosolic or membrane-associated ACM2 & CaM hybrid proteins. The colored boxes represent the ORFs of the different genes, with the arrow indicating the direction of transcription/translation. ACM2 is in blue, the leucine zipper motif (Zip) in violet, the OppB transmembrane segment (TM) in green, the V_9A_ nanobody is in yellow, and CaM is in red. The 3D1 blue rectangle in CaM plasmids correspond to the very C-terminal segment of AC containing the 3D1 epitope that was inserted during cloning of the leucine zipper motif. The chloramphenicol (Cm^R^) and kanamycin (Kan^R^) resistant markers as well as the p15A and ColE1 origins of replication are indicated by colored rectangles. Lower panel: Schematic representation of the expected topology of the different ACM2 and CaM hybrid proteins relative to the inner membrane (light green bar); (1) : ACM2-Zip / V_9A_-CaM-Zip; (2) : ACM2-TM-Zip / V_9A_-CaM-Zip; (3) : ACM2-Zip / V_9A_-CaM-TM-Zip; (4) : ACM2-TM-Zip / V_9A_-CaM-TM-Zip.

**Table 2:** Interaction assays of membrane associated fusion proteins.

|  | pK1Cam-V<sub>9A</sub> | pK1Cam-V<sub>9A</sub>-Zip | pK1Cam-V<sub>9A</sub>-TM-Zip |
| --- | --- | --- | --- |
| pACM2-Zip | 2 | 42 | 2 |
| pACM2-TM-Zip | 2 | 2 | 35 |
| pACM2-Gfp | 47 | 17 | 44 |
| pACM2-Fkbp | 2 | 2 | 2 |

*In vivo* complementation assays between various AC and CaM fusions. DHM1 bacteria were transformed with the indicated plasmids and plated on LB agar supplemented with appropriate antibiotics, IPTG, and X-gal and grown at 30°C for 36 hrs. The β-galactosidase activities (expressed in relative units) were determined on liquid cultures grown overnight at 30 °C in LB plus appropriate antibiotics and IPTG. For each transformant, the values are the average obtained on six to eight independent colonies (in all cases SD were below 20 % of the mean values).

### Expression levels of hybrid proteins

We next quantified the level of expression of the ACM and CaM hybrid proteins in the bacterial cells by western blot (WB). The ACM proteins could be specifically detected with a monoclonal antibody (Mab) 3D1 that recognizes an epitope located between residues 373 and 400 of AC ^33^. To quantify the amount of ACM2-FKBP protein produced per bacterial cell, the hybrid protein was over-expressed in *E. coli* (Material and Methods) and purified to near homogeneity (see Fig. S2) to serve as a standard. As shown in Fig. 5A, the 3D1 Mab was able to detect down to ≈ 0.01 ng of ACM2-FKBP fusion, which correspond to about 10^8^ ACM2-FKBP molecules (molecular weight of ≈ 73kDa). A bacterial extract corresponding to 1 ml of culture at OD_600_ = 1, i.e. about 10^9^ bacteria), of DHM1 cells expressing ACM2-FKBP and CaM-FRB (grown in the presence of rapamycin) was probed in parallel on the same WB. As shown in Fig. 5A, a very faint signal (in the range of that found for 0.01 ng of ACM2-FKBP fusion) could be detected by WB in this extract. This indicates that the bacteria harboring pACM2-FKBP expressed, on average, less than one ACM2-FKBP molecule per cell (i.e. below 0.1 ng of protein fusion per 10^9^ cells).

**Fig. 5:**
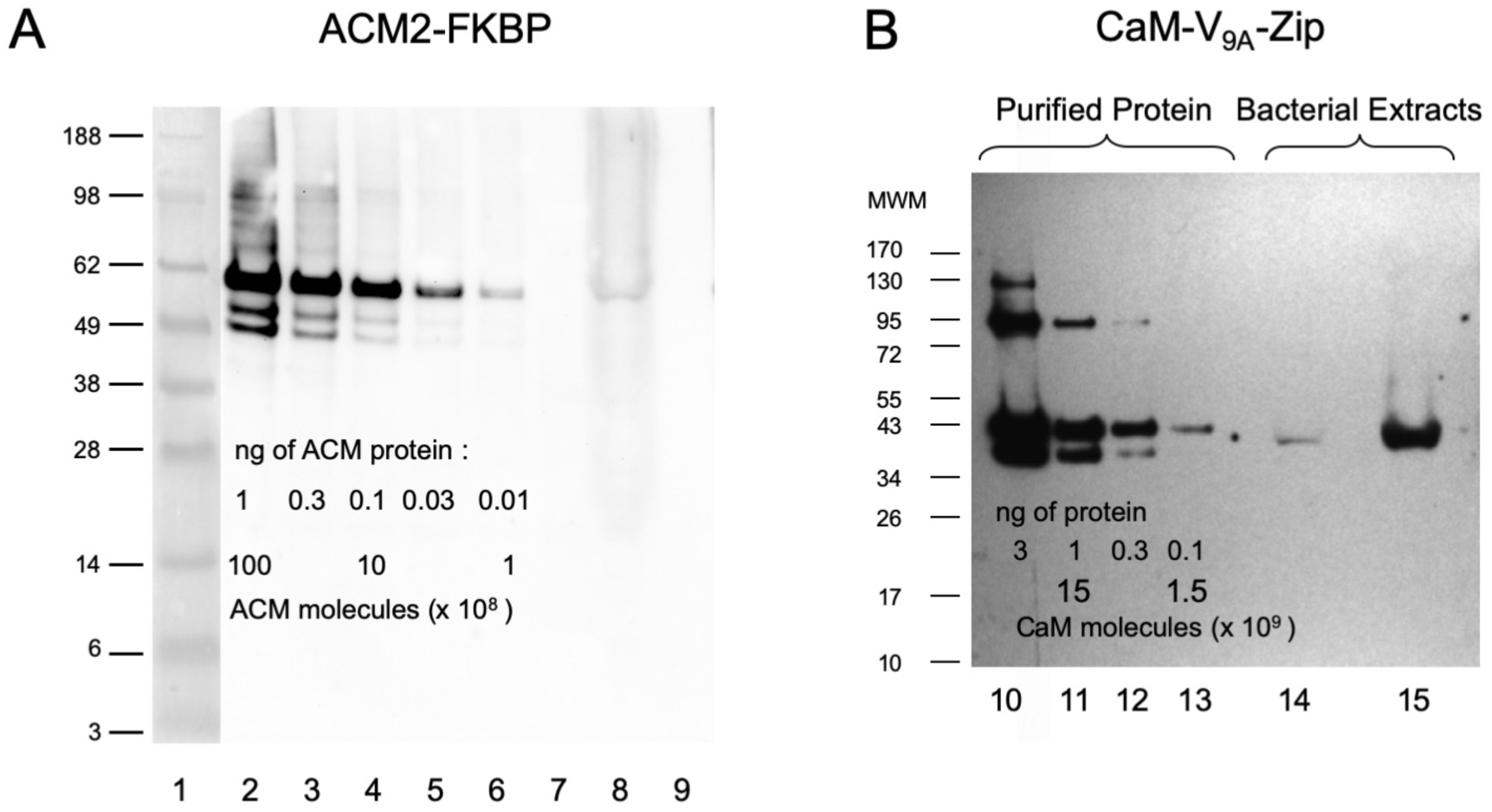
Expression levels of ACM and CaM hybrid proteins *in vivo*. A : Western blot analysis of the expression of the ACM2-FKBP hybrid protein in DHM1. Line 1: Molecular Weight Markers (size in kDa indicated on the left side). Lines 2 - 6 : 1, 0.3, 0.1, 0.03 and 0.01 ng respectively of the purified ACM2-FKBP hybrid protein (molecular weight of ≈ 56.5 kDa; 0.01 ng of ACM2-FKBP fusion correspond to ≈ 10^8^ protein molecules) were separated by electrophoresis, electro-transferred to nitrocellulose and detected with 3D1 monoclonal antibody. Line 7: no protein. Line 8 : Total bacterial extracts (corresponding to ≈ 10^9^ bacteria i.e. 1 ml of cell culture at OD_600_ = 1) from a mid-log phase culture (in the presence of rapamycin) of DHM1/pACM2-Fkbp/pK1Cam-Frb were probed in parallel by Western blot; Line 9: no protein. B : Western blot analysis of the expression of the CaM fusion proteins in DHM1. Lines 10-13 : 3, 1, 0.3, and 0.1 ng, respectively of the purified CaM-V_9A_-Zip protein (molecular weight of ≈ 43 kDa; 0.1 ng of CaM-V_9A_-Zip fusion correspond to ≈ 1.4 x 10^9^ molecules) were separated by electrophoresis, electro-transferred to nitrocellulose and detected with the 3D1 monoclonal antibody. Lines 14 and 15 : Total bacterial extracts (corresponding to ≈ 10^9^ bacteria; i.e. 1 OD_600_) from mid-log phase cultures of DHM1 cells harboring the indicated plasmids were probed in parallel by Western blot; line 14: pACM2-Zip/pK2Cam-V_9A_-Zip; line 15: pACM2-Zip/pK1Cam-V_9A_-Zip.

We analyzed similarly different extracts of bacterial cultures harboring various combinations of plasmids, e.g. pAC0-Gfp, pACM1-Gfp, or other pACM2 plasmids derivatives, (Fig. S3A), and in all cases the WB signal intensities (actually the absence of signal) confirm that less than 1 molecule of ACM fusions were expressed per cell with the different pACM1 and pACM2 hybrid proteins. In control experiments (Fig. S3B) we checked that the very low level of detection of ACM fusions in the bacterial extracts was not due to the presence of the large amount of bacterial proteins, as sub-ng amounts of purified ACM2-GFP spiked in the extract could be readily detected by WB with the 3D1 Mab. All together, these WB experiments indicate that the expression level of AC hybrid achieved with the ESACH vectors is on average **<u>below</u>** one molecule per bacterial cell.

The expression level of the CaM fusions produced from the pK1Cam and pK2Cam plasmids were similarly determined by WB using the same 3D1 Mab, as the CaM-V_9A_-Zip fusion also contains the AC-derived 3D1 epitope (initial attempts to quantify the CaM-fusion with a FLAG-tagged CaM turned out to be not sensitive enough, data not shown). The CaM-V_9A_-Zip fusion was overexpressed and purified to serve as standard (see Material and Methods). About 0.1 ng of the purified CaM-V_9A_-Zip fusion could be detected by Mab 3D1 in WB (Fig. 5B) corresponding to ≈ 14 x 10^8^ protein molecules (molecular weight of ≈ 42 kDa). A similar signal was detected in total extracts of 10^9^ DHM1 cells harboring pK2Cam-V_9A_-Zip and pACM2-Zip indicating a ratio of about 1 -3 CaM fusions per cell. A higher signal was found in extracts of DHM1 cells harboring pK1Cam-V_9A_-Zip and pACM2-Zip, corresponding to about 5 - 15 CaM hybrids per bacteria (note that given the mean volume of the *E. coli* cytosol, 1 molecule per cell corresponds roughly to a concentration of 1 nM) (Fig. 5B)

Altogether, these results highlight the exquisite sensitivity of the AC/CaM/cAMP signaling cascade that could reliably detect in *E. coli,* interactions between hybrid proteins expressed at a minimal level of few molecules per cell in the case of the CaM fusions, and more strikingly, at less than one molecule per cell, on average, in the case of the ACM fusions.

### Characterization of interactions involving toxic proteins

To further establish that the ACM fusions are expressed *in vivo* at an extremely low level, we explored the interaction of the toxic enzyme barnase, a ribonuclease secreted by the bacterium *Bacillus amyloliquefaciens,* with barstar a specific inhibitor that binds with high affinity to barnase and blocks its RNAse activity. Barnase is lethal to bacterial cells when expressed without its inhibitor Barstar ^34^ ^35^. Synthetic Barnase and Barstar genes were cloned into the pACM2 and pK1Cam plasmids respectively. DHM1 cells cotransformed with the two resulting plasmids, pACM2-Barnase and pK1Cam-Barstar, exhibited a strong Cya*^+^* phenotype, while control co-transformations with various pACM2 and pK1Cam derivatives demonstrated the selectivity of interaction between the Barnase and Barstar modules (Fig. 6). Noticeably, the pACM2-Barnase plasmid could be transformed into DHM1 cells that did not expressed any Barstar fusions (e. g. harboring pK1Cam-Frb, pK1Cam-V_9A_ or without additional plasmid) and the transformed cells did not exhibit any detectable growth problem (Fig. 6). This confirms that the ACM2-Barnase hybrid protein was expressed at a level low enough not to affect the bacterial physiology, yet sufficient to allow an efficient detection of the Barnase-Barstar interaction. Hence the ESACH system may be used to characterize the interaction properties of highly toxic proteins in bacteria, including the wide variety of toxin-antitoxin systems that are present in these organisms ^36^.

**Fig. 6:**
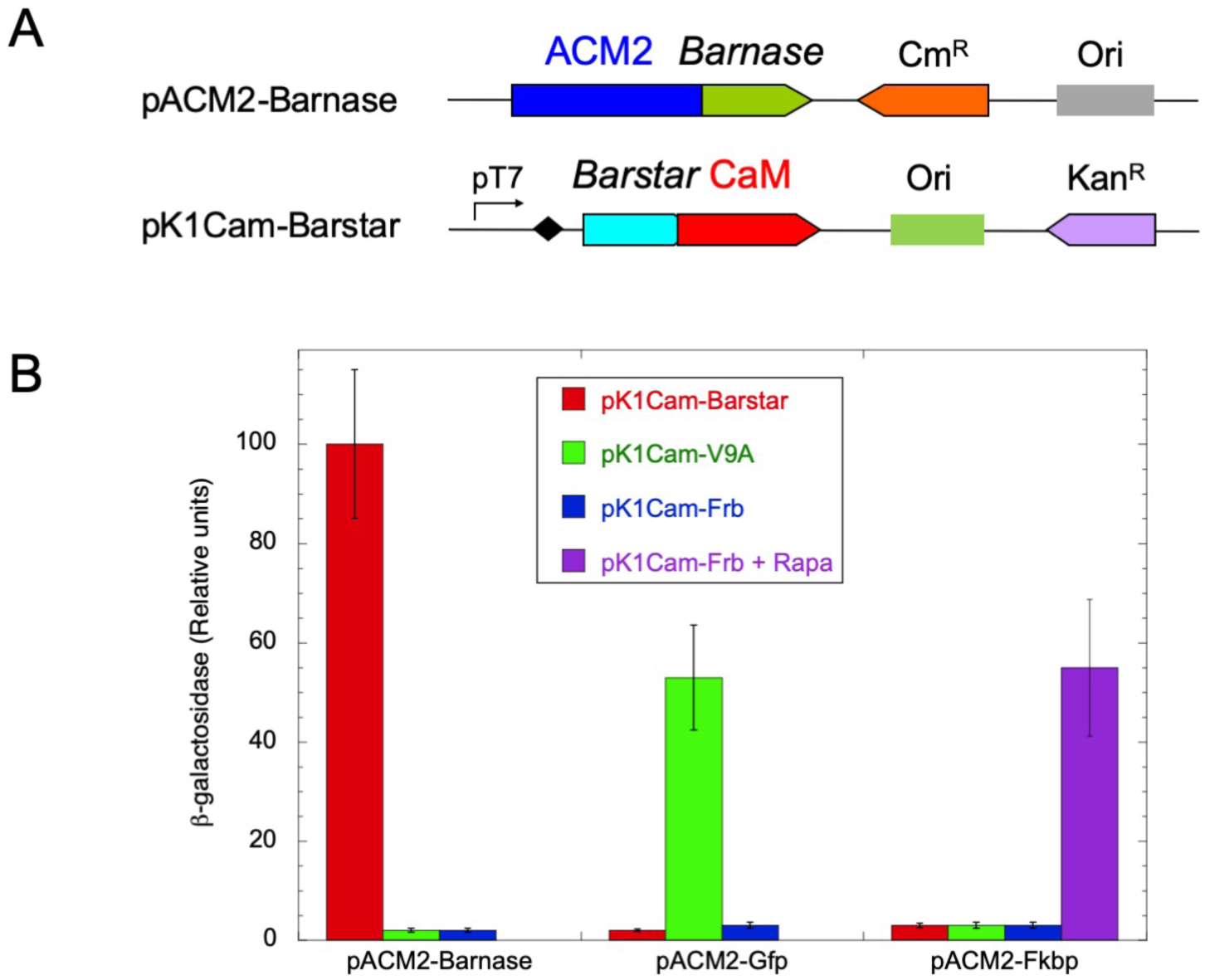
*In vivo* detection of Barnase/Barstar interaction with ESACH system. A: Schematic representation of pACM2-Barnase & pK1Cam-Barstar plasmids. The colored boxes represent the ORFs of different genes, with the arrow indicating the direction of transcription/translation. B: β-galactosidase activities (relative units) in DHM1 cells transformed with the indicated plasmids and grown at 30 °C in LB medium supplemented with IPTG and antibiotics, and 5 μM rapamycin when indicated (+ Rapa).

### CaM variants as AC activating partners *in vivo*

We also explored an alternative design for the ESACH system using as complementary modules, a wild-type AC enzyme and a modified CaM with decreased affinity for AC. We first tested a previously characterized CaM variant, CaM_VU8,_ in which three glutamic acid residues at position 82, 83, and 84 of CaM are substituted by lysines: CaM_VU8_ displays ≈ 1000-fold lower affinity for wild-type AC compared to native CaM ^37^. CaM_VU8_ expressed from the pCam_VU8_ plasmid (see Material & methods, Table S1 and Annex 1).) was unable to activate *in vivo* the wild-type AC or the AC-GFP fusion (Table 3). When CaM_VU8_ was expressed as a fusion with the V_1K_ nanobody, it efficiently activated the AC-GFP fusion but not AC alone. Furthermore, CaM_VU8_-V_1K_ could not activate the ACM2-GFP fusion, likely because of the too-low affinity of the mutant CaM_VU8_ for the ACM2 variant.

**Table 3:** CaM variants as activators of AC in ESACH interaction assays.

|  | pCam<sub>VU8</sub> | pCam<sub>VU8-V1K</sub> | pCam<sub>Cter-V1K</sub> |
| --- | --- | --- | --- |
| pAC0 | 6 | 5 | 29 (± 5) |
| pAC0-Gfp | 7 | 111 (± 10) | 187 (± 13) |
| pACM2 | 5 | 5 | 4 |
| pACM2-Gfp | 6 | 5 | 141 (± 15) |
| pACM2-Fkbp | 6 | 5 | 4 |

*In vivo* complementation assays between various AC and CaM fusions. DHM1 bacteria were transformed with the indicated plasmids and plated on LB agar supplemented with appropriate antibiotics, IPTG, and X-gal and grown at 30°C for 36 hrs in the presence of 0.5 mM IPTG plus appropriate antibiotics. The β-galactosidase activities (expressed in relative units) were determined on liquid cultures grown overnight at 30 °C in LB plus appropriate antibiotics and IPTG. For each transformant, the values are the average obtained on eight independent colonies (SD when not indicated were below 20 % of the mean).

We then tested the C-terminal moiety of CaM (i.e. residues 77 to 148), CaM_Cter_, as a potential activating partner of AC in ESACH assay. CaM_Cter_ has been shown to fully activate AC with a ≈ 10 fold lower affinity than native CaM ^38^. As shown in Table 3, CaM_Cter_ expressed as a fusion with the V_1K_ nanobody efficiently activated the AC-GFP fusion. Yet it also partly stimulated the AC alone, likely because of its relatively high affinity for the enzyme. More interestingly, the CaM_Cter_-V_1K_ fusion efficiently activated the ACM2-GFP fusion but not ACM2 or the ACM2-FKBP fusion. Therefore, the CaM_Cter_ fragment, which is only 72 amino-acid long, can be used in combination with the ACM2 variant for high sensitive detection of interactions *in vivo* in ESACH assay.

### Screening of nanobody-antigen interactions in bacteria

We showed above that the ESACH system can detect of the specific association of the two different nanobodies V_9A_ and V_1K_ with their target antigen GFP ^29^. To examine whether it could also be applied for direct *in vivo* selection of nanobodies (or other binding modules ^39^ ^40^, we randomly PCR-mutagenized the V_9A_ nanobody at amino acid residue 107, located at the interface with GFP in the crystal structure ^29^, and screened for variants unable to bind to GFP (see Fig. S4). One such variant, Y_107N_, harboring a change of Tyr107 into Asn, abolished the interaction with GFP as revealed by the Cya*^-^* phenotype of DHM1/pACM2-Gfp/pK2Cam-Y_107N_ transformants. The plasmid pK2Cam-Y_107N_ expressing the V_9A_-Y_107N_ variant was chosen for a second round of mutagenesis in which the three amino acid positions 105, 106 or 107 of the V_9A_ were randomly mutagenized (Fig. S4). The mutagenized plasmid pool was co-transformed into DHM1/pACM2-Gfp and plated on a selective medium ^41^ ^42^, a minimal medium containing maltose as a unique carbon source: as the maltose regulon is under a stringent cAMP/CAP control, only Cya*^+^* bacteria can grow on this medium (Xgal and IPTG were also added to better visualize the Cya*^+^*colonies). We randomly chose a number of Cya*^+^* colonies that grew on this selective medium and sequenced their pK2Cam-V_9A_ plasmids. As shown in Table S3, all fusions contained a tyrosine (or in one case a phenylalanine) at position 107, and a glycine residue at position 106, indicating that these residues were mostly critical for interaction of V_9A_ with GFP. In contrast, a variety of residues could be found in Cya^+^ clones at codon 105, indicating that this position was less important for interaction, in agreement with the 3D structure of the GFP/V_9A_ complex ^29^. As a control, the mutagenized plasmid pool was also co-transformed into DHM1/pAC2-Fkbp and plated on the same selective medium. No colonies could be detected in these conditions highlighting the stringency of the *in vivo* selection. All together, these experiments indicate that the ESACH system could be used for direct *in vivo* selection of binding proteins.

### Visualization of active hybrid AC/CaM complexes *in vivo*

To further document the counterintuitive observation that cells could express on average less than one ACM/CaM active complex per bacterium and yet display a selectable Cya*^+^* phenotype, we attempted to visualize the complementation between hybrid proteins *in vivo*, on individual bacteria through a fluorescent reporter. For this, the gene coding for the ZsGreen fluorescent protein (Clontech Laboratories) was placed under the transcriptional control of a cAMP/CAP dependent *lac* promoter and inserted into the pK1Cam-Frb plasmid (Fig. 7A). The resulting plasmid pK1Cam-Frb-Zs was co-transformed into DHM1 cells with either pACM2-Fkbp or pACM2-Gfp. The co-transformants displayed a background fluorescence signal when grown in LB medium but were highly fluorescent when grown in LB medium supplemented with cAMP, which can diffuse inside the cells to directly stimulate cAMP-dependent gene transcription (Fig. 7B). As expected, DHM1 cells co-transformed with pK1Cam-Frb-Zs and pACM2-Fkbp also displayed high fluorescence when grown in LB medium supplemented with rapamycin while those co-transformed with pK1Cam-Frb-Zs and pACM2-Gfp remained non-fluorescent (Fig.7B). This indicates that the placZsgreen fluorescent reporter could detect *in vivo* the rapamycin-induced interaction between ACM2-FKBP and FRB-CaM and the resulting activation of AC enzymatic activity. We then imaged the kinetics of rapamycin-induced activation of ACM2-FKBP by CaM-FRB in cells grown in LB medium. As shown in Fig.7C, within the first hours of growth in the presence of rapamycin, only about 20-25 % of the cells became fluorescent. This fraction progressively increased to more than 90 % of total population after an overnight culture. No fluorescent cells were detected when rapamycin was added to DHM1/pACM2-Gfp-GFP/pK1Cam-Frb-Zs. Yet, all these cells became highly fluorescent within 0.5-1 hr after addition of cAMP in the medium (“control” in Fig. 7C). We interpret these results as follows, as illustrated in Fig. 8: in DHM1 cells harboring pACM2-FKBP, the ACM2-FKBP fusion protein is stochastically expressed on average in about 20-25 % of the cells. In these cells, addition of rapamycin triggers the interaction of the hybrid enzyme with the co-expressed FRB-CaM, and these bacteria start to produce cAMP and consequently to express the ZsGreen fluorescent reporter. The other cells, lacking the ACM2-FKBP hybrid, obviously cannot produce cAMP and thus remain non-fluorescent. However, upon prolonged exposure to rapamycin, the progeny of these ZsGreen^-^ cells will progressively became fluorescent as a result of stochastic expression of the ACM2-FKBP that should occur statistically once every 2-3 cell cycles (if present statistically in 20-25 % of the cells). The progeny of the ZsGreen^+^ cells should retain the fluorescence of the mother cell due to the random distribution of the ZsGreen protein between the two daughter cells, as well as that of the cAMP/CAP complex, which could thus trigger transient *de novo* ZsGreen expression in daughter cells. In addition, one of the two daughter cells should inherit the active ACM/CaM complex to continue synthetizing high amounts of cAMP. All together these data support the view that the exquisite sensitivity of the ESACH system could allow bacterial colonies to be selected for their Cya^+^ phenotype, even though at any given time only a fraction of all bacteria may harbor an active ACM/CaM hybrid complex.

**Fig. 7:**
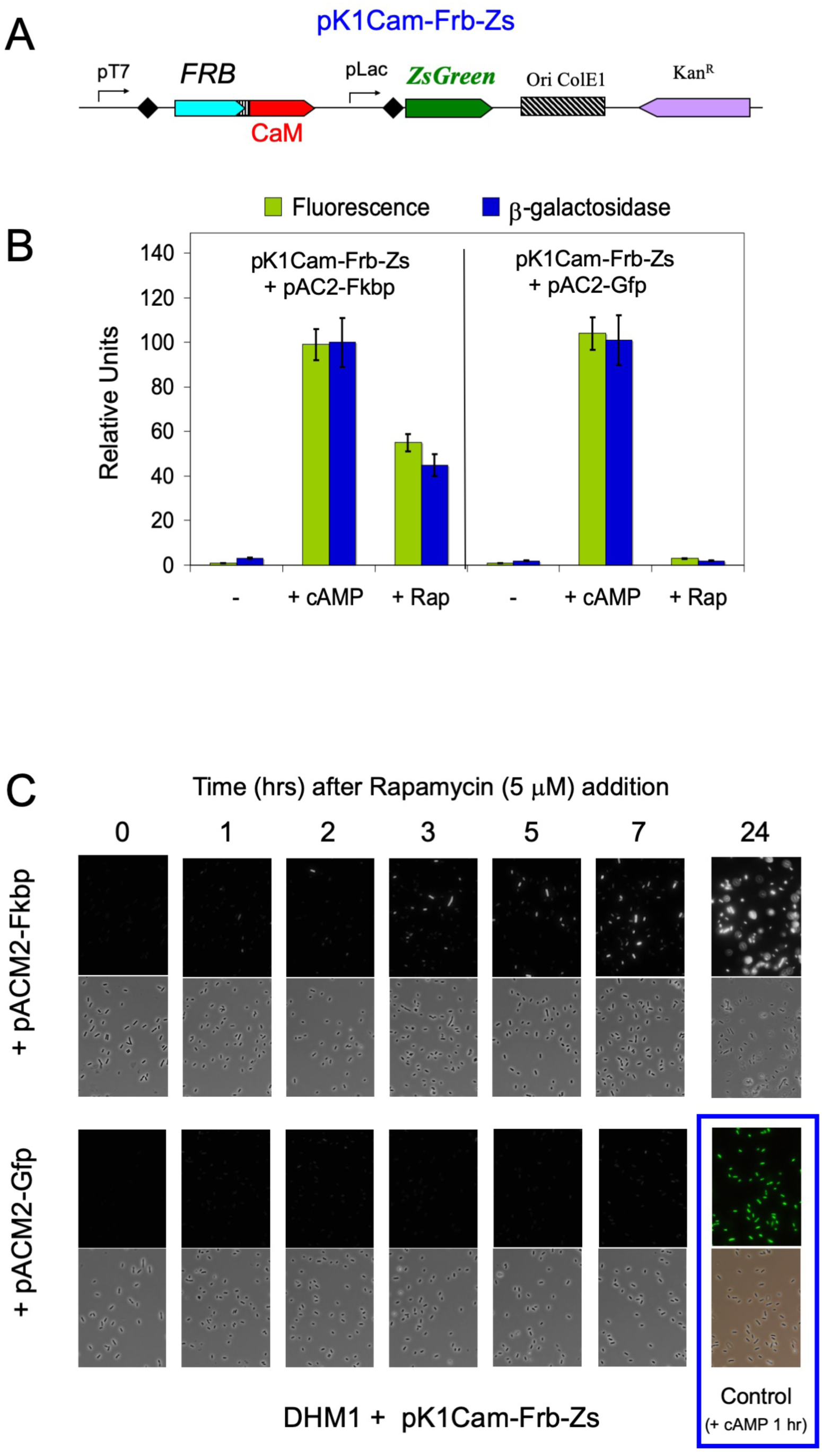
*In vivo* detection of rapamycin-induced ACM2-FKBP/CaM-FRB interaction. A: Schematic representation of pK1Cam-Frb-Zs plasmid. The colored boxes represent the ORFs of different genes, with the arrow indicating the direction of transcription/translation. pLac correspond to the lac promoter driving the expression of the ZsGreen fluorescent protein. B: DHM1 cells, harboring either pK1Cam-Frb-Zs and pACM2-Fkbp (left) or pK1Cam-Frb-Zs and pACM2-Gfp (right), were grown overnight at 30 °C in LB medium containing appropriate antibiotics, IPTG and when indicated, either 2 mM cAMP or 5 μM Rapamycin (Rap). ZsGreen fluorescence (green bars) and β-galactosidase activities (blue bars) were measured (on 8 distinct colonies for each condition) as described in Material and Methods. C: DHM1 cells, harboring pK1Cam-Frb-Zs and either pACM2-Fkbp or pACM2-Gfp, were grown overnight at 30 °C in LB medium containing appropriate antibiotics then diluted 1:100 in LB medium containing antibiotics and IPTG and further incubated at 30°C until early exponential phase. Rapamycin (5 μM) was added at time 0 and cells were imaged at the indicated time on a Nikon epi-fluorescence microscope. Bottom right : control experiment in which DHM1/pK1Cam-Frb-Zs/pACM2-Gfp cells were imaged 1 hr after addition of 2 mM cAMP instead of rapamycin.

**Fig. 8:**
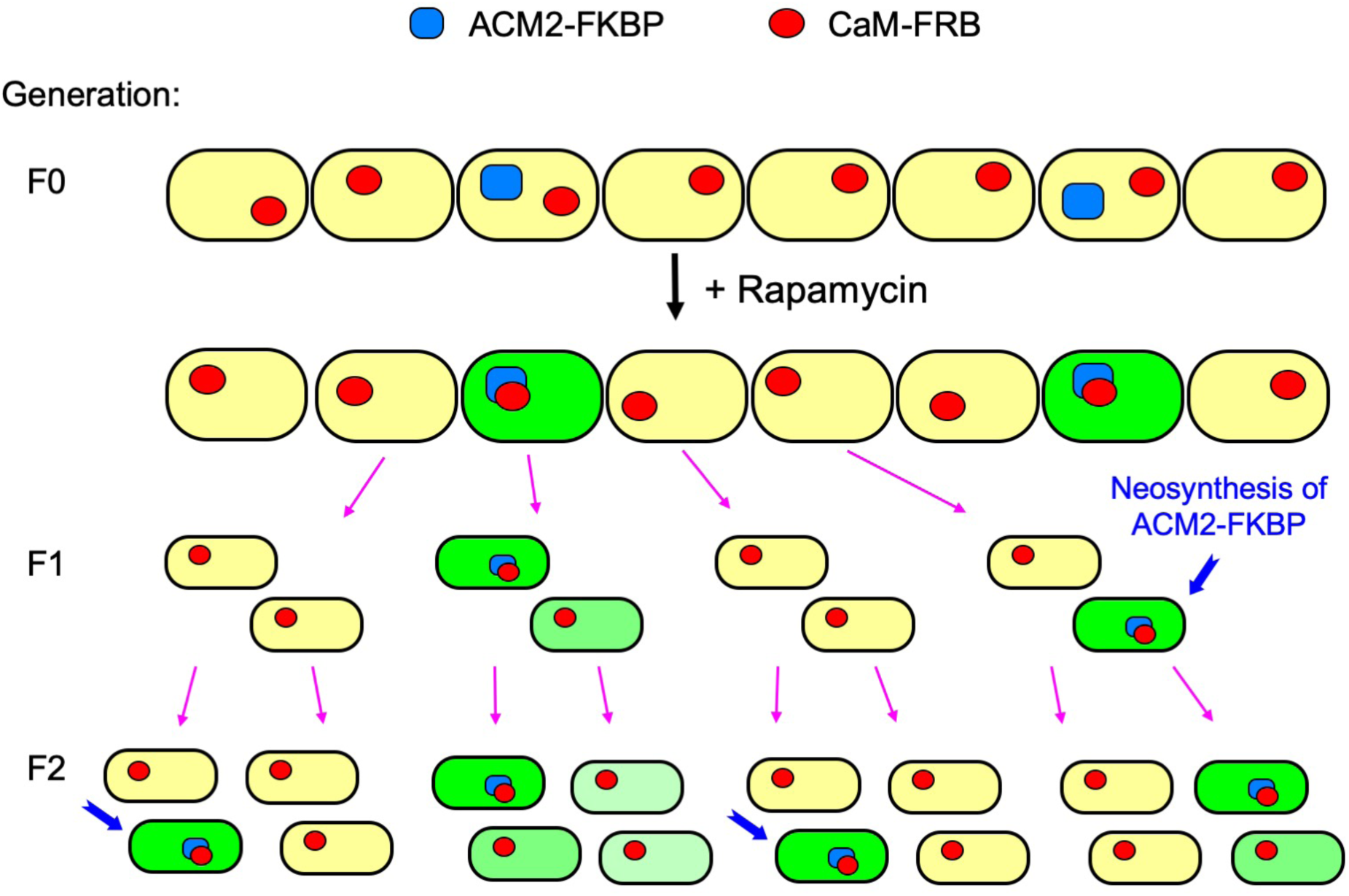
Model of phenotypic inheritance of Cya^+^ phenotype in ESACH system. In bacterial cells (yellow) harboring pACM2-Fkbp and pK1CaM-Frb, the ACM2-FKBP hybrid protein (blue) is expressed on average in 1 out 4 bacterial cells, while the CaM-FRB fusion (red) is expressed in all cells. Upon addition of rapamycin, cells harboring a ACM2-FKBP hybrid protein are becoming fluorescent as a result of cAMP-induced ZsGreen expression (green) while other remains non-fluorescent (yellow). Over time, the progeny of the ACM2-FKBP harboring cells will gradually loss the fluorescence (shaded green) due to progressive dilution of the ZsGreen protein, until new events of neo-synthesis of ACM2-FKBP (blue arrows) resume the full expression of ZsGreen.

## DISCUSSION

We describe here the design of an exquisitely sensitive genetic assay, ESACH, able to report protein interactions at the single molecule per cell level. Taking advantage of the high enzymatic activity of the *B. pertussis* adenylate cyclase (AC) when activated by its natural activator, the eukaryotic protein calmodulin (CaM), we demonstrate that AC can confer a selectable trait to a bacterial host even when it is expressed, on average, at less than one molecule per cell. This remarkable property originates from the particular characteristics of the cAMP signaling cascade, combined with the high turnover number of CaM-activated AC (k_cat_ > 1,000 s^-1^). Indeed, in a given bacterial cell, a single molecule of active AC/CaM complex can rapidly synthesize enough cAMP to saturate the CAP proteins and activate the transcription of cAMP/CAP dependent genes. When this cell divides, the daughter cell that does not inherit the single AC/CAM complex will nevertheless inherit the over-expressed metabolic enzymes (e. g. lactose or maltose-metabolizing ones) as well as the cAMP/CAP molecules produced in the mother cell (Fig. 8). Therefore, this daughter cell is able to grow for a while on the selection medium without actually harboring any active AC enzyme. As this daughter cell further divides, the concentrations of cAMP/CAP and cAMP-dependent catabolic enzymes will progressively decrease until they reach a level insufficient to sustain growth on the selective medium. These cells will resume their growth when a new stochastic event of AC expression restarts a new cycle. Although we did not precisely determine the frequency of the stochastic expression of the AC enzyme with our system, the fact that about 20-25 % of the cell population did express a Cya*^+^* phenotype when rapamycin was added to promote the interaction and subsequent activation of ACM2-FKBP by CaM-FRB, suggests that an ACM2-FKBP molecule is produced on average every two division cycles. In our experimental settings, CaM-FRB fusion was produced at an average level of about 3-10 molecules per cell so that enough CaM-FRB should be present in each cell to activate any neo-synthesized ACM2-FKBP.

The combined properties of high AC activity and of the cAMP/CAP signaling cascade in *E. coli* thus offers a unique approach to characterize protein interactions at the single-molecule-per-cell level. Single molecule experimentation has allowed exploring biological mechanisms that cannot be easily disclosed by ensemble-level experiments ^43^ ^44^ ^45^ ^46^. Most of these studies have been performed through optical imaging of GFP tagged components expressed at the minimal observable level. The ESACH system provides the extreme sensitivity required for functional characterization of biomolecular processes *in vivo* at the single molecule level.

Interestingly, it has been reported that a significant fraction of all proteins (about half of the 1000 tested) in an *E. coli* cell might be present at low levels, that is, below ten molecules per cell ^47^. Although most of these low-abundance proteins may be non-essential, a significant fraction of them were found to be implicated in genetic interactions as indicated by the patterns of synthetic sickness and/or synthetic lethality observed in experiments of double-knockout combinations ^48^. Characterization of the interaction networks *in vivo* of these low-abundance proteins at their native expression level may be instrumental to further explore their physiological function. The fact that numerous proteins may be expressed only at very low copy number in *E. coli* and possibly in other bacterial species, is calling for the development of technologies allowing such single-molecule experiments *in vivo*. Currently only high-resolution microscopy techniques are able to detect a single molecule in live cells ^45^ ^46^ ^49^. The ESACH system could provide a simple and efficient approach to study protein interactions, as well as protein modifications or degradations, at such extremely low levels of expression.

The exquisite sensitivity of the ESACH system might also be exploited for direct *in vivo* screening of high affinity antibodies (e.g. single chain antibody fragment, scFv, or nanobody ^40^) or other “binders” based on different scaffolds ^39^, to specific antigens or proteins of interest. We anticipate that ESACH could be efficiently combined with deep learning techniques to facilitate identification and evolution of recombinant antibodies or *in silico*-designed biomolecules with high affinity and selectivity ^50^ ^51^ ^52^ ^53^ ^54^ ^55^.

In summary, we have shown here that the high catalytic activity of the CaM-activated AC can be combined with the endogenous cAMP signaling cascade of *E. coli* to design a novel experimental tool for exploring biological processes at the single molecule level in living cells. This may open novel avenues for studying *in vivo* the molecular basis of biological mechanisms that haven’t been accessible up to now with current methodologies. In addition, it could also be exploited to design synthetic regulatory networks or biological logic-gates ^56^ operating at, or even below, one molecule per cell. The ESACH technique could also be an attractive experimental tool to explore the molecular basis of phenotypic heterogeneity arising from random segregation of a small number of proteins or complexes between daughter cells during cell division ^57^ ^58^ ^59^ ^60^ ^61^. These stochastic events are keys for cell-to-cell variability leading to bacterial individuality in microbial populations and are likely an important mechanism for adaptation and evolution ^62^.

### Experimental Procedures

#### General methods

Bacteria were routinely grown at 30°C in LB broth (0.5% yeast extract, 1% tryptone) containing 0.5% NaCl ^63^. Unless stated otherwise, antibiotics were added at the following concentrations: ampicillin (100 μg/ml), chloramphenicol (30 μg/ml), kanamycin (50 μg/ml). Standard protocols for molecular cloning, PCR, DNA analysis, and transformation were used ^63^. The *E. coli* strain XL1-Blue (Stratagene / Agilent Technologies) was used for all routine cloning experiments. PCR primer’s synthesis and DNA sequencing were carried out by the company Eurofins MWG Operon (Cologne, Germany). Synthetic genes coding for the V_9A_ and V_1K_ V_H_H, barnase and barstar were obtained from Geneart (Thermo Fisher Scientific).

#### Plasmid constructions

Plasmids coding for AC wild-type, ACM247, ACM335 and CaM were described in Ladant et al. ^27^ and Vougier et al. ^30^. The plasmids pAC0, pACM1 and pACM2 are derived from the pT25 plasmid harboring a p15A origin of replication and a chloramphenicol resistant marker ^21^. They were constructed by subcloning the full-length AC (or ACM1 = ACM247; ACM2 = ACM335) genes and removal of the transcriptional and translational sequences in front of AC or ACM coding regions. The maps and DNA sequences (full-length sequence for pAC0 and, for other plasmids only the specific *Cla*I-*Xho*I DNA fragments of the other plasmids) of these plasmids are shown in Annex 1 of Supplementary Materials.

Genes coding for GFP (from plasmid pDSW207-blr ^64^, FKBP (kindly provided by Dr Yves Jacob,Institut Pasteur), Barnase (synthetic gene, ordered from Geneart, Thermo Fisher Scientific) GCN4 leucine zipper (from plasmid pUT18C-zip ^65^), and the first transmembrane segment of OppB followed by the GCN4 leucine zipper (from plasmid pUTM18C-zip ^24^) were subcloned in the C-terminal multicloning site. Plasmid pACM2-TM-zip was constructed by subcloning between the *BamH*I and *Xho*I sites of pACM2-Gfp, a PCR-amplified DNA fragment (with appropriate primers introducing a *Bgl*II site -compatible with *BamH*I - and a *Xho*I site) that codes for the linker region, the *E. coli* OppB first TM segment and the GAL4 leucine zipper (zip) domain from plasmid pUT18C-TM-zip ^24^.

Plasmids pTrACM1-Fkbp, pTrACM2-Fkbp and pTrACM2-Gfp are derivatives of plasmid pTRAC384GK ^30^, in which the AC coding region is under the control of the phage λ Pr promoter, that is repressed at temperature below 32 °C by the thermosensitive λ repressor cI^857^ (Fig. 1B and Annex 1). They were constructed by subcloning appropriate fragments from plasmid pACM1-Fkbp, pACM2-Fkbp and pACM2-Gfp into pTRAC384GK.

Plasmid pDLTCaM41 ^30^ harbors a ColE1 origin of replication, an ampicillin resistant marker, and a wild-type CaM gene under the control of the phage λ Pr promoter that is repressed at temperature below 32 °C by the thermosensitive λ repressor cI^857^ (Fig. 1B). Plasmid pTCam is a derivative of pDLTCaM41, expressing, under control of the thermoinducible λ promoter, CaM fused to a 6-histidine tag at its N-ter and a tetra-cysteine tag at its C-terminus (Annex 1). Plasmid pTCam-V9A is a derivative of pTCam expressing CaM fused to the V_9A_ nanobody at its N-ter and a tetra-cysteine tag at C-terminus (Fig. 1B and Annex 1).

Plasmids pK1Cam and pK2Cam are derivatives of the pMK-RQ vector (Geneart, Thermo Fisher Scientific; contains a ColE1 origin of replication and a Kanamycin resistant marker) and harbor a synthetic CaM gene (Geneart) with a N-terminal multicloning site and an AviTag sequence (allowing for enzymatic biotinylation *in vivo*) appended to its C-terminus. The CaM gene is placed under the control of a T7 promoter with (pK1Cam) or without (pK2Cam) a ribosome binding site (Fig. 1B, Table S1, and Annex 1). Plasmid pDL1312, used here as a control plasmid, is the parental vector of pDLTCaM41 expressing neurocalcin instead of CaM ^66^. Genes coding for the camelidae V_H_H 3G9A (here called V_9A_) and V_H_H 3K1K, (here called V_1K_) ^29^, FRB, and barstar were subcloned in frame into the N-terminal multicloning site. Plasmids pK1Cam-V_9A_-Zip and pK2Cam-V_9A_-Zip (Fig. 4A Table S1, and Annex 1) were obtained by cloning a 274 bp *BstB*I-*Xho*I fragment from pACM2-Zip (encoding AC residues 373-400 followed by the GCN4 leucine zipper) between the *Cla*I and *Xho*I sites of the pK1Cam-V_9A_ or pK2Cam-V_9A_ vectors.

Plasmids pK1Cam-V_9A_-TM-Zip (Fig. 4A Table S1, and Annex 1) was obtained by cloning between the into *Xho*I & *BamH*I sites of pK1Cam-V_9A_ a PCR-amplified fragment from pACM2-TM-Zip, encoding the TM(OppB)-Zip segment.

Plasmid pK1Cam-Frb-Zs was obtained by inserting, between the *Spe*I and *BamH*I sites of pK1Cam-Frb plasmid, a PCR fragment encoding the ZsGreen fluorescent protein gene (from plasmid pZsGreen from Clontech, Takara) placed under a pLac promoter (see Fig. 5A Table S1, and Annex 1).

Plasmid pCaM_VU8_ was synthetized by Geneart and contains a synthetic gene encoding the CaM-VU8 variant fused to a HA tag and a multicloning site, cloned into the pMA-T vector (Geneart). Plasmid pCaM_VU8_-V_1K,_ was obtained by inserting a PCR fragment encoding the V_1K_ gene between the *Mlu*I and *Sac*I sites of pCaM_VU8_. Plasmid pCaM_Cter_-V_1K_ was obtained by inserting a PCR fragment encoding the C-terminal moiety of CaM (residues 77 to 149) between the *Nhe*I and *Spe*I sites of pCaM_VU8_-V_1K_

The sequences of all recombinant plasmids were verified by DNA sequencing (Eurofins MWG, Cologne, Germany).

#### ESACH complementation and screening assays

ESACH complementation assays were carried out in the *E. coli Δcya* strain DHM1 ^22^. After transformation with appropriate plasmids, cells were plated on LB agar containing X-Gal, IPTG plus antibiotics and incubated at 30°C for 24-36 hours. Efficiency of interaction between hybrid proteins was quantified by measuring β- galactosidase (β-Gal) activity in liquid cultures in 96-well format assay ^64^. For each set of transformation, the β-Gal assays were performed on eight overnight cultures each inoculated with distinct colony and grown at 30°C in 300 μL LB broth in the presence of 0.5 mM IPTG and appropriate antibiotics in a 96-well microtiter plate (2.2 ml 96-well storage plate, Thermo Fisher Scientific). For screening experiments, the DHM1 cells, after electroporation with appropriate plasmids, were incubated in LB broth at 30°C for 90 min, then washed several times with M63 synthetic medium ^63^ and spread on M63 solid medium supplemented with maltose (0.2%), 5-bromo-4-chloro-3-indolyl-b-D-galactoside (X-Gal, 40 μg/ml), isopropyl-β-D-galactopyronoside (IPTG, 0.5 mM), kanamycin (25 μg/ml) and chloramphenicol (20 μg/ml). Plates were incubated at 30°C for 6-10 days until appearance of blue, Cya^+^ (Mal^+^ and Lac^+^) colonies. Cyclic AMP was measured on boiled liquid culture with an ELISA assay as previously described ^21^ ^67^.

#### Purification of ACM and CaM fusion proteins

The ACM1-FKBP, ACM2-FKBP and ACM2-GFP ORFs were sub-cloned into plasmid pTRAC384GK ^30^ and the recombinant plasmids (pTrACM1-FKBP, pTrACM2-FKBP and pTrACM2-GFP, respectively) were transformed into the *E. coli* BLR strain (Novagen, Darmstadt, Germany). The transformants were grown at 30 °C in LB medium containing 100 μg/ml ampicillin and when the culture reached an optical density of 0.6 – 0.8 at 600 nm, expression of the proteins was triggered by shifting the growth temperature to 42 °C. After 150 min of additional growth at 42 °C, the cells were collected by centrifugation (20 min, 10,000 x *g*, 4 °C), and the cell pellets were frozen at - 20 °C. The cell pellets were resuspended in 20 mM HEPES-Na, pH 7.5, and disrupted by sonication at 4 °C. The sonicated suspension was centrifuged for 20 min at 13,000 x *g* at 4 °C. The supernatant was discarded, and the pellet (containing inclusion bodies) was resuspended in 8 M urea with 20 mM HEPES-Na (pH 7.5) and agitated overnight at 4 °C. After 20 min of centrifugation at 13,000 x *g* at 4 °C, the supernatant (“urea extract”) that contained the solubilized ACM fusion proteins was collected. The ACM fusions were purified by two sequential chromatographic treatments with DEAE-Sepharose as previously described ^30^. Briefly, the urea extract was first loaded onto a DEAE-Sepharose column (20 ml of packed resin) equilibrated in 8 M urea with 20 mM HEPES-Na (pH 7.5). In these conditions, the ACM proteins did not bind to the resin, and were recovered in the flow-through fractions. The collected flow-through fractions were then diluted 5 times with 20 mM HEPES-Na (pH 7.5) and applied to a second DEAE-Sepharose column (20 ml of packed resin), which had been equilibrated in 20 mM HEPES-Na (pH 7.5). In these conditions, the ACM proteins were retained on the resin and, after extensive washing with 20 mM HEPES-Na (pH 7.5), the proteins were eluted in a soluble form using 20 mM HEPES-Na (pH 7.5) containing 200 mM NaCl. The AC protein preparations were analyzed by SDS-PAGE analysis (see Supplementary Material Figure S2). The concentrations of ACM fusion proteins were determined from the absorption spectra using an extinction coefficient calculated from their amino acid content.

The FRB-CaM and CaM-V_9A_-Zip protein were expressed in *E. coli* after subcloning the corresponding gene into the pTCam plasmid derivative and purified as described in Vougier et al. ^30^. Protein purity was monitored by SDS-PAGE analysis and the protein concentrations were determined by absorption at 280 using molecular extinction coefficients calculated from their amino acid sequence.

#### *In vitro* assays of the rapamycin-induced activation of ACm-FKBP by CaM-FRB

AC assays were carried out as described by Davi et al ^32^, in a final volume of 100 μL of 50 mM Tris-HCl, pH 8.0, 7.5 mM MgCl2, 0.1 mM CaCl2, 0.5 mg/mL bovine serum albumin (BSA), containing 0.5 nM of ACM1-FKBP or ACM2-FKBP, various concentrations of CaM-FRB (from 0 to 100 μM) (all proteins were diluted in 10 mM Tris-HCl, pH 8.0, 0.1% Tween 20) and supplemented with 1 mM EGTA and /or 5 μM rapamycin, as indicated. The reaction mixtures were equilibrated at 30 °C for 10 min and the enzymatic reactions were initiated by adding 2 mM ATP and further incubated at 30 °C for 20 min. Blank assays containing no enzymes or no CaM-FRB were carried out in parallel. The cAMP produced was then determined spectrophotometrically as previously described ^32^.

#### Quantification of the *in vivo* expression of the hybrid proteins

For *in vivo* quantification of hybrid protein expression, DHM1 cells harboring the appropriate plasmids were grown until mid-log phase (starting from an over-night culture) washed with M63 medium, resuspended in 8 M Urea, 20 mM Hepes-Na pH 7.5, sonicated, and then mixed with SDS Gel loading buffer and heated at 70 °C for 5 min. Total bacterial extracts, corresponding to ≈ 10^9^ bacteria (i.e. 1 OD_600_ of bacteria) were loaded on a 10% SDS-polyacrylamide gel as well as various amounts (ranging from 10 to 0.01ng) of the purified ACM2-FKBP (521 aa, molecular weight = 56,443 Da; 0.1 ng of ACM2-FKBP fusion correspond to ≈ 1 x 10^9^ protein molecules), ACM2-GFP (664 aa; molecular weight = 72,667 Da; 0.1 ng of ACM2-GFP fusion correspond to ≈ 8 x 10^8^ protein molecules),or CaM-V_9A_-Zip (381 aa; molecular weight = 41,869 Da; 0.1 ng of CaM-V_9A_-Zip fusion correspond to ≈ 1.4 x 10^9^ molecules) hybrid proteins. After electrophoresis, the proteins were electro-transferred onto a polyvinylidene difluoride membrane (Millipore), incubated with the anti-cyaA monoclonal antibody 3D1 ^33^ (Santa Cruz Biotechnology; sc-13582) then with a horseradish peroxidase-conjugated goat anti-mouse secondary antibody (Santa Cruz Biotechnology, sc-2031) and finally detected by enhanced chemiluminescence (ECL-Plus kit from Amersham Biosciences (Cytiva) or the SuperSignal West Atto Ultimate kit from Thermo Fisher Scientific).

#### Mutagenesis of the V_9A_ gene and ESACH screening of antigen-antibody interactions

Mutagenesis of the V_9A_ gene on plasmid pK2Cam-V_9A_ was performed by a mutagenic PCR carried out with an oligonucleotide primer containing a degenerated codon NN(C/G) at position 107 (Fig. S3). The mutagenized plasmid pool was transformed into DHM1/pACM2-Gfp competent bacteria and plated on LB agar supplemented with X-gal (40 μg/ml), IPTG (0.5 mM), kanamycin (50 μg/ml), chloramphenicol (30 μg/ml) plates and grown for 36 hrs at 30 °C. Several white colonies, encoding non-interacting V_9A_ variants, were randomly picked for plasmid purification and the corresponding pK2Cam plasmids were sequenced. We selected one of these variants, pK2Cam-V_9A_-Y_107N_, in which the Y_107_ codon of V_9A_ is changed into an Asn codon for a second round of mutagenesis. It was performed by a mutagenic PCR carried out with oligonucleotide primers containing degenerated codons NN(C/G) at position 105-107 (Fig. S3). The mutagenized plasmid pool was transformed into DHM1/pACM2-Gfp competent bacteria and plated on LB-Xgal-IPTG-chloramphenicol-kanamycin plates and grown for 36 hrs at 30 °C. Both white (lac-) and blue (lac+) colonies were randomly picked for plasmid purification and the pK2Cam plasmids were sequenced (Eurofins MWG, Cologne, Germany). Alternatively, the transformed cells were plated on a selective medium made of M63 minimal medium agar supplemented with maltose (0.2%), kanamycin (25 μg/ml), chloramphenicol (20 μg/ml), IPTG (0.5 mM), and X-gal (40 μg/ml, to facilitate the detection of Cya+ clones that are Mal+ and also Lac+) and grown for several days at 30 °C ^41^ ^42^. Growing colonies(mal+) were randomly picked for plasmid purification and the pK2Cam plasmids were sequenced as above.

#### Fluorescence microscopy

For fluorescence microscopy studies, overnight cultures of DHM1 cells harboring appropriate plasmids were diluted 1:100 in LB medium containing IPTG (100 μM), and appropriate antibiotics and incubated until early exponential phase at 30°C. Rapamycin (5 μM) was added to induce association of ACM-FKBP with FRB-CaM. Images of living, non-fixed cells were acquired on a Nikon epi-fluorescence microscope Eclipse 80*i* equipped with a 100x Plan-Apo oil immersion objective and a 100W mercury lamp. Images were captured with a 5-megapixel colour CCD DS-5Mc device camera and processed using Adobe Photoshop software ^64^ ^41^.

## Supporting information

Annex - DNA Sequences

## Abbreviations

AC: *Bordetella pertussis* adenylate cyclase catalytic domain
ACM1 = ACM247 =: AC mutant with a Leu-Gln insertion between residues 247 and 248
ACM2 = ACM335 =: AC mutant with a Cys-Ser insertion between residues 335 and 336
BACTH: Bacterial Adenylate Cyclase Two-Hybrid
CaM: calmodulin (human)
cAMP: cyclic AMP
CAP (or CRP): catabolite activator protein
ESACH: Exquisitely Sensitive Adenylate Cyclase Hybrid system
FKBP: FK506-binding protein
FRB: FKBP-rapamycin binding domain
GFP: green fluorescent protein
IPTG: Isopropyl-β-D-thiogalactopyranoside
LB: Luria-Bertani broth
Mab: monoclonal antibody
PPI: protein-protein interactions
RBS: Ribosome Binding sequence
scFv: single chain antibody fragment
SD: standard deviation
T25, T18: AC fragments
V_H_H: camelidae variable heavy chain antibody fragment
V_9A_: camelidae variable heavy chain antibody fragment 3G9A
V_1K_: camelidae variable heavy chain antibody fragment 3K1K
WB: Western Blot
Xgal: 5-Bromo-4-chloro-3-indolyl-β-D-galactopyranoside
Zip: GCN4 leucine zipper motif

## ACKNOWLEDGEMENT

The project was supported by Institut Pasteur and the Centre National de la Recherche Scientifique, CNRS UMR 3528 (Biologie Structurale et Agents Infectieux).

## Competing interests

Both authors are co-authors on patent applications **EP 3 169 782 B1** and **US10760072 B2**, which cover the design of AC/CaM-based highly sensitive regulatory circuit to detect protein-protein interaction in bacteria with single molecule sensitivity.

## Author Contributions

Conceived and designed the experiments: DL. Performed the experiments: MD, DL. Analyzed the data: MD, DL. Wrote the manuscript: DL. Final editing: MD, DL.

## Supplementary Material

**Fig. S1:**
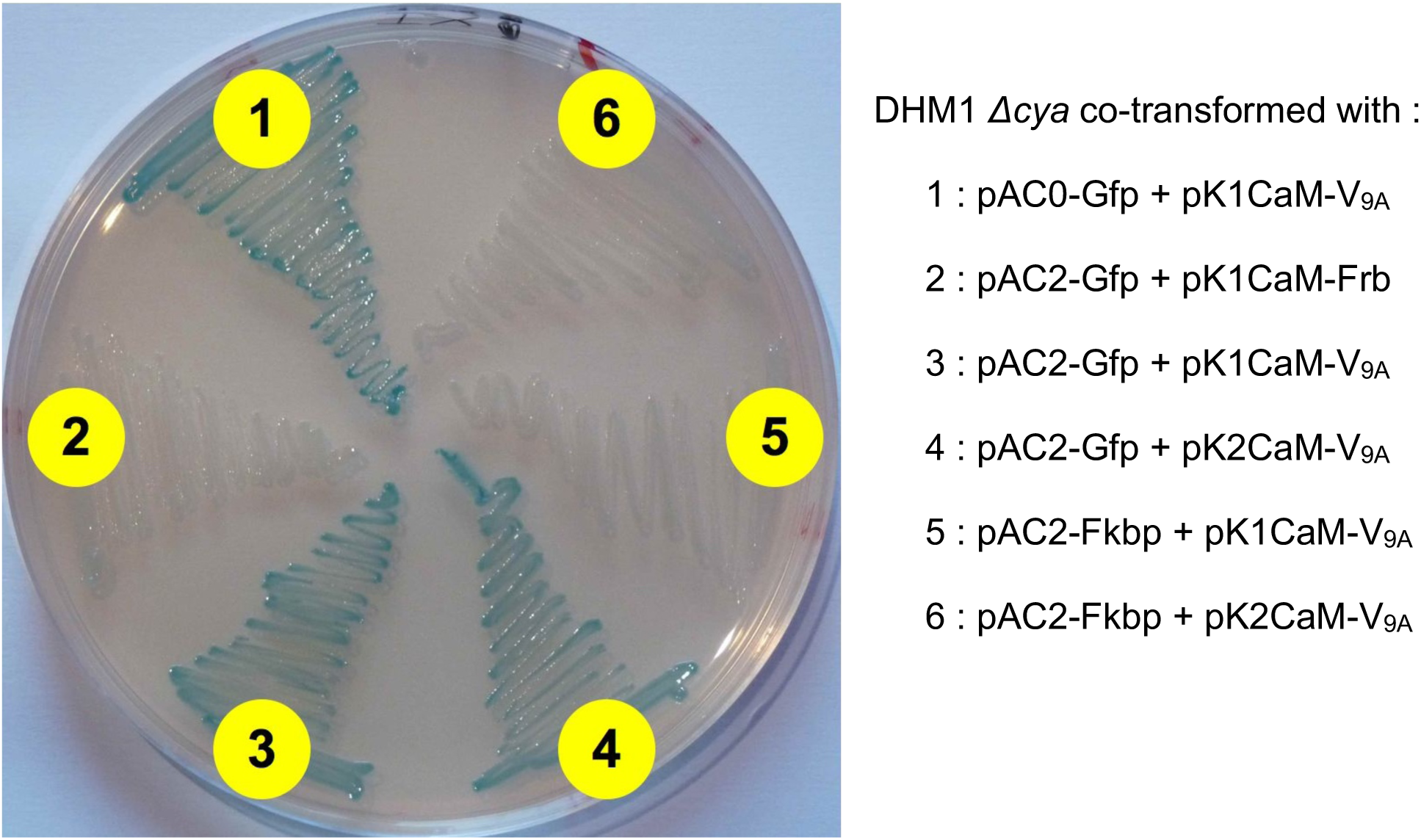
Phenotypic assay of protein interaction on LB–X-gal. DHM1 cells were co-transformed with the indicated plasmids and plated on LB–X-gal agar plates containing 0.5 mM IPTG, chloramphenicol and kanamycin, and incubated for 36 hr at 30°C.

**Fig. S2:**
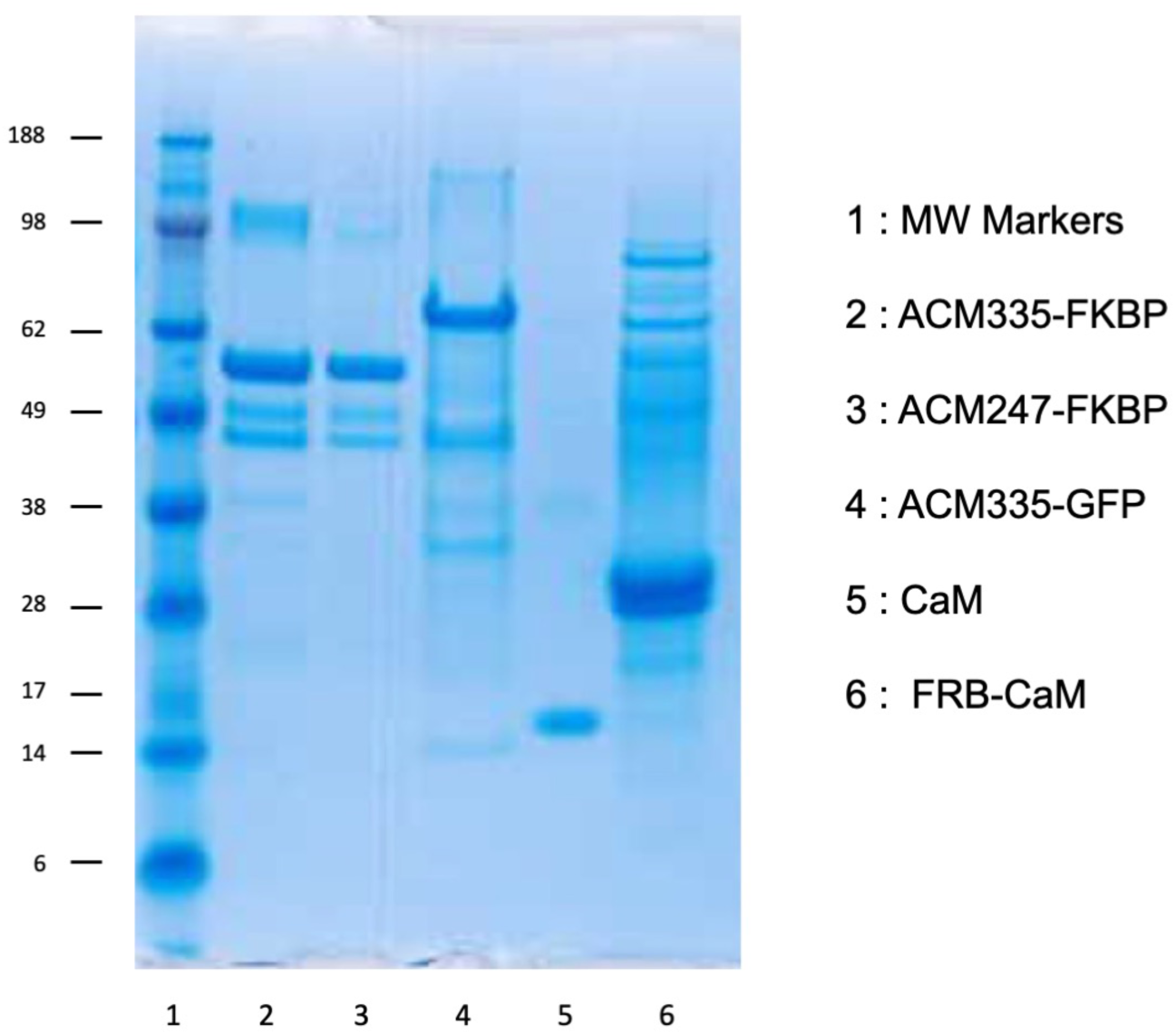
SDS-PAGE analysis of purified ACM and CaM fusions. The proteins (as indicated in the legend on the right) were separated by electrophoresis on a 4–12% SDS-polyacrylamide gel (Invitrogen). After migration, the gel was stained with PageBlue protein staining solution (Thermo Fisher Scientific).

**Fig. S3A:**
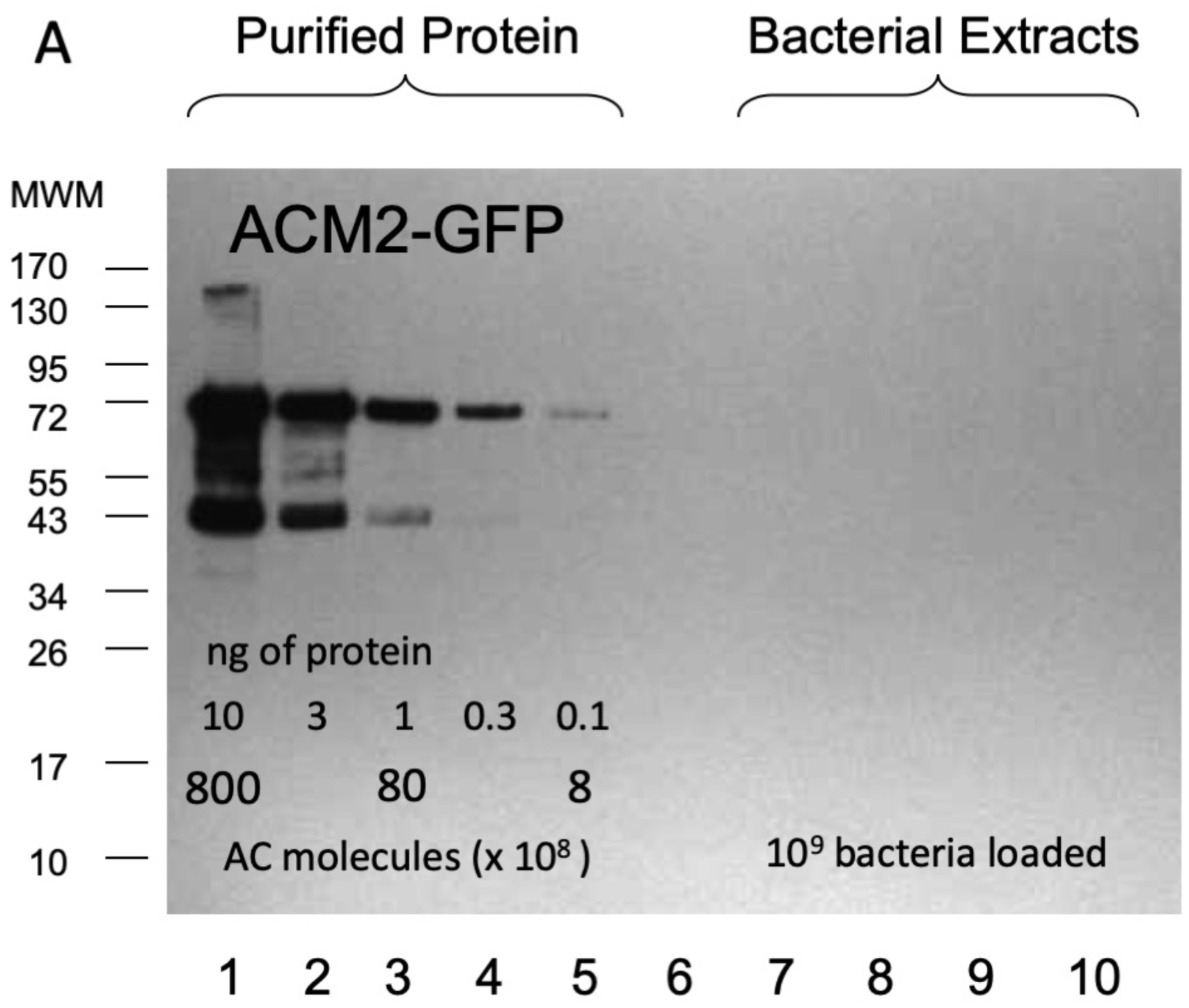
Western blot analysis of the expression of the ACM2-GFP hybrid protein in DHM1. Lines 1 - 5 : 10, 3, 1, 0.3, and 0.1ng respectively of the purified ACM2-GFP hybrid protein (molecular weight of ≈ 73kDa; 0.1 ng of ACM2-GFP fusion correspond to ≈ 8 x 10^8^ protein molecules) were separated by electrophoresis, electro-transferred to nitrocellulose and detected with 3D1 monoclonal antibody. Line 6: Molecular Weight Markers (size in kDa indicated on the left side). Line 7: no protein. Lines 8-10 : Total bacterial extracts (corresponding to ≈ 10^9^ bacteria i.e. 1 ml of cell culture at OD_600_ = 1) from DHM1 cells harboring the indicated plasmids were probed in parallel by Western blot; line 8: pACM2-Gfp/pK1C-V_9A_; line 9: pACM2-Gfp/pK2C-V_9A_; line 10: pACM2-Gfp/pTCam-V_9A_.

**Fig. S3B:**
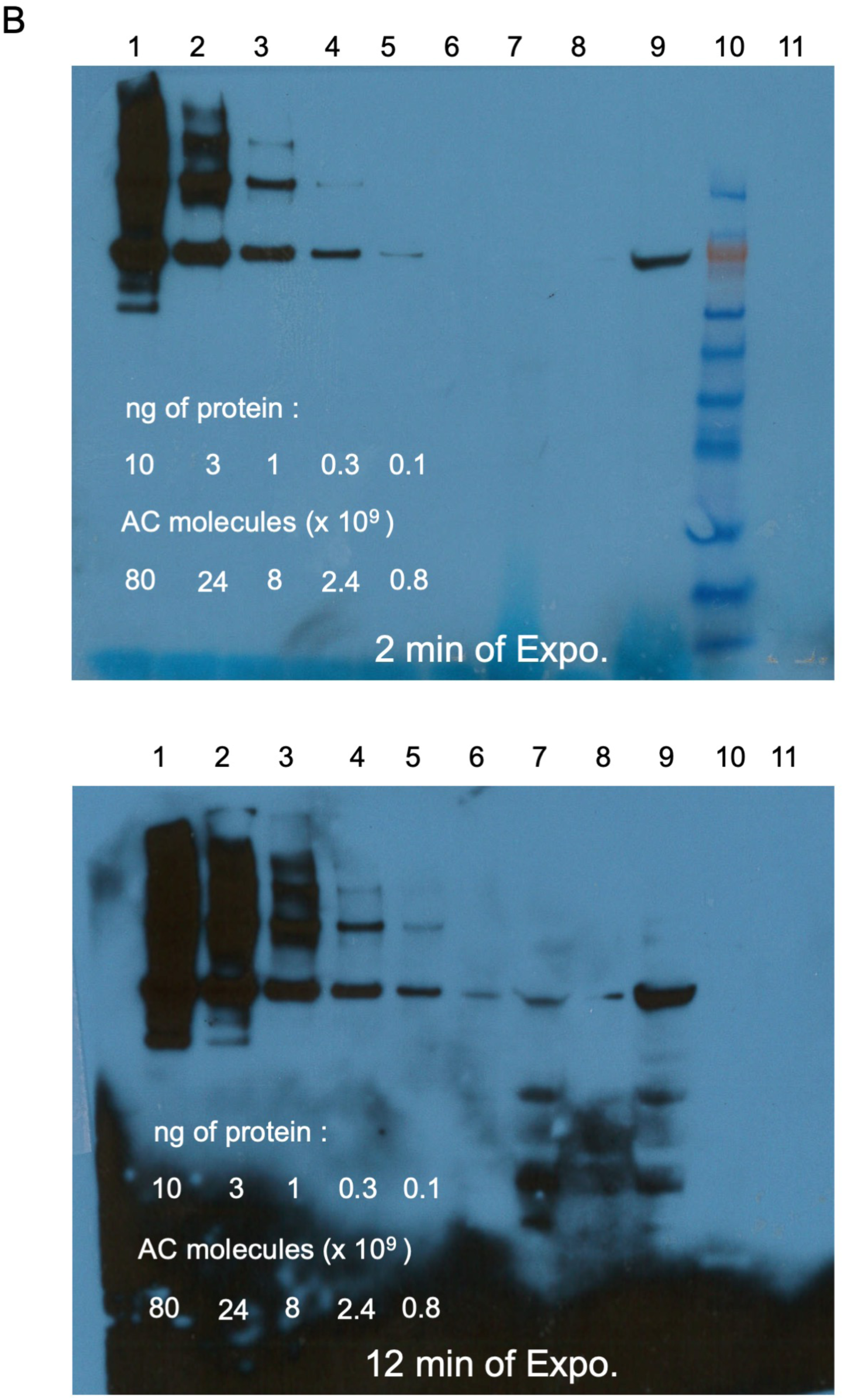
Western blot analysis of the expression of the ACM2-GFP hybrid protein in DHM1. Lines 1 - 5 : 10, 3, 1, 0.3, and 0.1ng respectively of the purified ACM2-GFP hybrid protein (molecular weight of ≈ 73kDa; 0.1 ng of ACM2-GFP fusion correspond to ≈ 8 x 10^8^ protein molecules) were separated by electrophoresis, electro-transferred to nitrocellulose, detected with 3D1 monoclonal antibody and finally revealed by enhanced chemiluminescence (2 different times of exposition are shown). Line 6: empty (the small band in the 12 min exposure blot is a carry-over of line 5). Line 7: Total bacterial extract corresponding to ≈ 10^9^ bacteria i.e. 1 ml of cell culture at OD_600_ = 1 of DHM1/pAC0-Gfp/pK1CaM-V_9A_; Line 8: empty; Line 9 : Total bacterial extract as in line 7 (ie ≈ 10^9^ DHM1/pAC0-Gfp/pK1CaM-V_9A_ bacteria) supplemented with 0.5 ng of purified ACM2-GFP protein. Line 10: Molecular Weight Markers. Line 11 : no protein.

**Fig. S4:**
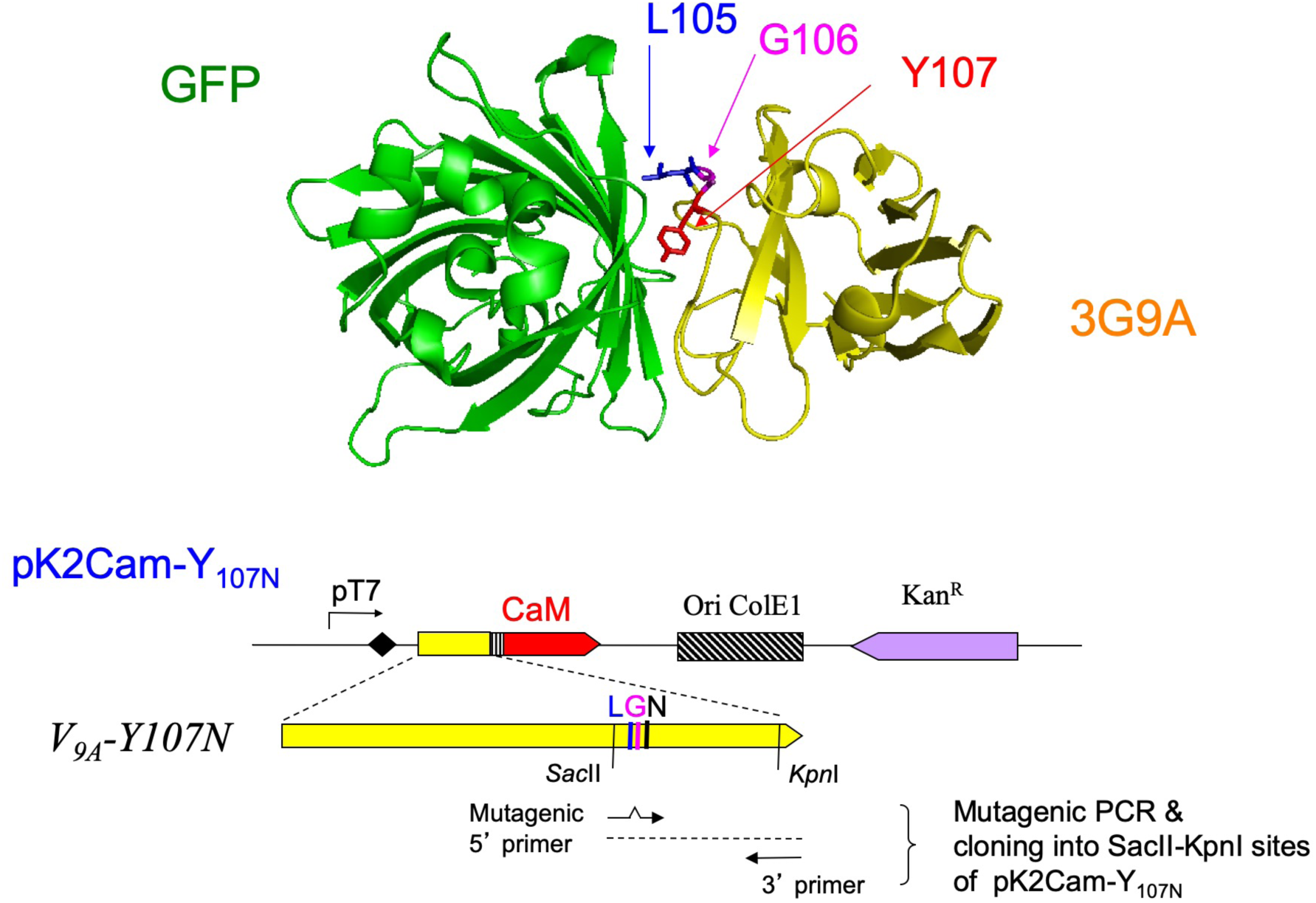
Mutagenesis of V_9A_ and selection of revertants on minimal medium Upper part: 3D structure of the GFP/ V_9A_ complex (3G9A.pdb) showing the key L_105_, G_106_, and Y_107_ residues at the protein interface. **Lower part :** Schematic representation of the pK2Cam-V_9A_-Y_107N_ plasmid and mutagenesis strategy to modify the 105-107 residues. Briefly, pK2Cam-V_9A_-Y_107N_ was mutagenized by mutagenic PCR with an oligonucleotide primer containing degenerated codons NN(C/G) at position 105-107 as described in Materials and Methods. The mutagenized plasmid pool was then transformed into DHM1/pACM2-Gfp competent bacteria and plated on an indicator medium (LB supplemented with Xgal, IPTG, chloramphenicol and kanamycin) or on a selective medium (M63 minimal medium supplemented with maltose, Xgal, IPTG, chloramphenicol and kanamycin).

**Table S1:**
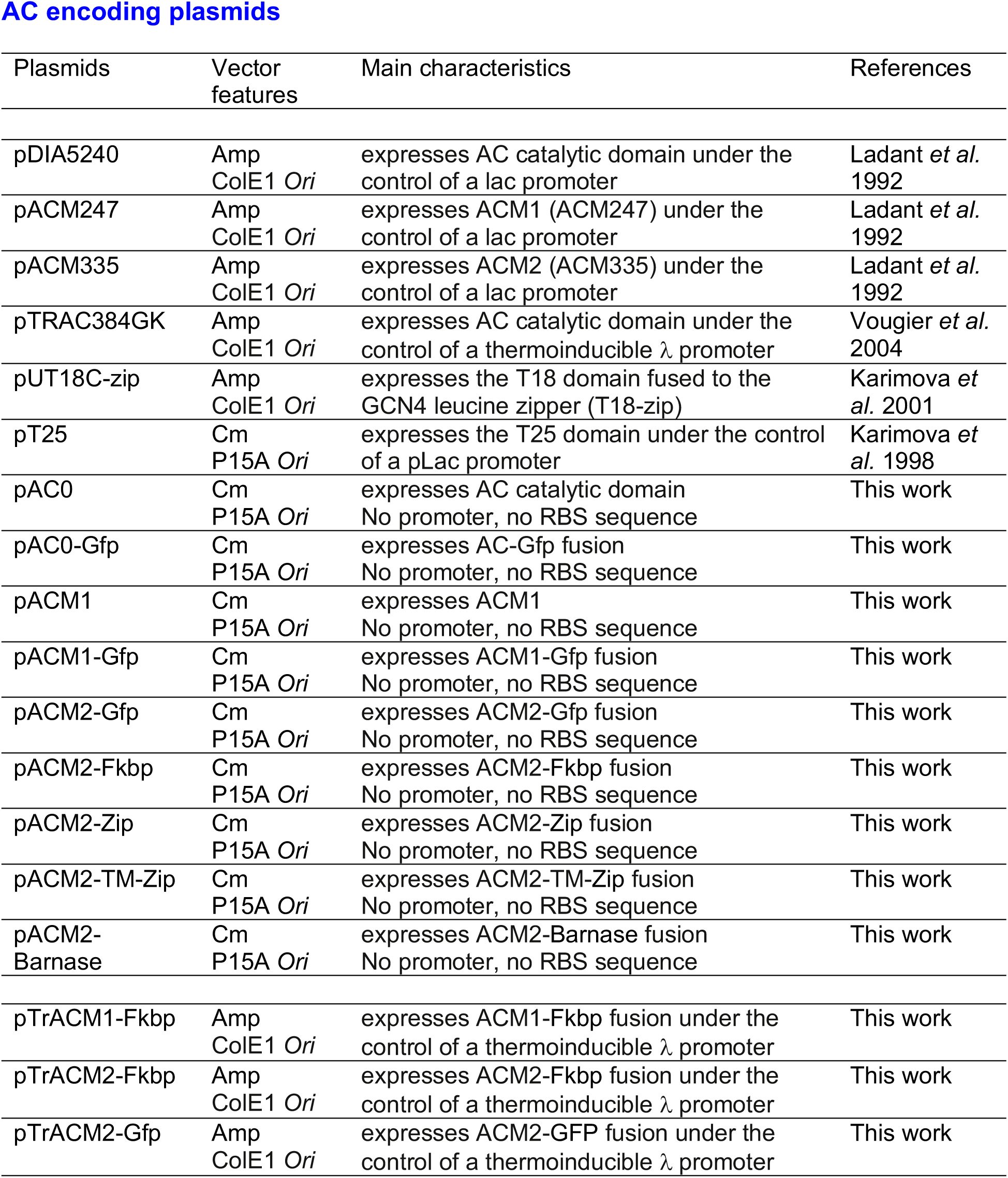
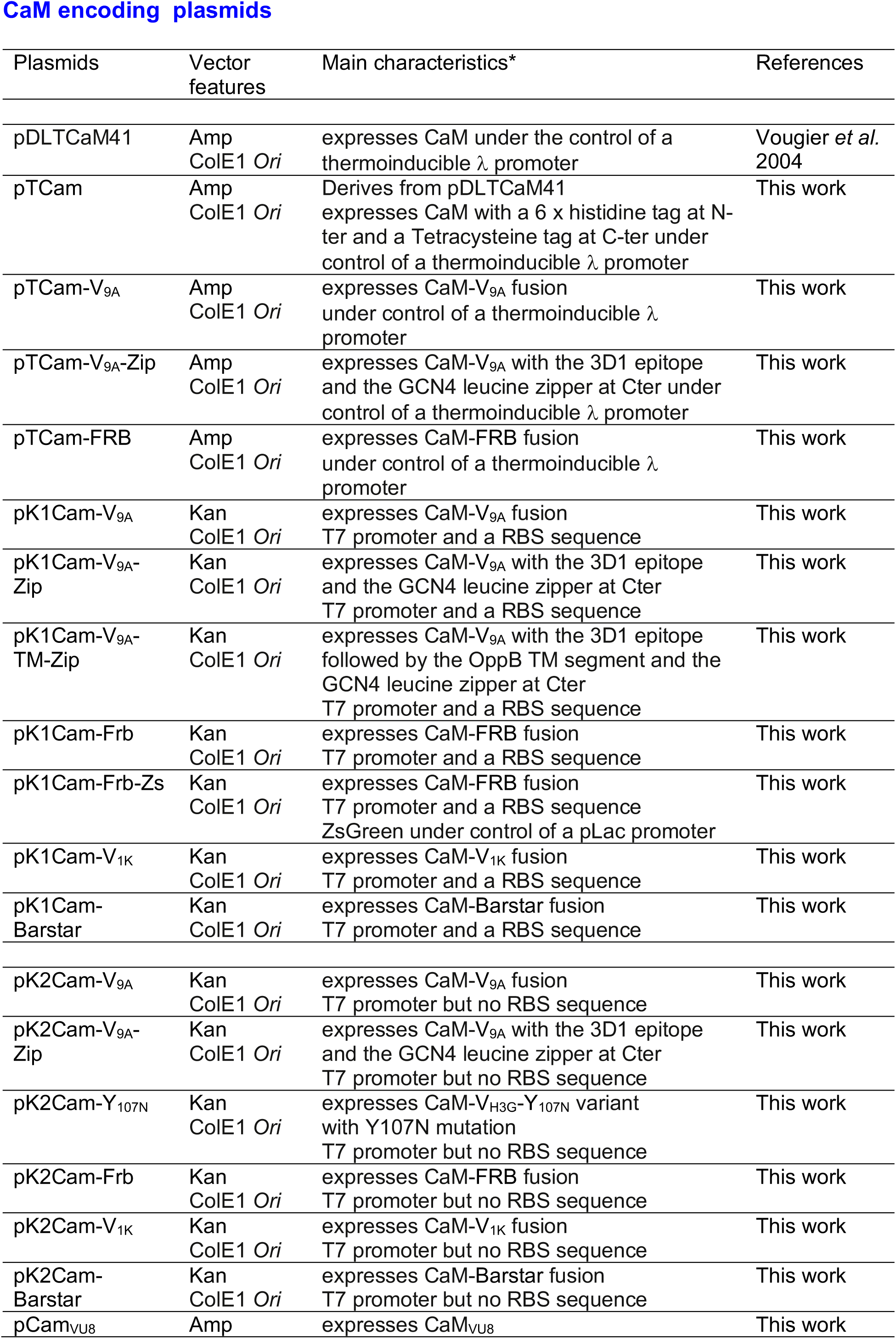
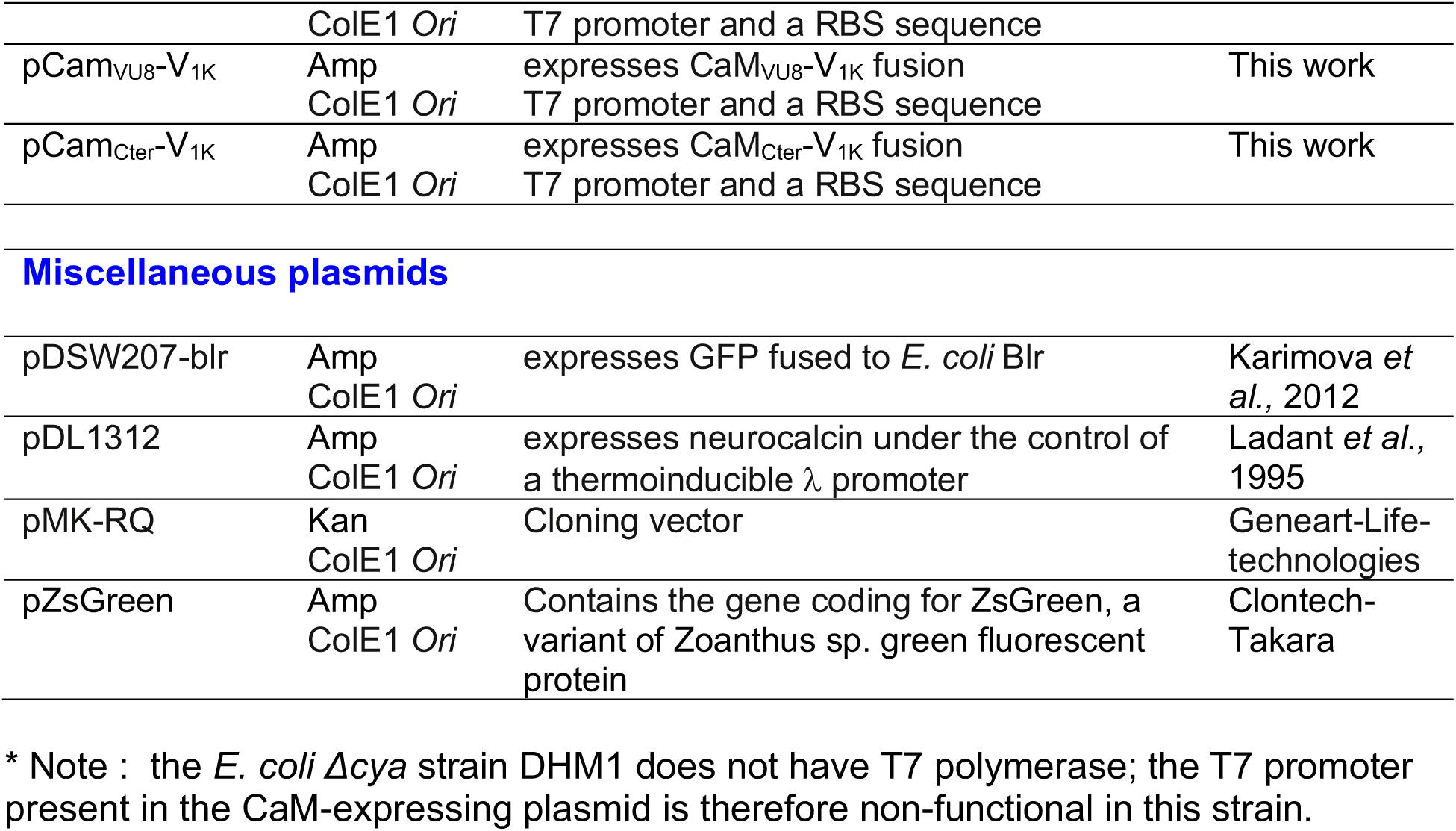
Plasmids used in this work.

**AC encoding plasmids**
| Plasmids | Vector features | Main characteristics | References |
| --- | --- | --- | --- |
| pDIA5240 | Amp<br>ColE1 *Ori* | expresses AC catalytic domain under the control of a lac promoter | Ladant *et al.* 1992 |
| pACM247 | Amp<br>ColE1 *Ori* | expresses ACM1 (ACM247) under the control of a lac promoter | Ladant *et al.* 1992 |
| pACM335 | Amp<br>ColE1 *Ori* | expresses ACM2 (ACM335) under the control of a lac promoter | Ladant *et al.* 1992 |
| pTRAC384GK | Amp<br>ColE1 *Ori* | expresses AC catalytic domain under the control of a thermoinducible $\lambda$ promoter | Vougier *et al.* 2004 |
| pUT18C-zip | Amp<br>ColE1 *Ori* | expresses the T18 domain fused to the GCN4 leucine zipper (T18-zip) | Karimova *et al.* 2001 |
| pT25 | Cm<br>P15A *Ori* | expresses the T25 domain under the control of a pLac promoter | Karimova *et al.* 1998 |
| pAC0 | Cm<br>P15A *Ori* | expresses AC catalytic domain<br>No promoter, no RBS sequence | This work |
| pAC0-Gfp | Cm<br>P15A *Ori* | expresses AC-Gfp fusion<br>No promoter, no RBS sequence | This work |
| pACM1 | Cm<br>P15A *Ori* | expresses ACM1<br>No promoter, no RBS sequence | This work |
| pACM1-Gfp | Cm<br>P15A *Ori* | expresses ACM1-Gfp fusion<br>No promoter, no RBS sequence | This work |
| pACM2-Gfp | Cm<br>P15A *Ori* | expresses ACM2-Gfp fusion<br>No promoter, no RBS sequence | This work |
| pACM2-Fkbp | Cm<br>P15A *Ori* | expresses ACM2-Fkbp fusion<br>No promoter, no RBS sequence | This work |
| pACM2-Zip | Cm<br>P15A *Ori* | expresses ACM2-Zip fusion<br>No promoter, no RBS sequence | This work |
| pACM2-TM-Zip | Cm<br>P15A *Ori* | expresses ACM2-TM-Zip fusion<br>No promoter, no RBS sequence | This work |
| pACM2-Barnase | Cm<br>P15A *Ori* | expresses ACM2-Barnase fusion<br>No promoter, no RBS sequence | This work |
| pTrACM1-Fkbp | Amp<br>ColE1 *Ori* | expresses ACM1-Fkbp fusion under the control of a thermoinducible $\lambda$ promoter | This work |
| pTrACM2-Fkbp | Amp<br>ColE1 *Ori* | expresses ACM2-Fkbp fusion under the control of a thermoinducible $\lambda$ promoter | This work |
| pTrACM2-Gfp | Amp<br>ColE1 *Ori* | expresses ACM2-GFP fusion under the control of a thermoinducible $\lambda$ promoter | This work |

CaM encoding plasmids
| Plasmids | Vector features | Main characteristics* | References |
| --- | --- | --- | --- |
| pDLTCaM41 | Amp<br>ColE1 Ori | expresses CaM under the control of a thermoinducible $\lambda$ promoter | Vougier *et al.* 2004 |
| pTCam | Amp<br>ColE1 Ori | Derives from pDLTCaM41<br>expresses CaM with a 6 x histidine tag at N-ter and a Tetracysteine tag at C-ter under control of a thermoinducible $\lambda$ promoter | This work |
| pTCam-V<sub>9A</sub> | Amp<br>ColE1 Ori | expresses CaM-V<sub>9A</sub> fusion under control of a thermoinducible $\lambda$ promoter | This work |
| pTCam-V<sub>9A</sub>-Zip | Amp<br>ColE1 Ori | expresses CaM-V<sub>9A</sub> with the 3D1 epitope and the GCN4 leucine zipper at Cter under control of a thermoinducible $\lambda$ promoter | This work |
| pTCam-FRB | Amp<br>ColE1 Ori | expresses CaM-FRB fusion under control of a thermoinducible $\lambda$ promoter | This work |
| pK1Cam-V<sub>9A</sub> | Kan<br>ColE1 Ori | expresses CaM-V<sub>9A</sub> fusion T7 promoter and a RBS sequence | This work |
| pK1Cam-V<sub>9A</sub>-Zip | Kan<br>ColE1 Ori | expresses CaM-V<sub>9A</sub> with the 3D1 epitope and the GCN4 leucine zipper at Cter T7 promoter and a RBS sequence | This work |
| pK1Cam-V<sub>9A</sub>-TM-Zip | Kan<br>ColE1 Ori | expresses CaM-V<sub>9A</sub> with the 3D1 epitope followed by the OppB TM segment and the GCN4 leucine zipper at Cter T7 promoter and a RBS sequence | This work |
| pK1Cam-Frb | Kan<br>ColE1 Ori | expresses CaM-FRB fusion T7 promoter and a RBS sequence | This work |
| pK1Cam-Frb-Zs | Kan<br>ColE1 Ori | expresses CaM-FRB fusion T7 promoter and a RBS sequence ZsGreen under control of a pLac promoter | This work |
| pK1Cam-V<sub>1K</sub> | Kan<br>ColE1 Ori | expresses CaM-V<sub>1K</sub> fusion T7 promoter and a RBS sequence | This work |
| pK1Cam-Barstar | Kan<br>ColE1 Ori | expresses CaM-Barstar fusion T7 promoter and a RBS sequence | This work |
| pK2Cam-V<sub>9A</sub> | Kan<br>ColE1 Ori | expresses CaM-V<sub>9A</sub> fusion T7 promoter but no RBS sequence | This work |
| pK2Cam-V<sub>9A</sub>-Zip | Kan<br>ColE1 Ori | expresses CaM-V<sub>9A</sub> with the 3D1 epitope and the GCN4 leucine zipper at Cter T7 promoter but no RBS sequence | This work |
| pK2Cam-Y<sub>107N</sub> | Kan<br>ColE1 Ori | expresses CaM-V<sub>H3G</sub>-Y<sub>107N</sub> variant with Y107N mutation T7 promoter but no RBS sequence | This work |
| pK2Cam-Frb | Kan<br>ColE1 Ori | expresses CaM-FRB fusion T7 promoter but no RBS sequence | This work |
| pK2Cam-V<sub>1K</sub> | Kan<br>ColE1 Ori | expresses CaM-V<sub>1K</sub> fusion T7 promoter but no RBS sequence | This work |
| pK2Cam-Barstar | Kan<br>ColE1 Ori | expresses CaM-Barstar fusion T7 promoter but no RBS sequence | This work |
| pCam<sub>VU8</sub> | Amp | expresses CaM<sub>VU8</sub> | This work |
|  | ColE1 Ori | T7 promoter and a RBS sequence |  |
| pCam<sub>VU8-V<sub>1K</sub></sub> | Amp<br>ColE1 Ori | expresses CaM<sub>VU8-V<sub>1K</sub></sub> fusion<br>T7 promoter and a RBS sequence | This work |
| pCam<sub>Cter-V<sub>1K</sub></sub> | Amp<br>ColE1 Ori | expresses CaM<sub>Cter-V<sub>1K</sub></sub> fusion<br>T7 promoter and a RBS sequence | This work |
| Miscellaneous plasmids |  |  |  |
| pDSW207-blr | Amp<br>ColE1 Ori | expresses GFP fused to *E. coli* Blr | Karimova *et al.*, 2012 |
| pDL1312 | Amp<br>ColE1 Ori | expresses neurocalcin under the control of a thermoinducible $\lambda$ promoter | Ladant *et al.*, 1995 |
| pMK-RQ | Kan<br>ColE1 Ori | Cloning vector | Geneart-Life-technologies |
| pZsGreen | Amp<br>ColE1 Ori | Contains the gene coding for ZsGreen, a variant of *Zoanthus* sp. green fluorescent protein | Clontech-Takara |

\* Note : the *E. coli* $\Delta$ *cya* strain DHM1 does not have T7 polymerase; the T7 promoter present in the CaM-expressing plasmid is therefore non-functional in this strain.

**Table S2:**
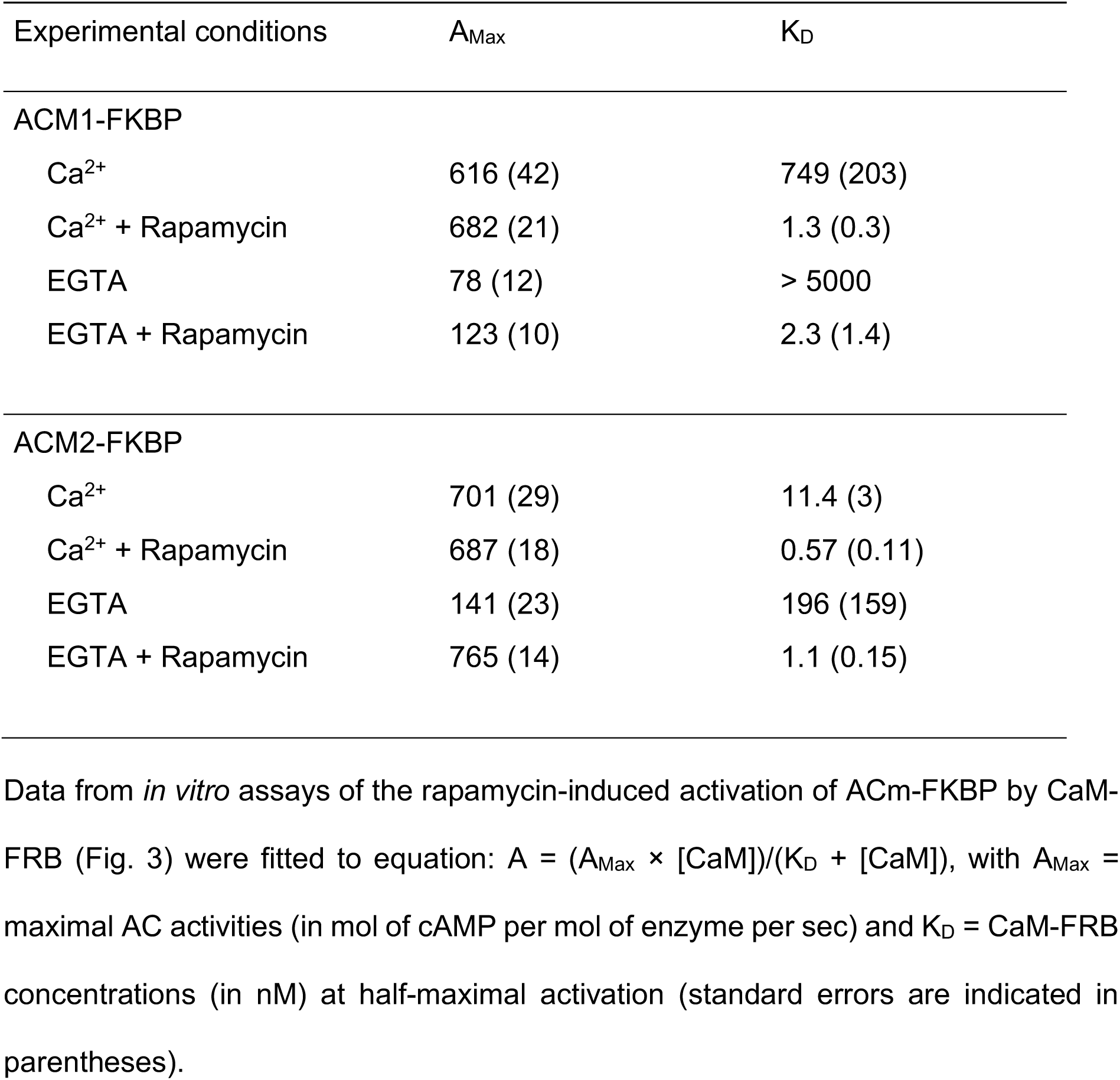
Kinetic constants of the rapamycin-induced activation of ACM-FKBP hybrids by CaM-FRB.

**Table S3:** Screening of V_9A_ variants on minimal medium.

| Alleles | Nucleotide sequences<br>at codons 105-107 | Amino acid at residues<br>105-107 |
| --- | --- | --- |
| Wt | CTG GGT TAC | L G Y |
| V<sub>9A</sub>-M1 | CTG GGT TTC | L G **F** |
| V<sub>9A</sub>-M2 | AAC GGC TAC | **N** G Y |
| V<sub>9A</sub>-M3 | CAG GGC TAC | **Q** G Y |
| V<sub>9A</sub>-M4 | CTG GGT TAC | L G Y |
| V<sub>9A</sub>-M5 | CTC GGG TAC | L G Y |
| V<sub>9A</sub>-M6 | ATG GGC TAC | **M** G Y |
| V<sub>9A</sub>-M7 | TTG GGC TAC | L G Y |
| V<sub>9A</sub>-M8 | ATG GGG TAC | **M** G Y |
| V<sub>9A</sub>-M9 | TTG GGG TAC | L G Y |
| V<sub>9A</sub>-M10 | ATG GGC TAC | **M** G Y |

## REFERENCES

1. Fields, S. & Song, O. A novel genetic system to detect protein-protein interactions. Nature 340, 245–246 (1989).

2. Gyuris, J., Golemis, E., Chertkov, H. & Brent, R. Cdi1, a human G1 and S phase protein phosphatase that associates with Cdk2. Cell 75, 791–803 (1993).

3. Johnsson, N. & Varshavsky, A. Split ubiquitin as a sensor of protein interactions in vivo. Proc. Natl. Acad. Sci. U.S.A. 91, 10340–10344 (1994).

4. Dove, S. L., Joung, J. K. & Hochschild, A. Activation of prokaryotic transcription through arbitrary protein–protein contacts. Nature 386, 627–630 (1997).

5. Rossi, F., Charlton, C. A. & Blau, H. M. Monitoring protein–protein interactions in intact eukaryotic cells by β-galactosidase complementation. Proc. Natl. Acad. Sci. U.S.A. 94, 8405–8410 (1997).

6. Pelletier, J. N., Campbell-Valois, F.-X. & Michnick, S. W. Oligomerization domain-directed reassembly of active dihydrofolate reductase from rationally designed fragments. Proc. Natl. Acad. Sci. U.S.A. 95, 12141–12146 (1998).

7. Russ, W. P. & Engelman, D. M. TOXCAT: A measure of transmembrane helix association in a biological membrane. Proc. Natl. Acad. Sci. U.S.A. 96, 863–868 (1999).

8. Eyckerman, S. et al. Design and application of a cytokine-receptor-based interaction trap. Nat Cell Biol 3, 1114–1119 (2001).

9. Paulmurugan, R. & Gambhir, S. S. Monitoring Protein−Protein Interactions Using Split Synthetic Renilla Luciferase Protein-Fragment-Assisted Complementation. Anal. Chem. 75, 1584–1589 (2003).

10. Magliery, T. J. et al. Detecting Protein−Protein Interactions with a Green Fluorescent Protein Fragment Reassembly Trap: Scope and Mechanism. J. Am. Chem. Soc. 127, 146–157 (2005).

11. Stefan, E. et al. Quantification of dynamic protein complexes using *Renilla* luciferase fragment complementation applied to protein kinase A activities *in vivo*. Proc. Natl. Acad. Sci. U.S.A. 104, 16916–16921 (2007).

12. Stynen, B., Tournu, H., Tavernier, J. & Van Dijck, P. Diversity in Genetic *In Vivo* Methods for Protein-Protein Interaction Studies: from the Yeast Two-Hybrid System to the Mammalian Split-Luciferase System. Microbiol Mol Biol Rev 76, 331–382 (2012).

13. Mehla, J., Caufield, J. H., Sakhawalkar, N. & Uetz, P. A Comparison of Two-Hybrid Approaches for Detecting Protein–Protein Interactions. in Methods in Enzymology vol. 586 333–358 (Elsevier, 2017).

14. Elhabashy, H., Merino, F., Alva, V., Kohlbacher, O. & Lupas, A. N. Exploring protein-protein interactions at the proteome level. Structure 30, 462–475 (2022).

15. Barnea, G. et al. The genetic design of signaling cascades to record receptor activation. Proc. Natl. Acad. Sci. U.S.A. 105, 64–69 (2008).

16. Kim, M. W. et al. Time-gated detection of protein-protein interactions with transcriptional readout. eLife 6, e30233 (2017).

17. Dixon, A. S. et al. NanoLuc Complementation Reporter Optimized for Accurate Measurement of Protein Interactions in Cells. ACS Chem. Biol. 11, 400–408 (2016).

18. Cassonnet, P. et al. Benchmarking a luciferase complementation assay for detecting protein complexes. Nat Methods 8, 990–992 (2011).

19. Choi, S. G. et al. Maximizing binary interactome mapping with a minimal number of assays. Nat Commun 10, 3907 (2019).

20. Tebo, A. G. & Gautier, A. A split fluorescent reporter with rapid and reversible complementation. Nat Commun 10, 2822 (2019).

21. Karimova, G., Pidoux, J., Ullmann, A. & Ladant, D. A bacterial two-hybrid system based on a reconstituted signal transduction pathway. Proc Natl Acad Sci U S A 95, 5752–5756 (1998).

22. Karimova, G., Dautin, N. & Ladant, D. Interaction network among Escherichia coli membrane proteins involved in cell division as revealed by bacterial two-hybrid analysis. J Bacteriol 187, 2233–2243 (2005).

23. Battesti, A. & Bouveret, E. The bacterial two-hybrid system based on adenylate cyclase reconstitution in Escherichia coli. Methods 58, 325–334 (2012).

24. Ouellette, S. P., Gauliard, E., Antosova, Z. & Ladant, D. A Gateway((R)) - compatible bacterial adenylate cyclase-based two-hybrid system. Environ Microbiol Rep 6, 259–267 (2014).

25. Boldridge, W. C. et al. A multiplexed bacterial two-hybrid for rapid characterization of protein–protein interactions and iterative protein design. Nat Commun 14, 4636 (2023).

26. Ladant, D. & Ullmann, A. Bordetella pertussis adenylate cyclase: a toxin with multiple talents. Trends Microbiol 7, 172–176 (1999).

27. Ladant, D., Glaser, P. & Ullmann, A. Insertional mutagenesis of Bordetella pertussis adenylate cyclase. J Biol Chem 267, 2244–2250 (1992).

28. Glaser, P. et al. Identification of residues essential for catalysis and binding of calmodulin in Bordetella pertussis adenylate cyclase by site-directed mutagenesis. EMBO J 8, 967–972 (1989).

29. Kirchhofer, A. et al. Modulation of protein properties in living cells using nanobodies. Nat Struct Mol Biol 17, 133–138 (2010).

30. Vougier, S. et al. Essential role of methionine residues in calmodulin binding to Bordetella pertussis adenylate cyclase, as probed by selective oxidation and repair by the peptide methionine sulfoxide reductases. J Biol Chem 279, 30210–30218 (2004).

31. Banaszynski, L. A., Liu, C. W. & Wandless, T. J. Characterization of the FKBP·Rapamycin·FRB Ternary Complex. J. Am. Chem. Soc. 127, 4715–4721 (2005).

32. Davi, M., Sadi, M., Pitard, I., Chenal, A. & Ladant, D. A Robust and Sensitive Spectrophotometric Assay for the Enzymatic Activity of Bacterial Adenylate Cyclase Toxins. Toxins 14, 691 (2022).

33. Lee, S. J., Gray, M. C., Guo, L., Sebo, P. & Hewlett, E. L. Epitope mapping of monoclonal antibodies against Bordetella pertussis adenylate cyclase toxin. Infect Immun 67, 2090–2095 (1999).

34. Jucovic, M. & Hartley, R. W. Protein-protein interaction: a genetic selection for compensating mutations at the barnase-barstar interface. Proc. Natl. Acad. Sci. U.S.A. 93, 2343–2347 (1996).

35. Frisch, C., Schreiber, G., Johnson, C. M. & Fersht, A. R. Thermodynamics of the interaction of barnase and barstar: changes in free energy versus changes in enthalpy on mutation 1 1Edited by J. Karn. Journal of Molecular Biology 267, 696– 706 (1997).

36. Jurėnas, D., Fraikin, N., Goormaghtigh, F. & Van Melderen, L. Biology and evolution of bacterial toxin–antitoxin systems. Nat Rev Microbiol 20, 335–350 (2022).

37. Haiech, J. et al. Affinity-based chromatography utilizing genetically engineered proteins. Interaction of Bordetella pertussis adenylate cyclase with calmodulin. J Biol Chem 263, 4259–4262 (1988).

38. Wolff, J., Newton, D. L. & Klee, C. B. Activation of Bordetella pertussis adenylate cyclase by the carboxyl-terminal tryptic fragment of calmodulin. Biochemistry 25, 7950–7955 (1986).

39. Banta, S., Dooley, K. & Shur, O. Replacing Antibodies: Engineering New Binding Proteins. Annu. Rev. Biomed. Eng. 15, 93–113 (2013).

40. Muyldermans, S. Nanobodies: Natural Single-Domain Antibodies. Annu. Rev. Biochem. 82, 775–797 (2013).

41. Karimova, G., Robichon, C. & Ladant, D. Characterization of YmgF, a 72-residue inner membrane protein that associates with the Escherichia coli cell division machinery. J Bacteriol 191, 333–346 (2009).

42. Karimova, G., Gauliard, E., Davi, M., Ouellette, S. P. & Ladant, D. Protein-Protein Interaction: Bacterial Two-Hybrid. Methods Mol Biol 1615, 159–176 (2017).

43. Li, G.-W. & Xie, X. S. Central dogma at the single-molecule level in living cells. Nature 475, 308–315 (2011).

44. Lenn, T. & Leake, M. C. Experimental approaches for addressing fundamental biological questions in living, functioning cells with single molecule precision. Open Biol. 2, 120090 (2012).

45. Gahlmann, A. & Moerner, W. E. Exploring bacterial cell biology with single-molecule tracking and super-resolution imaging. Nat Rev Microbiol 12, 9–22 (2014).

46. Elf, J. & Barkefors, I. Single-Molecule Kinetics in Living Cells. Annu. Rev. Biochem. 88, 635–659 (2019).

47. Taniguchi, Y. et al. Quantifying *E. coli* Proteome and Transcriptome with Single-Molecule Sensitivity in Single Cells. Science 329, 533–538 (2010).

48. Butland, G. et al. eSGA: E. coli synthetic genetic array analysis. Nat Methods 5, 789–795 (2008).

49. Lepore, A. et al. Quantification of very low-abundant proteins in bacteria using the HaloTag and epi-fluorescence microscopy. Sci Rep 9, 7902 (2019).

50. Parkinson, J., Hard, R. & Wang, W. The RESP AI model accelerates the identification of tight-binding antibodies. Nat Commun 14, 454 (2023).

51. Akbar, R. et al. A compact vocabulary of paratope-epitope interactions enables predictability of antibody-antigen binding. Cell Reports 34, 108856 (2021).

52. Laustsen, A. H., Greiff, V., Karatt-Vellatt, A., Muyldermans, S. & Jenkins, T. P. Animal Immunization, in Vitro Display Technologies, and Machine Learning for Antibody Discovery. Trends in Biotechnology 39, 1263–1273 (2021).

53. Watson, J. L. et al. De novo design of protein structure and function with RFdiffusion. Nature 620, 1089–1100 (2023).

54. Hie, B. L. et al. Efficient evolution of human antibodies from general protein language models. Nat Biotechnol 42, 275–283 (2024).

55. Bennett, N. R. et al. Atomically accurate de novo design of single-domain antibodies. Preprint at 10.1101/2024.03.14.585103 (2024).

56. Chen, Z. & Elowitz, M. B. Programmable protein circuit design. Cell 184, 2284– 2301 (2021).

57. Cai, L., Friedman, N. & Xie, X. S. Stochastic protein expression in individual cells at the single molecule level. Nature 440, 358–362 (2006).

58. Huh, D. & Paulsson, J. Random partitioning of molecules at cell division. Proc. Natl. Acad. Sci. U.S.A. 108, 15004–15009 (2011).

59. Huh, D. & Paulsson, J. Non-genetic heterogeneity from stochastic partitioning at cell division. Nat Genet 43, 95–100 (2011).

60. Okumus, B. et al. Mechanical slowing-down of cytoplasmic diffusion allows in vivo counting of proteins in individual cells. Nat Commun 7, 11641 (2016).

61. Bergmiller, T. et al. Biased partitioning of the multidrug efflux pump AcrAB-TolC underlies long-lived phenotypic heterogeneity. Science 356, 311–315 (2017).

62. Keegstra, J. M. et al. Phenotypic diversity and temporal variability in a bacterial signaling network revealed by single-cell FRET. eLife 6, e27455 (2017).

63. Miller, J. H. A Short Course in Bacterial Genetics: A Laboratory Manual and Handbook for Escherichia Coli and Related Bacteria. vol. Volume 1 (CSHL Press, Cold Spring Harbor, NY., 1992).

64. Karimova, G., Davi, M. & Ladant, D. The beta-lactam resistance protein Blr, a small membrane polypeptide, is a component of the Escherichia coli cell division machinery. J Bacteriol 194, 5576–5588 (2012).

65. Karimova, G., Ullmann, A. & Ladant, D. Protein-protein interaction between Bacillus stearothermophilus tyrosyl-tRNA synthetase subdomains revealed by a bacterial two-hybrid system. J Mol Microbiol Biotechnol 3, (2001).

66. Ladant, D. Calcium and membrane binding properties of bovine neurocalcin delta expressed in Escherichia coli. J Biol Chem 270, 3179–3185 (1995).

67. Karimova, G., Ullmann, A. & Ladant, D. A bacterial two-hybrid system that exploits a cAMP signaling cascade in Escherichia coli. Methods Enzymol 328, (2000).

68. Guo, Q. et al. Structural basis for the interaction of Bordetella pertussis adenylyl cyclase toxin with calmodulin. EMBO J 24, 3190–3201 (2005).

