## Supplementary material for "*In vivo* detection of protein-protein interactions with single molecule resolution": Annex - DNA Sequences

**Annex 1 : DNA sequences of ESACH plasmids.**

AC fusion encoding plasmids

pAC0

pAC0-Gfp

pACM1-Gfp

pACM2-Fkbp

pACM2-Zip

pACM2-TM-Zip

pACM2-Barnase

pTrACM2-Gfp

CaM fusion encoding plasmids

pTCam-$V_{9A}$

pTCam-$V_{9A}$-Zip

pTCam-Frb

pK1Cam-$V_{9A}$

pK2Cam-$V_{9A}$

pK1Cam-$V_{9A}$-Zip

pK1Cam-$V_{9A}$-TM-Zip

pK1Cam-Frb

pK1Cam-Frb-Zs

pK1Cam-$V_{1K}$

pK1Cam-Barstar

$pCam_{VU8}$

$pCam_{VU8}$-$V_{1K}$

$pCam_{Cter}$-$V_{1K}$

#### pAC0 (3610 bp):

#### Map

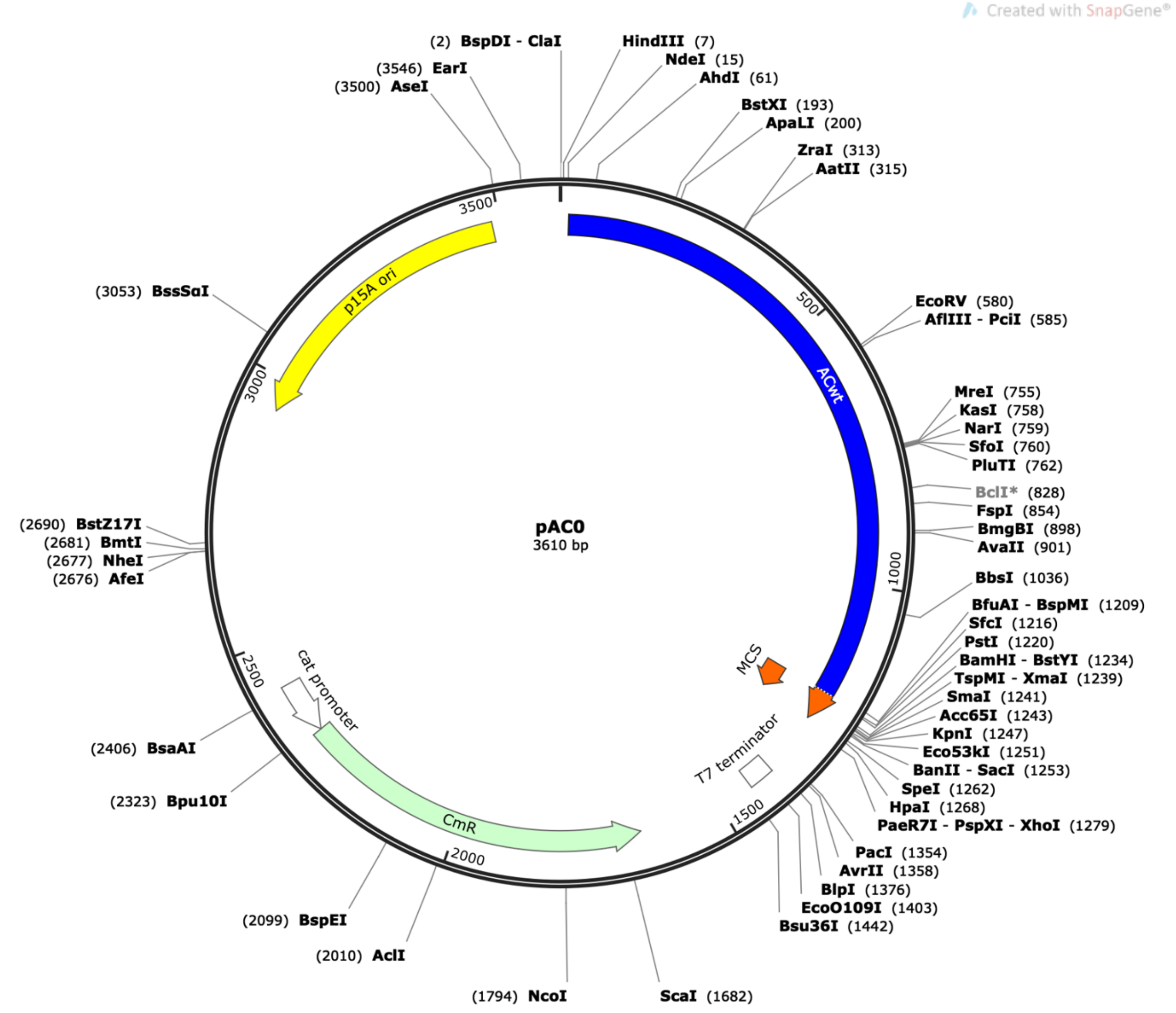


#### pAC0 full DNA sequence:

**atcgat**aagcttgcatATGCAGCAATCGCATCAGGCTGGTTACGCAAACGCCGCCGACCGGGAGTCTGGCATCCC
CGCAGCCGTACTCGATGGCATCAAGGCCGTGGCGAAGGAAAAAAACGCCACATTGATGTTCCGCCTGGTCAACCC
CCATTCCACCAGCCTGATTGCCGAAGGGGTGGCCACCAAAGGATTGGGCGTGCACGCCAAGTCGTCCGATTGGGG
GTTGCAGGCGGGCTACATTCCCGTCAACCCGAATCTTTCCAAACTGTTCGGCCGTGCGCCCGAGGTGATCGCGCG
GGCCGACAACGACGTCAACAGCAGCCTGGCGCATGGCCATACCGCGGTCGACCTGACGCTGTCGAAAGAGCGGCT
TGACTATCTGCGGCAAGCGGGCCTGGTCACCGGCATGGCCGATGGCGTGGTCGCGAGCAACCACGCAGGCTACGA
GCAGTTCGAGTTTCGCGTGAAGGAAACCTCGGACGGGCGCTATGCCGTGCAGTATCGCCGCAAGGGCGGCGACGA
TTTCGAGGCGGTCAAGGTGATCGGCAATGCCGCCGGTATTCCACTGACGGCGGATATCGACATGTTCGCCATTAT
GCCGCATCTGTCCAACTTCCGCGACTCGGCGCGCAGTTCGGTGACCAGCGGCGATTCGGTGACCGATTACCTGGC
GCGCACGCGGCGGGCCGCCAGCGAGGCCACGGGCGGCCTGGATCGCGAACGCATCGACTTGTTGTGGAAAATCGC
TCGCGCCGGCGCCCGTTCCGCAGTGGGCACCGAGGCGCGTCGCCAGTTCCGCTACGACGGCGACATGAATATCGG
CGTGATCACCGATTTCGAGCTGGAAGTGCGCAATGCGCTGAACAGGCGGGCGCACGCCGTCGGCGCGCAGGACGT
GGTCCAGCATGGCACTGAGCAGAACAATCCTTTCCCGGAGGCAGATGAGAAGATTTTCGTCGTATCGGCCACCGG
TGAAAGCCAGATGCTCACGCGCGGGCAACTGAAGGAATACATTGGCCAGCAGCGCGGCGAGGGCTATGTCTTCTA
CGAGAACCGTGCATACGGCGTGGCGGGGAAAAGCCTGTTCGACGATGGGCTGGGAGCCGCGCCCGGCGTGCCGAG
CGGACGTTCGAAGTTCTCGCCGGATGTACTGGAAACGGTGCCGGCGTCACCCGGATTGCGGCGGCCGTCGCTGGG
CGCAGTGGAACGCCACTGCAGGTCGACTCTAGAGGATCCCCGGGTACCGAGCTCGAATTCTACTAGTTAActaag

taa**ctcgag**tctggtaaagaaaccgctgctgcgaaatttgaacgccagcacatggactcgtctactagcgcagct
taattaacctaggctgctgccaccgctgagcaataactagcataaccccttggggcctctaaacgggtcttgagg
ggttttttgctgaaacctcaggcatttgagaagcacacggtcacactgcttccggtagtcaataaaccggtaaac
cagcaatagacataagcggctatttaacgaccctgccctgaaccgacgaccgggtcgaatttgctttcgaatttc
tgccattcatccgcttattatcacttattcaggcgtagcaccaggcgtttaagggcaccaataactgccttaaaa
aaattacgccccgccctgccactcatcgcagtactgttgtaattcattaagcattctgccgacatggaagccatc
acagacggcatgatgaacctgaatcgccagcggcatcagcaccttgtcgccttgcgtataatatttgcccatggt
gaaaacgggggcgaagaagttgtccatattggccacgtttaaatcaaaactggtgaaactcacccagggattggc
tgagacgaaaaacatattctcaataaaccctttagggaaataggccaggttttcaccgtaacacgccacatcttg
cgaatatatgtgtagaaactgccggaaatcgtcgtggtattcactccagagcgatgaaaacgtttcagtttgctc
atggaaaacggtgtaacaagggtgaacactatcccatatcaccagctcaccgtctttcattgccatacggaattc
cggatgagcattcatcaggcgggcaagaatgtgaataaaggccggataaaacttgtgcttatttttctttacggt
ctttaaaaaggccgtaatatccagctgaacggtctggttataggtacattgagcaactgactgaaatgcctcaaa
atgttctttacgatgccattgggatatatcaacggtggtatatccagtgatttttttctccattttagcttcctt
agctcctgaaaatctcgataactcaaaaaatacgcccggtagtgatcttatttcattatggtgaaagttggaacc
tcttacgtgccgatcaacgtctcattttcgccaaaagttggcccagggcttcccggtatcaacagggacaccagg
atttatttattctgcgaagtgatcttccgtcacaggtatttattcggcgcaaagtgcgtcgggtgatgctgccaa
cttactgatttagtgtatgatggtgtttttgaggtgctccagtggcttctgtttctatcagctgtccctcctgtt
cagctactgacggggtggtgcgtaacggcaaaagcaccgccggacatcagcgctagcggagtgtatactggctta
ctatgttggcactgatgagggtgtcagtgaagtgcttcatgtggcaggagaaaaaaggctgcaccggtgcgtcag
cagaatatgtgatacaggatatattccgcttcctcgctcactgactcgctacgctcggtcgttcgactgcggcga
gcggaaatggcttacgaacggggcggagatttcctggaagatgccaggaagatacttaacagggaagtgagaggg
ccgcggcaaagccgtttttccataggctccgcccccctgacaagcatcacgaaatctgacgctcaaatcagtggt
ggcgaaacccgacaggactataaagataccaggcgtttccccctggcggctccctcgtgcgctctcctgttcctgc
ctttcggtttaccggtgtcattccgctgttatggccgcgtttgtctcattccacgcctgacactcagttccgggt
aggcagttcgctccaagctggactgtatgcacgaaccccccgttcagtccgaccgctgcgccttatccggtaact
atcgtcttgagtccaacccggaaagacatgcaaaagcaccactggcagcagccactggtaattgatttagaggag
ttagtcttgaagtcatgcgccggttaaggctaaactgaaaggacaagttttggtgactgcgctcctccaagccag
ttacctcggttcaaagagttggtagctcagagaaccttcgaaaaaccgccctgcaaggcggttttttcgttttca
gagcaagagattacgcgcagaccaaaacgatctcaagaagatcatcttattaatcagataaaatatttctagatt
tcagtgcaatttatctcttcaaatgtagcacctgaagtcagccccatacgatataagttgtaattctcatgtttg
acagcttatc

#### pAC0-GFP (4337 bp):

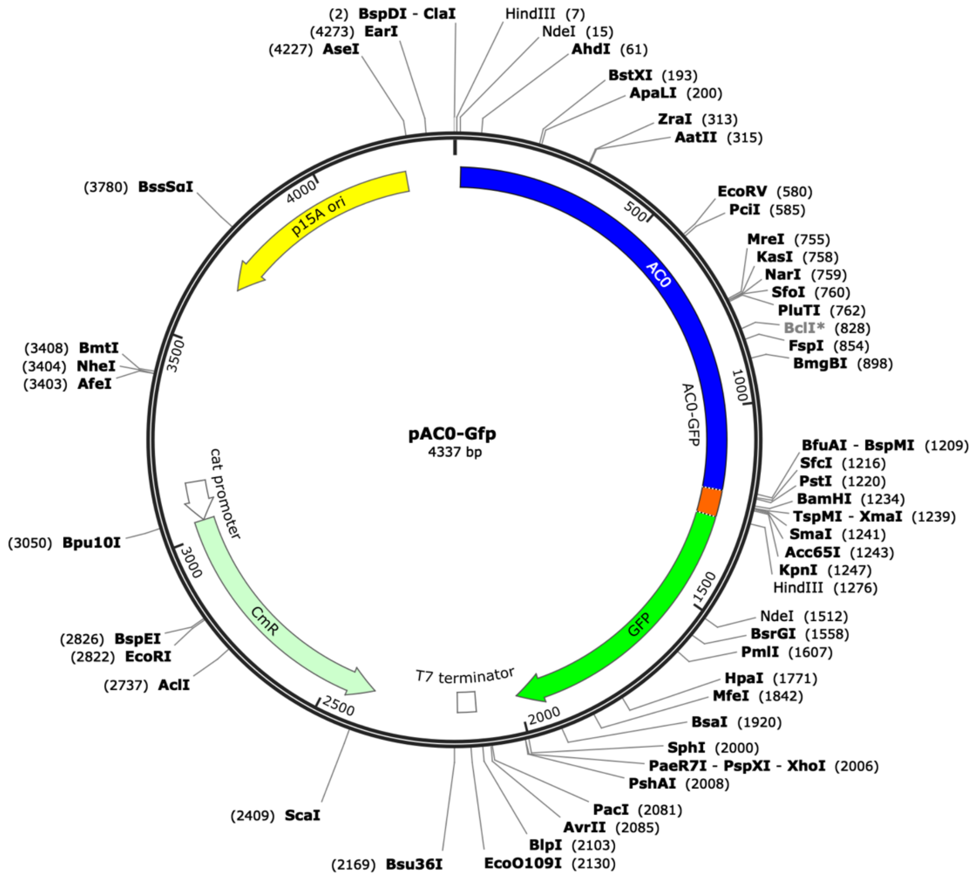


#### DNA sequence (*Cla*I-*Xho*I fragment)

**atcgat**aagcttgcatATGCAGCAATCGCATCAGGCTGGTTACGCAAACGCCGCCGACCGGGAGTCTGGCATCCC
CGCAGCCGTACTCGATGGCATCAAGGCCGTGGCGAAGGAAAAAAACGCCACATTGATGTTCCGCCTGGTCAACCC
CCATTCCACCAGCCTGATTGCCGAAGGGGTGGCCACCAAAGGATTGGGCGTGCACGCCAAGTCGTCCGATTGGGG
GTTGCAGGCGGGCTACATTCCCGTCAACCCGAATCTTTCCAAACTGTTCGGCCGTGCGCCCGAGGTGATCGCGCG
GGCCGACAACGACGTCAACAGCAGCCTGGCGCATGGCCATACCGCGGTCGACCTGACGCTGTCGAAAGAGCGGCT
TGACTATCTGCGGCAAGCGGGCCTGGTCACCGGCATGGCCGATGGCGTGGTCGCGAGCAACCACGCAGGCTACGA
GCAGTTCGAGTTTCGCGTGAAGGAAACCTCGGACGGGCGCTATGCCGTGCAGTATCGCCGCAAGGGCGGCGACGA
TTTCGAGGCGGTCAAGGTGATCGGCAATGCCGCCGGTATTCCACTGACGGCGGATATCGACATGTTCGCCATTAT
GCCGCATCTGTCCAACTTCCGCGACTCGGCGCGCAGTTCGGTGACCAGCGGCGATTCGGTGACCGATTACCTGGC
GCGCACGCGGCGGGCCGCCAGCGAGGCCACGGGCGGCCTGGATCGCGAACGCATCGACTTGTTGTGGAAAATCGC
TCGCGCCGGCGCCCGTTCCGCAGTGGGCACCGAGGCGCGTCGCCAGTTCCGCTACGACGGCGACATGAATATCGG
CGTGATCACCGATTTCGAGCTGGAAGTGCGCAATGCGCTGAACAGGCGGGCGCACGCCGTCGGCGCGCAGGACGT
GGTCCAGCATGGCACTGAGCAGAACAATCCTTTCCCGGAGGCAGATGAGAAGATTTTCGTCGTATCGGCCACCGG
TGAAAGCCAGATGCTCACGCGCGGGCAACTGAAGGAATACATTGGCCAGCAGCGCGGCGAGGGCTATGTCTTCTA
CGAGAACCGTGCATACGGCGTGGCGGGGAAAAGCCTGTTCGACGATGGGCTGGGAGCCGCGCCCGGCGTGCCGAG
CGGACGTTCGAAGTTCTCGCCGGATGTACTGGAAACGGTGCCGGCGTCACCCGGATTGCGGCGGCCGTCGCTGGG
CGCAGTGGAACGCCACTGCAGGTCGACTCTAGAGGATCCCCGGGTACCGGGGGGGTCTATGACCATGATTACGCC
AAGCTTGATGAGTAAAGGAGAAGAACTTTTCACTGGAGTTGTCCCAATTCTTGTTGAATTAGATGGTGATGTTAA
TGGGCACAAATTTTCTGTCAGTGGAGAGGGTGAAGGTGATGCAACATACGGAAAACTTACCCTTAAATTTATTTG
CACTACTGGAAAACTACCTGTTCCATGGCCAACACTTGTCACTACTTTCGCGTATGGTCTTCAATGCTTTGCGAG
ATACCCAGATCATATGAAACAGCATGACTTTTTCAAGAGTGCCATGCCCGAAGGTTATGTACAGGAAAGAACTAT
ATTTTTCAAAGATGACGGGAACTACAAGACACGTGCTGAAGTCAAGTTTGAAGGTGATACCCTTGTTAATAGAAT
CGAGTTAAAAGGTATTGATTTTAAAGAAGATGGAAACATTCTTGGACACAAATTGGAATACAACTATAACTCACA
CAATGTATACATCATGGCAGACAAACAAAAGAATGGAATCAAAGTTAACTTCAAAATTAGACACAACATTGAAGA
TGGAAGCGTTCAACTAGCAGACCATTATCAACAAAATACTCCAATTGGCGATGGCCCTGTCCTTTTACCAGACAA
CCATTACCTGTCCACACAATCTGCCCTTTCGAAAGATCCCAACGAAAAGAGAGACCACATGGTCCTTCTTGAGTT
TGTAACAGCTGCTGGGATTACACATGGCATGGATGAACTATACAAGCATGCGTGA**ctcgag**

#### pACM1-GFP (4343 bp):

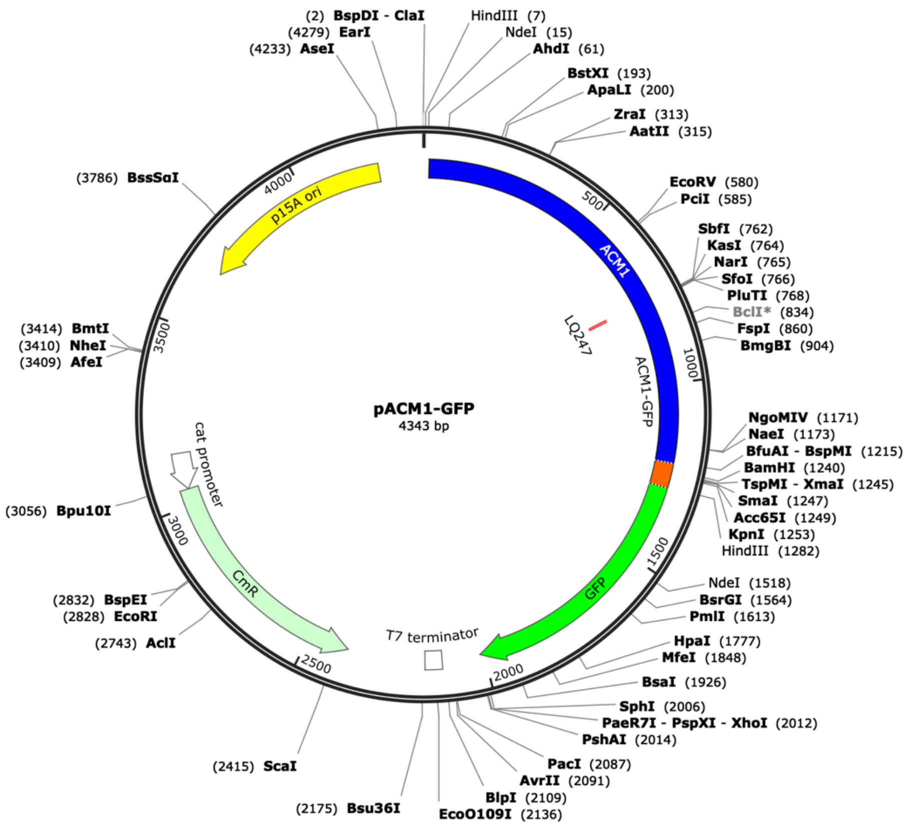


#### DNA sequence (*Cla*I-*Xho*I fragment)

**atcgat**aagcttgcatATGCAGCAATCGCATCAGGCTGGTTACGCAAACGCCGCCGACCGGGAGTCTGGCATCCC
CGCAGCCGTACTCGATGGCATCAAGGCCGTGGCGAAGGAAAAAAACGCCACATTGATGTTCCGCCTGGTCAACCC
CCATTCCACCAGCCTGATTGCCGAAGGGGTGGCCACCAAAGGATTGGGCGTGCACGCCAAGTCGTCCGATTGGGG
GTTGCAGGCGGGCTACATTCCCGTCAACCCGAATCTTTCCAAACTGTTCGGCCGTGCGCCCGAGGTGATCGCGCG
GGCCGACAACGACGTCAACAGCAGCCTGGCGCATGGCCATACCGCGGTCGACCTGACGCTGTCGAAAGAGCGGCT
TGACTATCTGCGGCAAGCGGGCCTGGTCACCGGCATGGCCGATGGCGTGGTCGCGAGCAACCACGCAGGCTACGA
GCAGTTCGAGTTTCGCGTGAAGGAAACCTCGGACGGGCGCTATGCCGTGCAGTATCGCCGCAAGGGCGGCGACGA
TTTCGAGGCGGTCAAGGTGATCGGCAATGCCGCCGGTATTCCACTGACGGCGGATATCGACATGTTCGCCATTAT
GCCGCATCTGTCCAACTTCCGCGACTCGGCGCGCAGTTCGGTGACCAGCGGCGATTCGGTGACCGATTACCTGGC
GCGCACGCGGCGGGCCGCCAGCGAGGCCACGGGCGGCCTGGATCGCGAACGCATCGACTTGTTGTGGAAAATCGC
TCGCGCC**CTGCAG**GGCGCCCGTTCCGCAGTGGGCACCGAGGCGCGTCGCCAGTTCCGCTACGACGGCGACATGAA
TATCGGCGTGATCACCGATTTCGAGCTGGAAGTGCGCAATGCGCTGAACAGGCGGGCGCACGCCGTCGGCGCGCA
GGACGTGGTCCAGCATGGCACTGAGCAGAACAATCCTTTCCCGGAGGCAGATGAGAAGATTTTCGTCGTATCGGC
CACCGGTGAAAGCCAGATGCTCACGCGCGGGCAACTGAAGGAATACATTGGCCAGCAGCGCGGCGAGGGCTATGT
CTTCTACGAGAACCGTGCATACGGCGTGGCGGGGAAAAGCCTGTTCGACGATGGGCTGGGAGCCGCGCCCGGCGT
GCCGAGCGGACGTTCGAAGTTCTCGCCGGATGTACTGGAAACGGTGCCGGCGTCACCCGGATTGCGGCGGCCGTC
GCTGGGCGCAGTGGAACGCCACTGCAGGTCGACTCTAGAGGATCCCCGGGTACCGGGGGGGTCTATGACCATGAT
TACGCCAAGCTTGATGAGTAAAGGAGAAGAACTTTTCACTGGAGTTGTCCCAATTCTTGTTGAATTAGATGGTGA
TGTTAATGGGCACAAATTTTCTGTCAGTGGAGAGGGTGAAGGTGATGCAACATACGGAAAACTTACCCTTAAATT
TATTTGCACTACTGGAAAACTACCTGTTCCATGGCCAACACTTGTCACTACTTTCGCGTATGGTCTTCAATGCTT
TGCGAGATACCCAGATCATATGAAACAGCATGACTTTTTCAAGAGTGCCATGCCCGAAGGTTATGTACAGGAAAG
AACTATATTTTTCAAAGATGACGGGAACTACAAGACACGTGCTGAAGTCAAGTTTGAAGGTGATACCCTTGTTAA
TAGAATCGAGTTAAAAGGTATTGATTTTAAAGAAGATGGAAACATTCTTGGACACAAATTGGAATACAACTATAA
CTCACACAATGTATACATCATGGCAGACAAACAAAAGAATGGAATCAAAGTTAACTTCAAAATTAGACACAACAT
TGAAGATGGAAGCGTTCAACTAGCAGACCATTATCAACAAAATACTCCAATTGGCGATGGCCCTGTCCTTTTACC
AGACAACCATTACCTGTCCACACAATCTGCCCTTTCGAAAGATCCCAACGAAAAGAGAGACCACATGGTCCTTCT
TGAGTTTGTAACAGCTGCTGGGATTACACATGGCATGGATGAACTATACAAGCATGCGTGA**ctcgag**

#### pACM2-Fkbp (3914 bp):

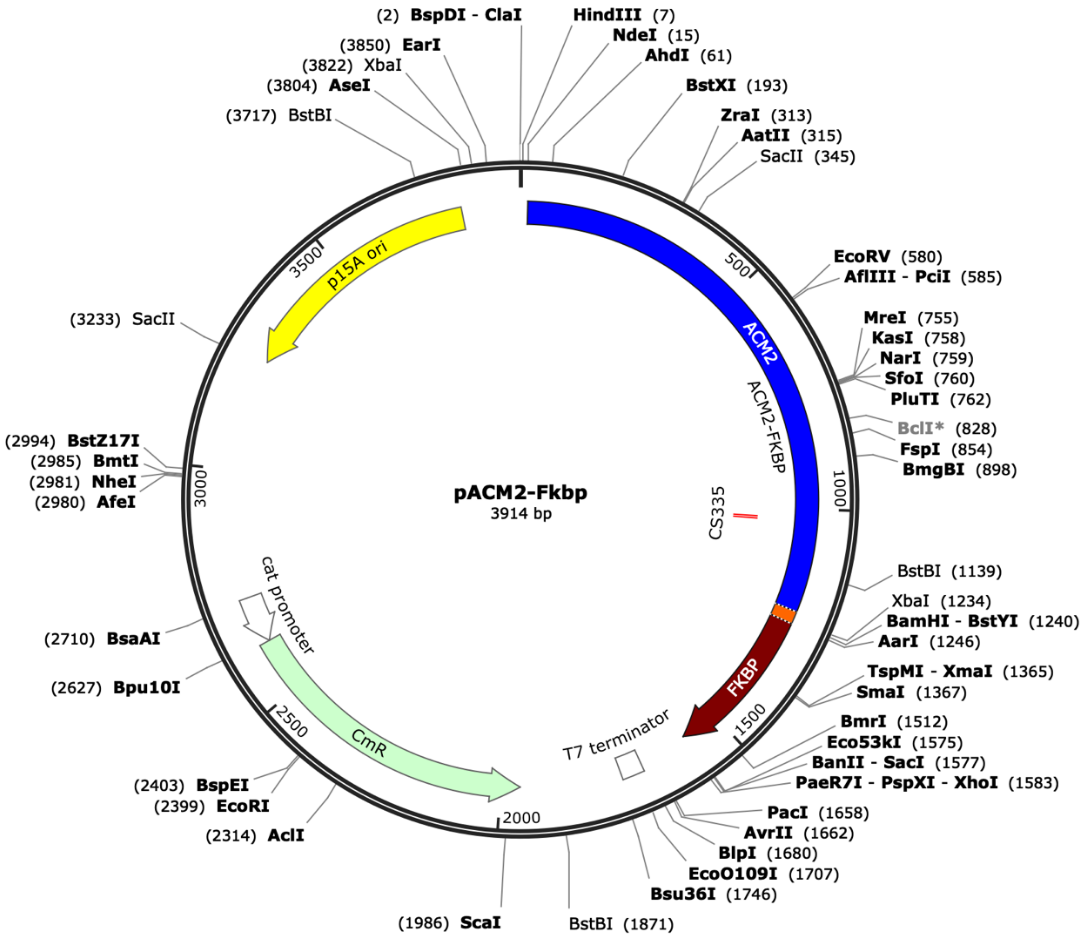


#### DNA sequence (*Cla*I-*Xho*I fragment)

**atcgat**aagcttgcatATGCAGCAATCGCATCAGGCTGGTTACGCAAACGCCGCCGACCGGGAGTCTGGCATCCC
CGCAGCCGTACTCGATGGCATCAAGGCCGTGGCGAAGGAAAAAAACGCCACATTGATGTTCCGCCTGGTCAACCC
CCATTCCACCAGCCTGATTGCCGAAGGGGTGGCCACCAAAGGATTGGGCGTGCACGCCAAGTCGTCCGATTGGGG
GTTGCAGGCGGGCTACATTCCCGTCAACCCGAATCTTTCCAAACTGTTCGGCCGTGCGCCCGAGGTGATCGCGCG
GGCCGACAACGACGTCAACAGCAGCCTGGCGCATGGCCATACCGCGGTCGACCTGACGCTGTCGAAAGAGCGGCT
TGACTATCTGCGGCAAGCGGGCCTGGTCACCGGCATGGCCGATGGCGTGGTCGCGAGCAACCACGCAGGCTACGA
GCAGTTCGAGTTTCGCGTGAAGGAAACCTCGGACGGGCGCTATGCCGTGCAGTATCGCCGCAAGGGCGGCGACGA
TTTCGAGGCGGTCAAGGTGATCGGCAATGCCGCCGGTATTCCACTGACGGCGGATATCGACATGTTCGCCATTAT
GCCGCATCTGTCCAACTTCCGCGACTCGGCGCGCAGTTCGGTGACCAGCGGCGATTCGGTGACCGATTACCTGGC
GCGCACGCGGCGGGCCGCCAGCGAGGCCACGGGCGGCCTGGATCGCGAACGCATCGACTTGTTGTGGAAAATCGC
TCGCGCCGGCGCCCGTTCCGCAGTGGGCACCGAGGCGCGTCGCCAGTTCCGCTACGACGGCGACATGAATATCGG
CGTGATCACCGATTTCGAGCTGGAAGTGCGCAATGCGCTGAACAGGCGGGCGCACGCCGTCGGCGCGCAGGACGT
GGTCCAGCATGGCACTGAGCAGAACAATCCTTTCCCGGAGGCAGATGAGAAGATTTTCGTCGTATCGGCCACCGG
TGAAAGCCAGATGCTCACGCGCGGGCAACTGAAGGAATACATTGGC**TGCAGC**CAGCAGCGCGGCGAGGGCTATGT
CTTCTACGAGAACCGTGCATACGGCGTGGCGGGGAAAAGCCTGTTCGACGATGGGCTGGGAGCCGCGCCCGGCGT
GCCGAGCGGACGTTCGAAGTTCTCGCCGGATGTACTGGAAACGGTGCCGGCGTCACCCGGATTGCGGCGGCCGTC
GCTGGGCGCAGTGGAACGCCACTGCAGGTCGACTCTAGAGGATCCCATGGGCGTGCAGGTGGAGACTATCTCCCC
AGGAGACGGGCGCACCTTCCCCAAGCGCGGCCAGACCTGCGTGGTGCACTACACCGGGATGCTTGAAGATGGAAA
GAAATTTGATTCCTCCCGGGACAGAAACAAGCCCTTTAAGTTTATGCTAGGCAAGCAGGAGGTGATCCGAGGCTG
GGAAGAAGGGGTTGCCCAGATGAGTGTGGGTCAGAGAGCCAAACTGACTATATCTCCAGATTATGCCTATGGTGC
CACTGGGCACCCAGGCATCATCCCACCACATGCCACTCTCGTCTTCGATGTGGAGCTTCTAAAACTGGAACGGAG
CTCTTAA**ctcgag**

#### pACM2-Zip ( 3743 bp):

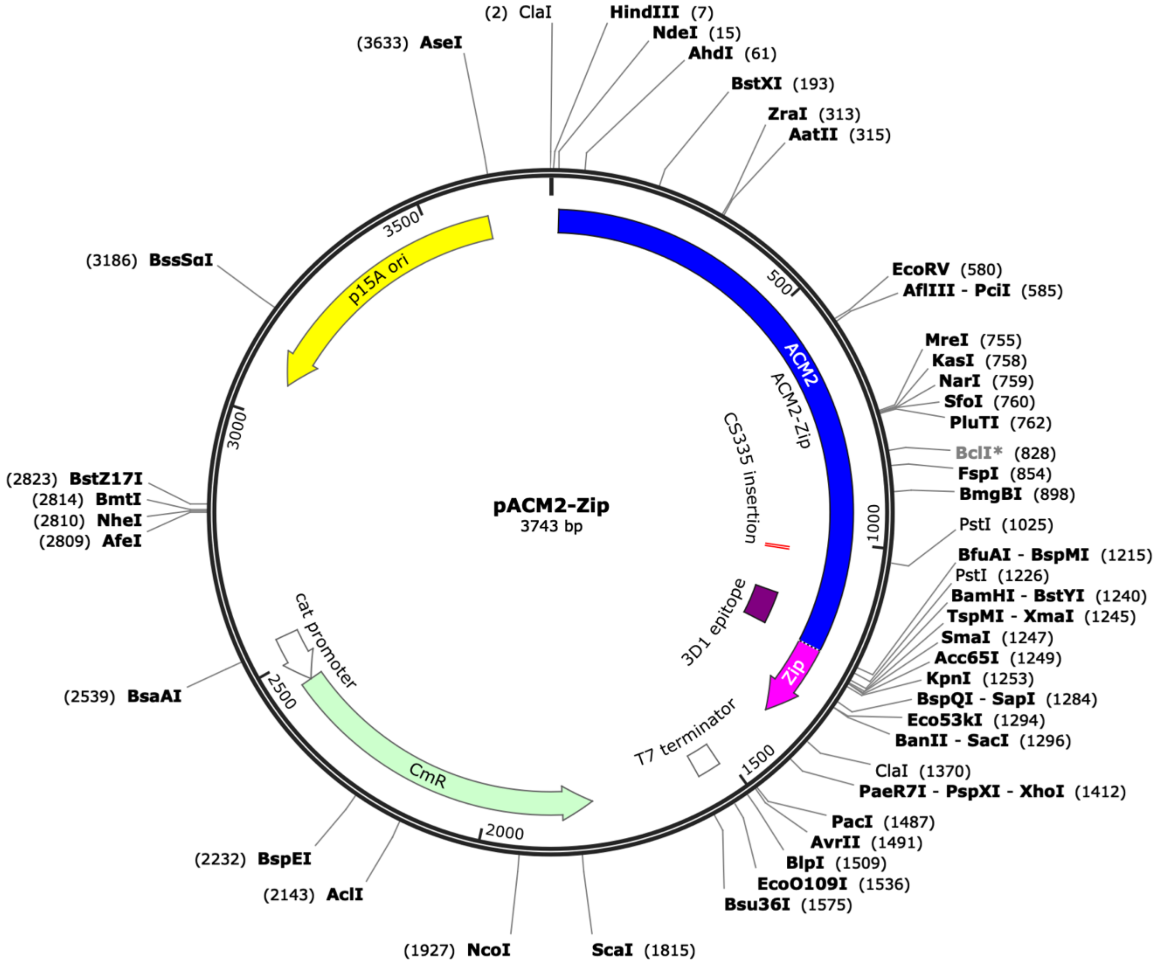


#### DNA sequence (*Cla*I-*Xho*I fragment)

**atcgat**aagcttgcatATGCAGCAATCGCATCAGGCTGGTTACGCAAACGCCGCCGACCGGGAGTCTGGCATCCC
CGCAGCCGTACTCGATGGCATCAAGGCCGTGGCGAAGGAAAAAAACGCCACATTGATGTTCCGCCTGGTCAACCC
CCATTCCACCAGCCTGATTGCCGAAGGGGTGGCCACCAAAGGATTGGGCGTGCACGCCAAGTCGTCCGATTGGGG
GTTGCAGGCGGGCTACATTCCCGTCAACCCGAATCTTTCCAAACTGTTCGGCCGTGCGCCCGAGGTGATCGCGCG
GGCCGACAACGACGTCAACAGCAGCCTGGCGCATGGCCATACCGCGGTCGACCTGACGCTGTCGAAAGAGCGGCT
TGACTATCTGCGGCAAGCGGGCCTGGTCACCGGCATGGCCGATGGCGTGGTCGCGAGCAACCACGCAGGCTACGA
GCAGTTCGAGTTTCGCGTGAAGGAAACCTCGGACGGGCGCTATGCCGTGCAGTATCGCCGCAAGGGCGGCGACGA
TTTCGAGGCGGTCAAGGTGATCGGCAATGCCGCCGGTATTCCACTGACGGCGGATATCGACATGTTCGCCATTAT
GCCGCATCTGTCCAACTTCCGCGACTCGGCGCGCAGTTCGGTGACCAGCGGCGATTCGGTGACCGATTACCTGGC
GCGCACGCGGCGGGCCGCCAGCGAGGCCACGGGCGGCCTGGATCGCGAACGCATCGACTTGTTGTGGAAAATCGC
TCGCGCCGGCGCCCGTTCCGCAGTGGGCACCGAGGCGCGTCGCCAGTTCCGCTACGACGGCGACATGAATATCGG
CGTGATCACCGATTTCGAGCTGGAAGTGCGCAATGCGCTGAACAGGCGGGCGCACGCCGTCGGCGCGCAGGACGT
GGTCCAGCATGGCACTGAGCAGAACAATCCTTTCCCGGAGGCAGATGAGAAGATTTTCGTCGTATCGGCCACCGG
TGAAAGCCAGATGCTCACGCGCGGGCAACTGAAGGAATACATTGGC**TGCAGC**CAGCAGCGCGGCGAGGGCTATGT
CTTCTACGAGAACCGTGCATACGGCGTGGCGGGGAAAAGCCTGTTCGACGATGGGCTGGGAGCCGCGCCCGGCGT
GCCGAGCGGACGTTCGAAGTTCTCGCCGGATGTACTGGAAACGGTGCCGGCGTCACCCGGATTGCGGCGGCCGTC
GCTGGGCGCAGTGGAACGCCACTGCAGGTCGACTCTAGAGGATCCCCGGGTACCTATCCAGCGTATGAAACAGCT
GGAAGACAAAGTTGAAGAGCTCCTGAGCAAAAACTACCACCTGGAGAACGAAGTTGCGCGCCTGAAAAAACTGGT
GGGTGAACGTGGGAATTCATCGATATAActaagtaatatggtgcactctcagtacaatctg**ctcgag**

#### pACM2-TM-Zip ( 3869 bp):

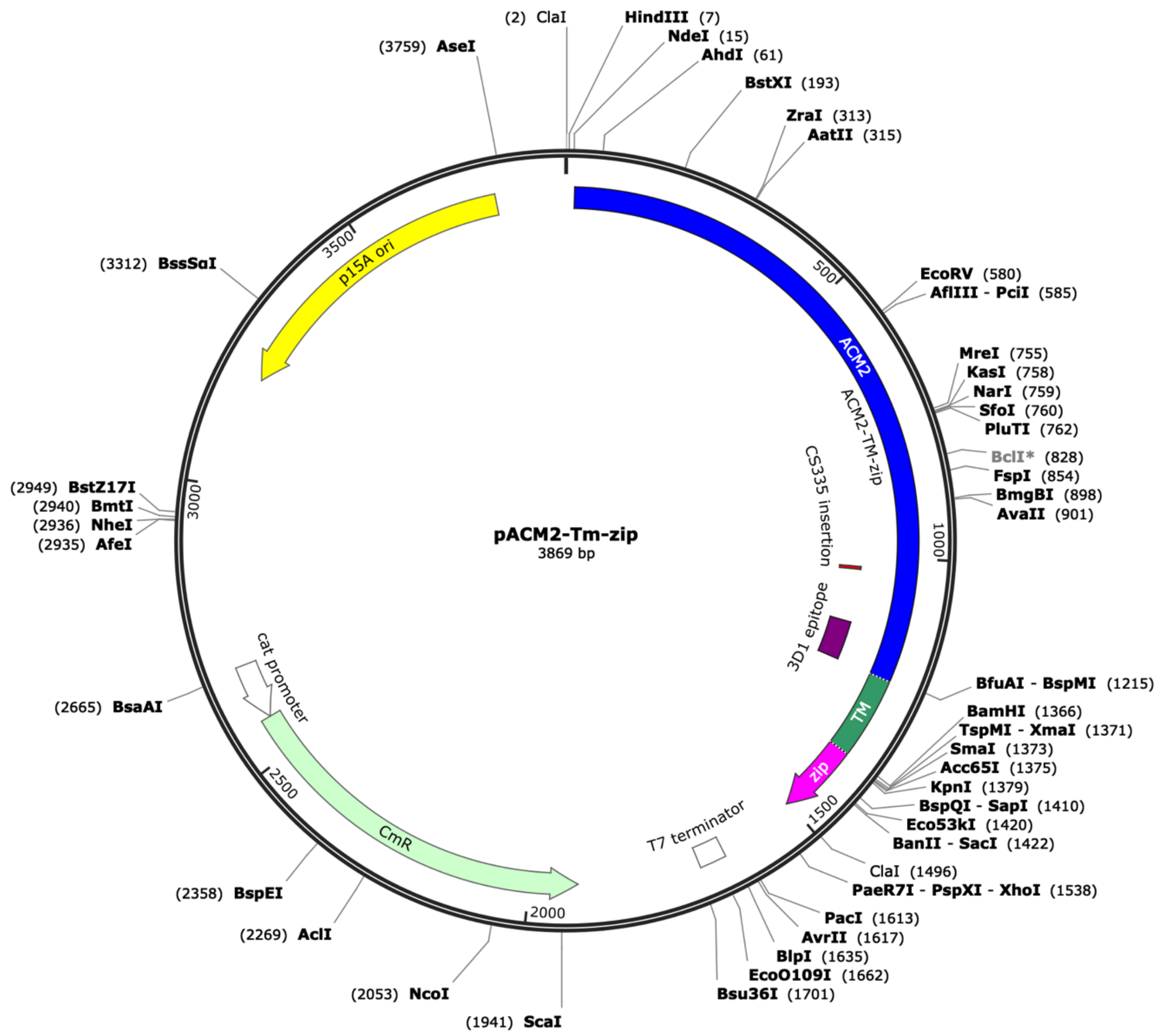


#### DNA sequence (*Cla*I-*Xho*I fragment)

**atcgat**aagcttgcatATGCAGCAATCGCATCAGGCTGGTTACGCAAACGCCGCCGACCGGGAGTCTGGCATCCC
CGCAGCCGTACTCGATGGCATCAAGGCCGTGGCGAAGGAAAAAAACGCCACATTGATGTTCCGCCTGGTCAACCC
CCATTCCACCAGCCTGATTGCCGAAGGGGTGGCCACCAAAGGATTGGGCGTGCACGCCAAGTCGTCCGATTGGGG
GTTGCAGGCGGGCTACATTCCCGTCAACCCGAATCTTTCCAAACTGTTCGGCCGTGCGCCCGAGGTGATCGCGCG
GGCCGACAACGACGTCAACAGCAGCCTGGCGCATGGCCATACCGCGGTCGACCTGACGCTGTCGAAAGAGCGGCT
TGACTATCTGCGGCAAGCGGGCCTGGTCACCGGCATGGCCGATGGCGTGGTCGCGAGCAACCACGCAGGCTACGA
GCAGTTCGAGTTTCGCGTGAAGGAAACCTCGGACGGGCGCTATGCCGTGCAGTATCGCCGCAAGGGCGGCGACGA
TTTCGAGGCGGTCAAGGTGATCGGCAATGCCGCCGGTATTCCACTGACGGCGGATATCGACATGTTCGCCATTAT
GCCGCATCTGTCCAACTTCCGCGACTCGGCGCGCAGTTCGGTGACCAGCGGCGATTCGGTGACCGATTACCTGGC
GCGCACGCGGCGGGCCGCCAGCGAGGCCACGGGCGGCCTGGATCGCGAACGCATCGACTTGTTGTGGAAAATCGC
TCGCGCCGGCGCCCGTTCCGCAGTGGGCACCGAGGCGCGTCGCCAGTTCCGCTACGACGGCGACATGAATATCGG
CGTGATCACCGATTTCGAGCTGGAAGTGCGCAATGCGCTGAACAGGCGGGCGCACGCCGTCGGCGCGCAGGACGT
GGTCCAGCATGGCACTGAGCAGAACAATCCTTTCCCGGAGGCAGATGAGAAGATTTTCGTCGTATCGGCCACCGG
TGAAAGCCAGATGCTCACGCGCGGGCAACTGAAGGAATACATTGGC**TGCAGC**CAGCAGCGCGGCGAGGGCTATGT
CTTCTACGAGAACCGTGCATACGGCGTGGCGGGAAAAGCCTGTTCGACGATGGGCTGGGAGCCGCGCCCGGCGT
GCCGAGCGGACGTTCGAAGTTCTCGCCGGATGTACTGGAAACGGTGCCGGCGTCACCCGGATTGCGGCGGCCGTC
GCTGGGCGCAGTGGAACGCCACTGCAGGTCGACTCTAGAGGATCTAAAATTTATTCTACGTCGCTGTCTGGAAGC
GATTCCGACGCTATTTATTCTTATTACTATTTCGTTCTTTATGATGCGCCTCGCGCCGGGAAGCCCTTTTACCGG
CGAACGTACTTTAGCGGATCCCCGGGTACCTATCCAGCGTATGAAACAGCTGGAAGACAAAGTTGAAGAGCTCCT
GAGCAAAAACTACCACCTGGAGAACGAAGTTGCGCGCCTGAAAAAACTGGTGGGTGAACGTGGGAATTCATCGAT
ATAActaagtaatatggtgcactctcagtacaatctg**ctcgag**

#### pACM2-Barnase ( 3960 bp):

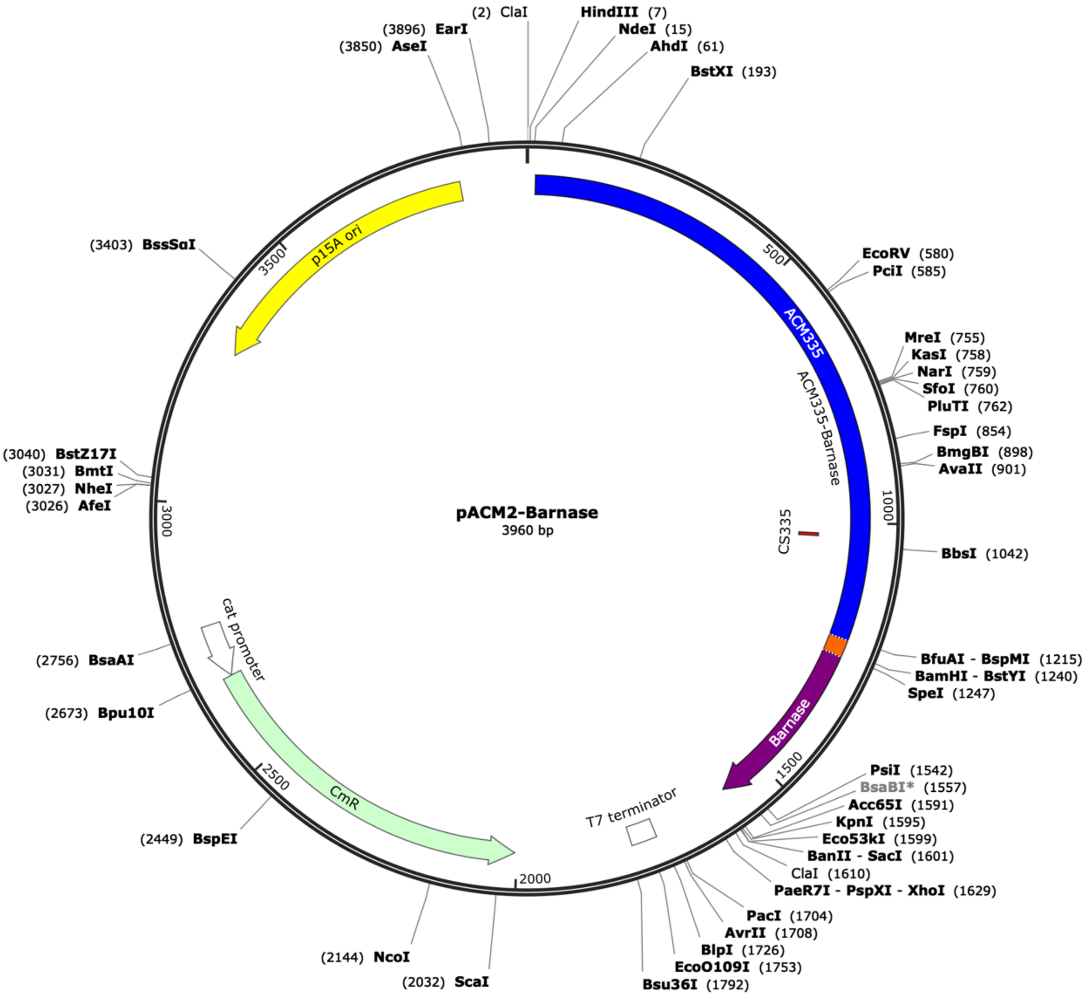


#### DNA sequence (*Cla*I-*Xho*I fragment)

**atcgat**aagcttgcatATGCAGCAATCGCATCAGGCTGGTTACGCAAACGCCGCCGACCGGGAGTCTGGCATCCC
CGCAGCCGTACTCGATGGCATCAAGGCCGTGGCGAAGGAAAAAAACGCCACATTGATGTTCCGCCTGGTCAACCC
CCATTCCACCAGCCTGATTGCCGAAGGGGTGGCCACCAAAGGATTGGGCGTGCACGCCAAGTCGTCCGATTGGGG
GTTGCAGGCGGGCTACATTCCCGTCAACCCGAATCTTTCCAAACTGTTCGGCCGTGCGCCCGAGGTGATCGCGCG
GGCCGACAACGACGTCAACAGCAGCCTGGCGCATGGCCATACCGCGGTCGACCTGACGCTGTCGAAAGAGCGGCT
TGACTATCTGCGGCAAGCGGGCCTGGTCACCGGCATGGCCGATGGCGTGGTCGCGAGCAACCACGCAGGCTACGA
GCAGTTCGAGTTTCGCGTGAAGGAAACCTCGGACGGGCGCTATGCCGTGCAGTATCGCCGCAAGGGCGGCGACGA
TTTCGAGGCGGTCAAGGTGATCGGCAATGCCGCCGGTATTCCACTGACGGCGGATATCGACATGTTCGCCATTAT
GCCGCATCTGTCCAACTTCCGCGACTCGGCGCGCAGTTCGGTGACCAGCGGCGATTCGGTGACCGATTACCTGGC
GCGCACGCGGCGGGCCGCCAGCGAGGCCACGGGCGGCCTGGATCGCGAACGCATCGACTTGTTGTGGAAAATCGC
TCGCGCCGGCGCCCGTTCCGCAGTGGGCACCGAGGCGCGTCGCCAGTTCCGCTACGACGGCGACATGAATATCGG
CGTGATCACCGATTTCGAGCTGGAAGTGCGCAATGCGCTGAACAGGCGGGCGCACGCCGTCGGCGCGCAGGACGT
GGTCCAGCATGGCACTGAGCAGAACAATCCTTTCCCGGAGGCAGATGAGAAGATTTTCGTCGTATCGGCCACCGG
TGAAAGCCAGATGCTCACGCGCGGGCAACTGAAGGAATACATTGGC**TGCAGC**CAGCAGCGCGGCGAGGGCTATGT
CTTCTACGAGAACCGTGCATACGGCGTGGCGGGGAAAAGCCTGTTCGACGATGGGCTGGGAGCCGCGCCCGGCGT
GCCGAGCGGACGTTCGAAGTTCTCGCCGGATGTACTGGAAACGGTGCCGGCGTCACCCGGATTGCGGCGGCCGTC
GCTGGGCGCAGTGGAACGCCACTGCAGGTCGACTCTAGAGGATCCAACTAGTGCACAGGTTATCAACACGTTTGA
CGGGGTTGCGGATTATCTTCAGACATATCATAAGCTACCTGATAATTACATTACAAAATCAGAAGCACAAGCCCT
CGGCTGGGTGGCATCAAAAGGGAACCTTGCAGACGTCGCTCCGGGGAAAAGCATCGGCGGAGACATCTTCTCAAA
CAGGGAAGGCAAACTCCCGGGCAAAAGCGGACGAACATGGCGTGAAGCGGATATTAACTATACATCAGGCTTCAG
AAATTCAGACCGGATTCTTTACTCAAGCGACTGGCTGATTTATAAAACAACTGATCATTATCAAACGTTTACAAA
AATCAGATAAccccgggtaccgagctcgaattcatcgatataactaagtaata**ctcgag**

#### pTrACM2-Gfp ( 5264 bp):

#### Map

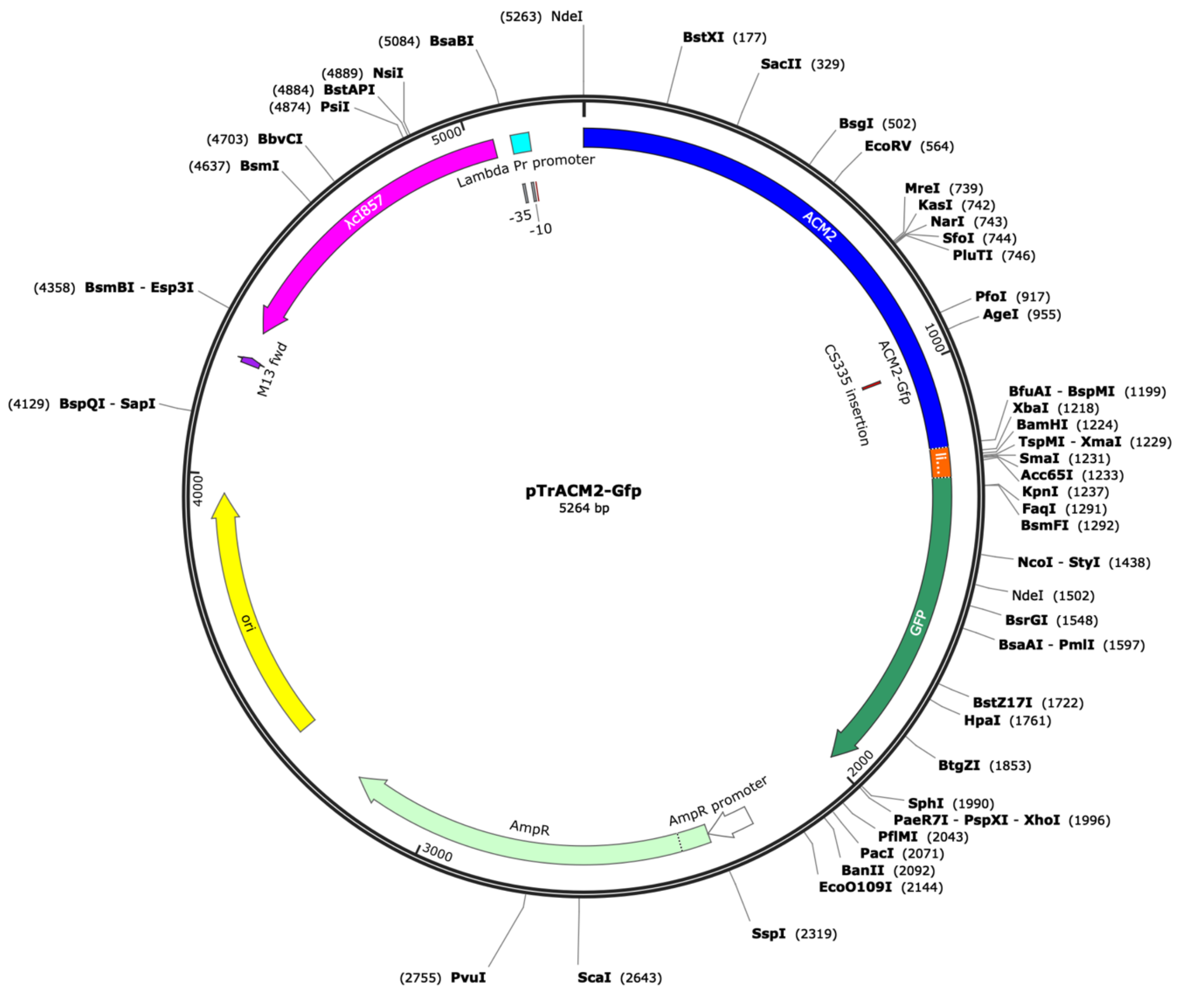


#### DNA sequence

ATGCAGCAATCGCATCAGGCTGGTTACGCAAACGCCGCCGACCGGGAGTCTGGCATCCCCGCAGCCGTACTCGAT
GGCATCAAGGCCGTGGCGAAGGAAAAAAACGCCACATTGATGTTCCGCCTGGTCAACCCCCATTCCACCAGCCTG
ATTGCCGAAGGGGTGGCCACCAAAGGATTGGGCGTGCACGCCAAGTCGTCCGATTGGGGGTTGCAGGCGGGCTAC
ATTCCCGTCAACCCGAATCTTTCCAAACTGTTCGGCCGTGCGCCCGAGGTGATCGCGCGGGCCGACAACGACGTC
AACAGCAGCCTGGCGCATGGCCATACCGCGGTCGACCTGACGCTGTCGAAAGAGCGGCTTGACTATCTGCGGCAA
GCGGGCCTGGTCACCGGCATGGCCGATGGCGTGGTCGCGAGCAACCACGCAGGCTACGAGCAGTTCGAGTTTCGC
GTGAAGGAAACCTCGGACGGGCGCTATGCCGTGCAGTATCGCCGCAAGGGCGGCGACGATTTCGAGGCGGTCAAG
GTGATCGGCAATGCCGCCGGTATTCCACTGACGGCGGATATCGACATGTTCGCCATTATGCCGCATCTGTCCAAC
TTCCGCGACTCGGCGCGCAGTTCGGTGACCAGCGGCGATTCGGTGACCGATTACCTGGCGCGCACGCGGCGGGCC
GCCAGCGAGGCCACGGGCGGCCTGGATCGCGAACGCATCGACTTGTTGTGGAAAATCGCTCGCGCCGGCGCCCGT
TCCGCAGTGGGCACCGAGGCGCGTCGCCAGTTCCGCTACGACGGCGACATGAATATCGGCGTGATCACCGATTTC
GAGCTGGAAGTGCGCAATGCGCTGAACAGGCGGGCGCACGCCGTCGGCGCGCAGGACGTGGTCCAGCATGGCACT
GAGCAGAACAATCCTTTCCCGGAGGCAGATGAGAAGATTTTCGTCGTATCGGCCACCGGTGAAAGCCAGATGCTC
ACGCGCGGGCAACTGAAGGAATACATTGGC**TGCAGC**CAGCAGCGCGGCGAGGGCTATGTCTTCTACGAGAACCGT
GCATACGGCGTGGCGGGGAAAAGCCTGTTCGACGATGGGCTGGGAGCCGCGCCCGGCGTGCCGAGCGGACGTTCG
AAGTTCTCGCCGGATGTACTGGAAACGGTGCCGGCGTCACCCGGATTGCGGCGGCCGTCGCTGGGCGCAGTGGAA
CGCCACTGCAGGTCGACTCTAGAGGATCCCCGGGTACCGGGGGGGTCTATGACCATGATTACGCCAAGCTTGATG
AGTAAAGGAGAAGAACTTTTCACTGGAGTTGTCCCAATTCTTGTTGAATTAGATGGTGATGTTAATGGGCACAAA
TTTTCTGTCAGTGGAGAGGGTGAAGGTGATGCAACATACGGAAAACTTACCCTTAAATTTATTTGCACTACTGGA

AAACTACCTGTTCCATGGCCAACACTTGTCACTACTTTCGCGTATGGTCTTCAATGCTTTGCGAGATACCCAGAT
CATATGAAACAGCATGACTTTTTCAAGAGTGCCATGCCCGAAGGTTATGTACAGGAAAGAACTATATTTTTCAAA
GATGACGGGAACTACAAGACACGTGCTGAAGTCAAGTTTGAAGGTGATACCCTTGTTAATAGAATCGAGTTAAAA
GGTATTGATTTTAAAGAAGATGGAAACATTCTTGGACACAAATTGGAATACAACTATAACTCACACAATGTATAC
ATCATGGCAGACAAACAAAAGAATGGAATCAAAGTTAACTTCAAAATTAGACACAACATTGAAGATGGAAGCGTT
CAACTAGCAGACCATTATCAACAAAATACTCCAATTGGCGATGGCCCTGTCCTTTTACCAGACAACCATTACCTG
TCCACACAATCTGCCCTTTCGAAAGATCCCAACGAAAAGAGAGACCACATGGTCCTTCTTGAGTTTGTAACAGCT
GCTGGGATTACACATGGCATGGATGAACTATACAAGCATGCGTGActcgagtctggtaaagaaaccgctgctgcg
aaatttgaacgccagcacatggactcgtctactagcgcagcttaattaacctagagtcgaccgagcccgcctaat
gagcgggcttttttttcatgcaagctaattcttgaagacgaaagggcctcgtgatacgcctatttttataggtta
atgtcatgataataatggtttcttagacgtcaggtggcacttttcggggaaatgtgcgcggaacccctatttgtt
tatttttctaaatacattcaaatatgtatccgctcatgagacaataaccctgataaatgcttcaataatattgaa
aaaggaagagtatgagtattcaacatttccgtgtcgcccttattcccttttttgcggcattttgccttcctgttt
ttgctcacccagaaacgctggtgaaagtaaaagatgctgaagatcagttgggtgcacgagtgggttacatcgaac
tggatctcaacagcggtaagatccttgagagttttcgccccgaagaacgttttccaatgatgagcacttttaaag
ttctgctatgtggcgcggtattatcccgtattgacgtcgggcaagagcaactcggtcgccgcatacactattctc
agaatgacttggttgagtactcaccagtcacagaaaagcatcttacggatggcatgacagtaagagaattatgca
gtgctgccataaccatgagtgataacactgcggccaacttacttctgacaacgatcggaggaccgaaggagctaa
ccgcttttttgcacaacatgggggatcaagtaactcgccttgatcgttgggaaccggagctgaatgaagccatac
caaacgacgagcgtgacaccacgatgcctgtagcaatggcaacaacgttgcgcaaactattaactggcgaactac
ttactctagcttcccggcaacaattaatagactggatggaggcggataaagttgcaggaccacttctgcgctcgg
cccttccggctggctggtttattgctgataaatctggagccggtgagcgtgggtctcgcggtatcattgcagcac
tggggccagatggtaagccctcccgtatcgtagttatctacacgacggggagtcaggcaactatggatgaacgaa
atagacagatcgctgagataggtgcctcactgattaagcattggtaactgtcagaccaagtttactcatatatac
tttagattgatttaaaacttcatttttaatttaaaaggatctaggtgaagatcctttttgataatctcatgacca
aaatcccttaacgtgagttttcgttccactgagcgtcagaccccgtagaaaagatcaaaggatcttcttgagatc
ctttttttctgcgcgtaatctgctgcttgcaaacaaaaaaaccaccgctaccagcggtggtttgtttgccggatc
aagagctaccaactctttttccgaaggtaactggcttcagcagagcgcagataccaaatactgtccttctagtgt
agccgtagttaggccaccacttcaagaactctgtagcaccgcctacatacctcgctctgctaatcctgttaccag
tggctgctgccagtggcgataagtcgtgtcttaccgggttggactcaagacgatagttaccggataaggcgcagc
ggtcgggctgaacggggggttcgtgcacacagcccagcttggagcgaacgacctacaccgaactgagatacctac
agcgtgagcattgagaaagcgccacgcttcccgaagggagaaaggcggacaggtatccggtaagcggcagggtcg
gaacaggagagcgcacgagggagcttccagggggaaacgcctggtatctttatagtcctgtcgggtttcgccacc
tctgacttgagcgtcgatttttgtgatgctcgtcaggggggcggagcctatggaaaaacgccagcaacgcggcct
ttttacggttcctggccttttgctggccttttgctcacatgttctctcctgcgttatcccctgattctgtggata
accgtattaccgcctttgagtgagctgataccgctcgccgcagccgaacgaccgagcgcagcgagtcagtgagcg
aggaagcggaagagcgcccaatacgcaaaccgcctctccccgcgcgttggccgattcattaatgcagctggcgaa
agggggatgtgctgcaaggcgattaagttgggtaacgccagggttttcccagtcacgacgttgtaaaacgacggc
cagtgccaagcttgaagattcttgctcaattgttatcagctatgcgccgaccagaacaccttgccgatcagccaa
acgtctcttcaggccactgactagcgataactttccccacaacggaacaactctcattgcatgggatcattgggt
actgtgggtttagtggttgtaaaaacacctgaccgctatccctgatcagtttcttgaaggtaaactcatcacccc
caagtctggctatgcagaaatcacctggctcaacagcctgctcagggtcaacgagaattaacattccgtcaggaa
agcttggcttggagcctgttggtgcggtcatggaattaccttcaacctcaagccagaatgcagaatcactggctt
ttttggttgtgcttacccatctctccgcatcacctttggtaaaggttctaagctgaggtgagaacatccctgcct
gaacatgagaaaaaacagggtactcatactcacttctaagtgacggctgcatactaaccgcttcatacatctcgt
agatttctctggcgattgaagggctaaattcttcaacgctaactttgagaatttttgtaagcaatgcggcgttat
aagcatttaatgcattgatgccattaaataaagcaccaacgcctgactgccccatccccatcttgtctacgacag
attcctgggataagccaagttcatttttctttttttcataaattgctttaaggcgacgtgcgtcctcaagctgct
cttgtgttaatggtttctttttttgtgctcatacgttaaatctatcaccgcaagggataaatatctaacaccgtgc
gtgttgactattttacctctggcggtgataatggttgcatgtactaaggaggttgtatggaacaacgcataaccc
tgaaagattatgcaatgcgctttgggcaaaccaagacagctaaagatcctagaaataattttgtttaactttaag
aaggagatatacat

#### pTCam-V$_{9A}$ ( 4544 bp)

#### Map

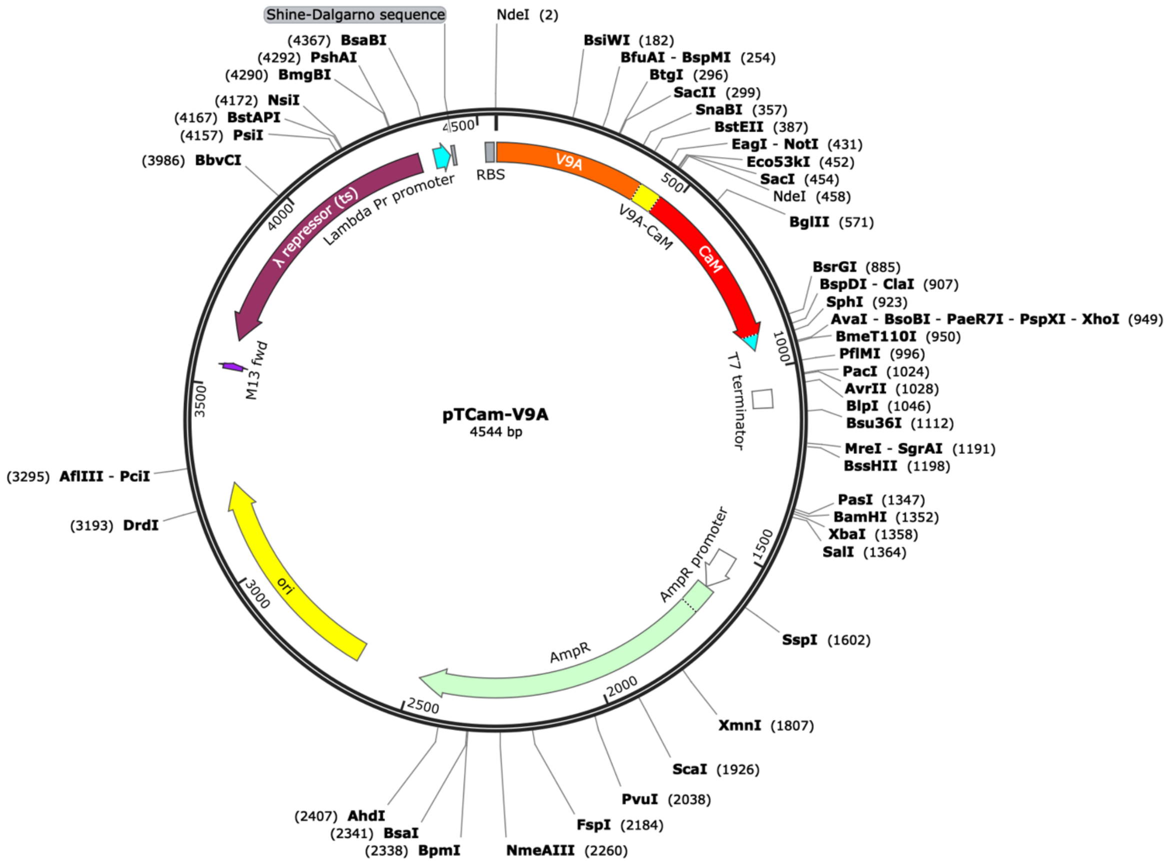


#### Full DNA sequence

**cat**ATGGCTGACGTTCAGCTGCAGGAATCTGGTGGTGGTTCTGTTCAGGCGGGTGGTTCTCTGCGTCTGTCTTGC
GCGGCTAGCGGTGACACCTTCTCTTCTTACTCTATGGCGTGGTTCCGTCAGGCGCCGGGTAAAGAATGCGAACTG
GTTTCTAACATCCTGCGTGACGGTACTACCACGTACGCCGGCTCTGTTAAAGGTCGTTTCACCATCTCTCGTGAC
GACGCGAAAAACACCGTTTACCTGCAGATGGTTAACCTGAAATCTGAAGACACCGCGCGTTACTACTGCGCCGCG
GACTCTGGTACTCAGCTGGGTTACGTTGGTGCGGTTGGTCTGTCTTGCCTGGACTACGTAATGGACTACTGGGGT
AAAGGTACTCAGGTTACCGTTTCTTCTGAACCGAAAACCCCGAAACCGCAGCCAGCGGCCGCTGAAAAGGTACCG
AGCTCCCATATGGCTGACCAACTGACAGAAGAGCAGATTGCAGAATTCAAAGAAGCTTTTTCACTATTTGACAAA
GATGGTGATGGAACTATAACAACAAAGGAATTGGGAACTGTAATGAGATCTCTTGGGCAGAATCCCACAGAAGCA
GAGTTACAGGACATGATTAATGAAGTAGATGCTGATGGTAATGGCACAATTGACTTCCCTGAATTTCTGACAATG
ATGGCAAGAAAAATGAAAGACACAGACAGTGAAGAAGAAATTAGAGAAGCATTCCGTGTGTTTGATAAGGATGGC
AATGGCTATATTAGTGCTGCAGAACTTCGCCATGTGATGACAAACCTTGGAGAGAAGTTAACAGATGAAGAAGTT
GATGAAATGATCAGGGAAGCAGATATTGATGGTGATGGTCAAGTAAACTATGAAGAGTTTGTACAAATGATGACA
GCGAAATCGATGGTACCCGCATGCTGTCCGGGTTGCTGTGCTAGCTAA**ctcgag**tctggtaaagaaaccgctgct
gcgaaatttgaacgccagcacatggactcgtctactagcgcagcttaattaacctaggctgctgccaccgctgag
caataactagcataaccccttggggcctctaaacgggtcttgaggggttttttgctgaaacctcaggcatttgag
aagcacacggtcacactgcttccggtagtcaataaaccggaccccggcgcggcggcggctgccccgccggcggcg
cgcgtgccggacacgctgatgcagtccctggctgtcaactggcgctgaagcgccgtgaatcacggcccgcctgcc
tcgcgcggcggcgccgtctctttgcgttcttctccgaggtatttcccatcatgaattcttttgcttttttaccctg
gggatcctctagagtcgaccgagcccgcctaatgagcgggcttttttttcatgcaagctaattcttgaagacgaa
agggcctcgtgatacgcctatttttataggttaatgtcatgataataatggtttcttagacgtcaggtggcactt
ttcggggaaatgtgcgcggaacccctatttgtttatttttctaaatacattcaaatatgtatccgctcatgagac
aataaccctgataaatgcttcaataatattgaaaaaggaagagtatgagtattcaacatttccgtgtcgccctta

ttcccttttttgcggcattttgccttcctgtttttgctcacccagaaacgctggtgaaagtaaaagatgctgaag
atcagttgggtgcacgagtgggttacatcgaactggatctcaacagcggtaagatccttgagagttttcgccccg
aagaacgttttccaatgatgagcacttttaaagttctgctatgtggcgcggtattatcccgtattgacgtcgggc
aagagcaactcggtcgccgcatacactattctcagaatgacttggttgagtactcaccagtcacagaaaagcatc
ttacggatggcatgacagtaagagaattatgcagtgctgccataaccatgagtgataacactgcggccaacttac
ttctgacaacgatcggaggaccgaaggagctaaccgcttttttgcacaacatgggggatcaagtaactcgccttg
atcgttgggaaccggagctgaatgaagccataccaaacgacgagcgtgacaccacgatgcctgtagcaatggcaa
caacgttgcgcaaactattaactggcgaactacttactctagcttcccggcaacaattaatagactggatggagg
cggataaagttgcaggaccacttctgcgctcggcccttccggctggctggtttattgctgataaatctggagccg
gtgagcgtgggtctcgcggtatcattgcagcactggggccagatggtaagccctcccgtatcgtagttatctaca
cgacggggagtcaggcaactatggatgaacgaaatagacagatcgctgagataggtgcctcactgattaagcatt
ggtaactgtcagaccaagtttactcatatatactttagattgatttaaaacttcatttttaatttaaaaggatct
aggtgaagatcctttttgataatctcatgaccaaaatcccttaacgtgagttttcgttccactgagcgtcagacc
ccgtagaaaagatcaaaggatcttcttgagatcctttttttctgcgcgtaatctgctgcttgcaaacaaaaaaac
caccgctaccagcggtggtttgtttgccggatcaagagctaccaactctttttccgaaggtaactggcttcagca
gagcgcagataccaaatactgtccttctagtgtagccgtagttaggccaccacttcaagaactctgtagcaccgc
ctacatacctcgctctgctaatcctgttaccagtggctgctgccagtggcgataagtcgtgtcttaccgggttgg
actcaagacgatagttaccggataaggcgcagcggtcgggctgaacggggggttcgtgcacacagcccagcttgg
agcgaacgacctacaccgaactgagatacctacagcgtgagcattgagaaagcgccacgcttcccgaagggagaa
aggcggacaggtatccggtaagcggcagggtcggaacaggagagcgcacgagggagcttccagggggaaacgcct
ggtatctttatagtcctgtcgggtttcgccacctctgacttgagcgtcgatttttgtgatgctcgtcaggggggc
ggagcctatggaaaaacgccagcaacgcggcctttttacggttcctggccttttgctggccttttgctcacatgt
tctcctgcgttatcccctgattctgtggataaccgtattaccgcctttgagtgagctgataccgctcgccgca
gccgaacgaccgagcgcagcgagtcagtgagcgaggaagcggaagagcgcccaatacgcaaaccgcctctccccg
cgcgttggccgattcattaatgcagctggcgaaagggggatgtgctgcaaggcgattaagttgggtaacgccagg
gttttcccagtcacgacgttgtaaaacgacggccagtgccaagcttgaagattcttgctcaattgttatcagcta
tgcgccgaccagaacaccttgccgatcagccaaacgtctcttcaggccactgactagcgataactttccccacaa
cggaacaactctcattgcatgggatcattgggtactgtgggtttagtggttgtaaaaacacctgaccgctatccc
tgatcagtttcttgaaggtaaactcatcacccccaagtctggctatgcagaaatcacctggctcaacagcctgct
cagggtcaacgagaattaacattccgtcaggaaagcttggcttggagcctgttggtgcggtcatggaattacctt
caacctcaagccagaatgcagaatcactggctttttttggttgtgcttacccatctctccgcatcacctttggtaa
aggttctaagctgaggtgagaacatccctgcctgaacatgagaaaaaacagggtactcatactcacttctaagtg
acggctgcatactaaccgcttcatacatctcgtagatttctctggcgattgaagggctaaattcttcaacgctaa
ctttgagaatttttgtaagcaatgcggcgttataagcatttaatgcattgatgccattaaataaagcaccaacgc
ctgactgccccatccccatcttgtctacgacagattcctgggataagccaagttcatttttctttttttcataaa
ttgctttaaggcgacgtgcgtcctcaagctgctcttgtgttaatggtttcttttttgtgctcatacgttaaatct
atcaccgcaagggataaatatctaacaccgtgcgtgttgactattttacctctggcggtgataatggttgcatgt
actaaggaggttgtatggaacaacgcataaccctgaaagattatgcaatgcgctttgggcaaaccaagacagcta
aagatcctagaaataattttgtttaactttaagaaggagatata

#### pTCam-V$_{9A}$-Zip ( 4778 bp)

##### Map

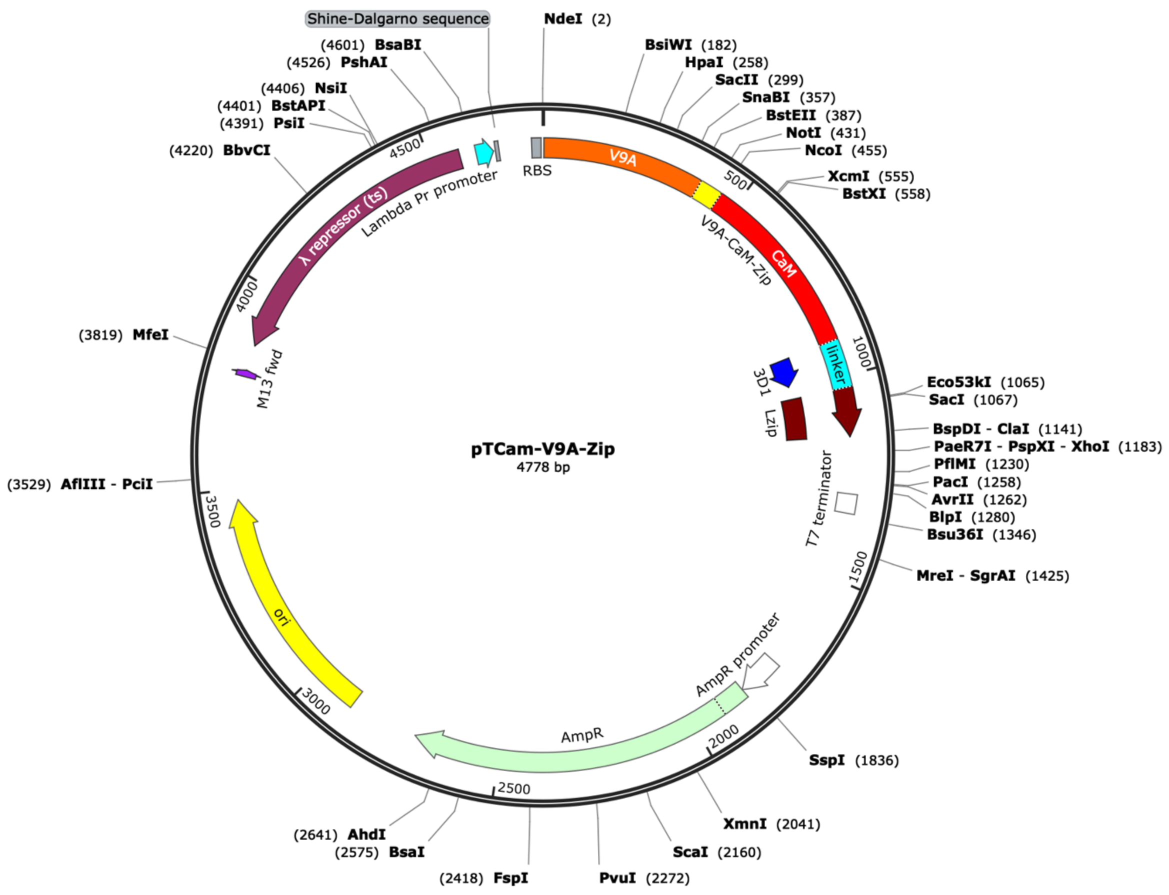


##### DNA sequence (*Nde*I-*Xho*I fragment)

**catATG**GCTGACGTTCAGCTGCAGGAATCTGGTGGTGGTTCTGTTCAGGCGGGTGGTTCTCTGCGTCTGTCTTGC
GCGGCTAGCGGTGACACCTTCTCTTCTTACTCTATGGCGTGGTTCCGTCAGGCGCCGGGTAAAGAATGCGAACTG
GTTTCTAACATCCTGCGTGACGGTACTACCACGTACGCCGGCTCTGTTAAAGGTCGTTTCACCATCTCTCGTGAC
GACGCGAAAAACACCGTTTACCTGCAGATGGTTAACCTGAAATCTGAAGACACCGCGCGTTACTACTGCGCCGCG
GACTCTGGTACTCAGCTGGGTTACGTTGGTGCGGTTGGTCTGTCTTGCCTGGACTACGTAATGGACTACTGGGGT
AAAGGTACTCAGGTTACCGTTTCTTCTGAACCGAAAACCCCGAAACCGCAGCCAGCGGCCGCTGAAAAGGTACCC
GGGTCCATGGCTAGCGCCGACCAGCTGACCGAGGAGCAGATCGCCGAGTTCAAGGAGGCCTTCTCCCTGTTCGAC
AAGGACGGCGACGGCACCATCACCACCAAGGAGCTGGGCACCGTCATGCGGTCCCTGGGCCAGAACCCCACCGAG
GCCGAGCTTCAGGACATGATCAACGAGGTCGACGCCGACGGCAACGGCACCATCGACTTCCCCGAGTTCCTGACC
ATGATGGCCCGGAAGATGAAGGACACCGACTCCGAGGAGGAGATCCGGGAGGCCTTCCGGGTCTTCGACAAGGAC
GGCAACGGCTATATCTCCGCCGCCGAGCTGCGGCACGTCATGACCAACCTGGGCGAGAAGCTGACCGACGAGGAG
GTCGACGAGATGATCCGGGAGGCCGACATCGACGGCGACGGCCAGGTCAACTATGAGGAGTTCGTCCAGATGATG
ACCGCCAAATCGAAGTTCTCGCCGGATGTACTGGAAACGGTGCCGGCGTCACCCGGATTGCGGCGGCCGTCGCTG
GGCGCAGTGGAACGCCACTGCAGGTCGACTCTAGAGGATCCCCGGGTACCTATCCAGCGTATGAAACAGCTGGAA
GACAAAGTTGAAGAGCTCCTGAGCAAAAACTACCACCTGGAGAACGAAGTTGCGCGCCTGAAAAAACTGGTGGGT
GAACGTGGGAATTCATCGATATAActaagtaatatggtgcactctcagtacaatctg**ctcgag**

### pTCam-Frb (4384 bp)

#### Map

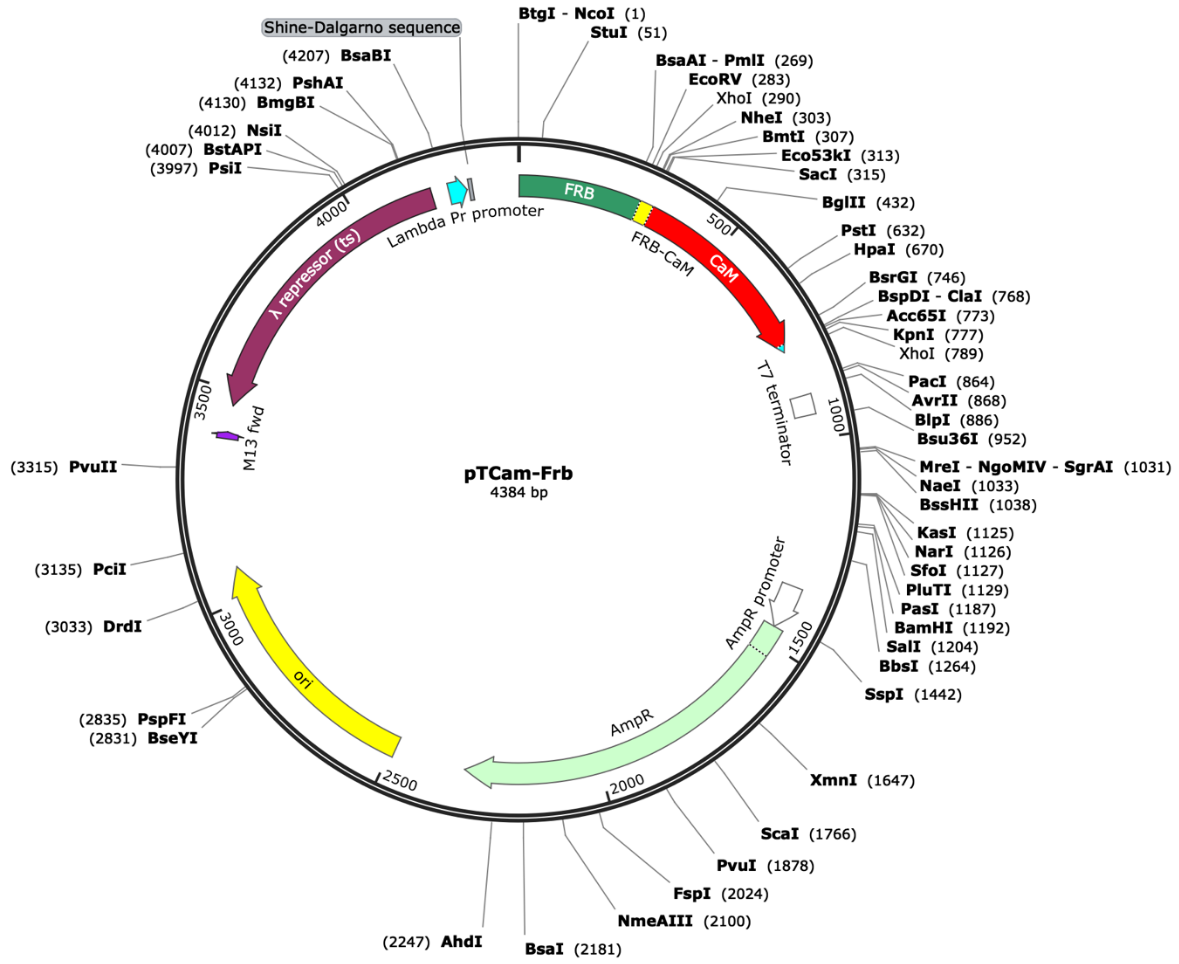


#### DNA sequence (*Nco*I-*Xho*I fragment)

ccATGGTAGCCATCCTCTGGCATGAGATGTGGCATGAAGGTCTAGAAGAGGCCTCTCGCTTGTACTTTGGGGAGA
GGAACGTCAAAGGCATGTTTGAGGTGCTGGAGCCCCTGCATGCTATGATGGAACGCGGTCCCCAGACCCTGAAGG
AAACGTCCTTTAATCAGGCATATGGTCGAGATTTAATGGAGGCACAAGAATGGTGCCGAAAGTACATGAAATCAG
GGAACGTCAAGGACCTCACCCAAGCCTGGGACCTCTACTATCACGTGTTCAGACGGATATCAGGCTCGAGTGGCC
CGGCTAGCCCGAGCTCCCATATGGCTGACCAACTGACAGAAGAGCAGATTGCAGAATTCAAAGAAGCTTTTTCAC
TATTTGACAAAGATGGTGATGGAACTATAACAACAAAGGAATTGGGAACTGTAATGAGATCTCTTGGGCAGAATC
CCACAGAAGCAGAGTTACAGGACATGATTAATGAAGTAGATGCTGATGGTAATGGCACAATTGACTTCCCTGAAT
TTCTGACAATGATGGCAAGAAAAATGAAAGACACAGACAGTGAAGAAGAAATTAGAGAAGCATTCCGTGTGTTTG
ATAAGGATGGCAATGGCTATATTAGTGCTGCAGAACTTCGCCATGTGATGACAAACCTTGGAGAGAAGTTAACAG
ATGAAGAAGTTGATGAAATGATCAGGGAAGCAGATATTGATGGTGATGGTCAAGTAAACTATGAAGAGTTTGTAC
AAATGATGACAGCGAAATCGATGGTACCCGCATGCTAActcgag

#### pK1Cam-V$_{9A}$ ( 3333 bp)

#### Map

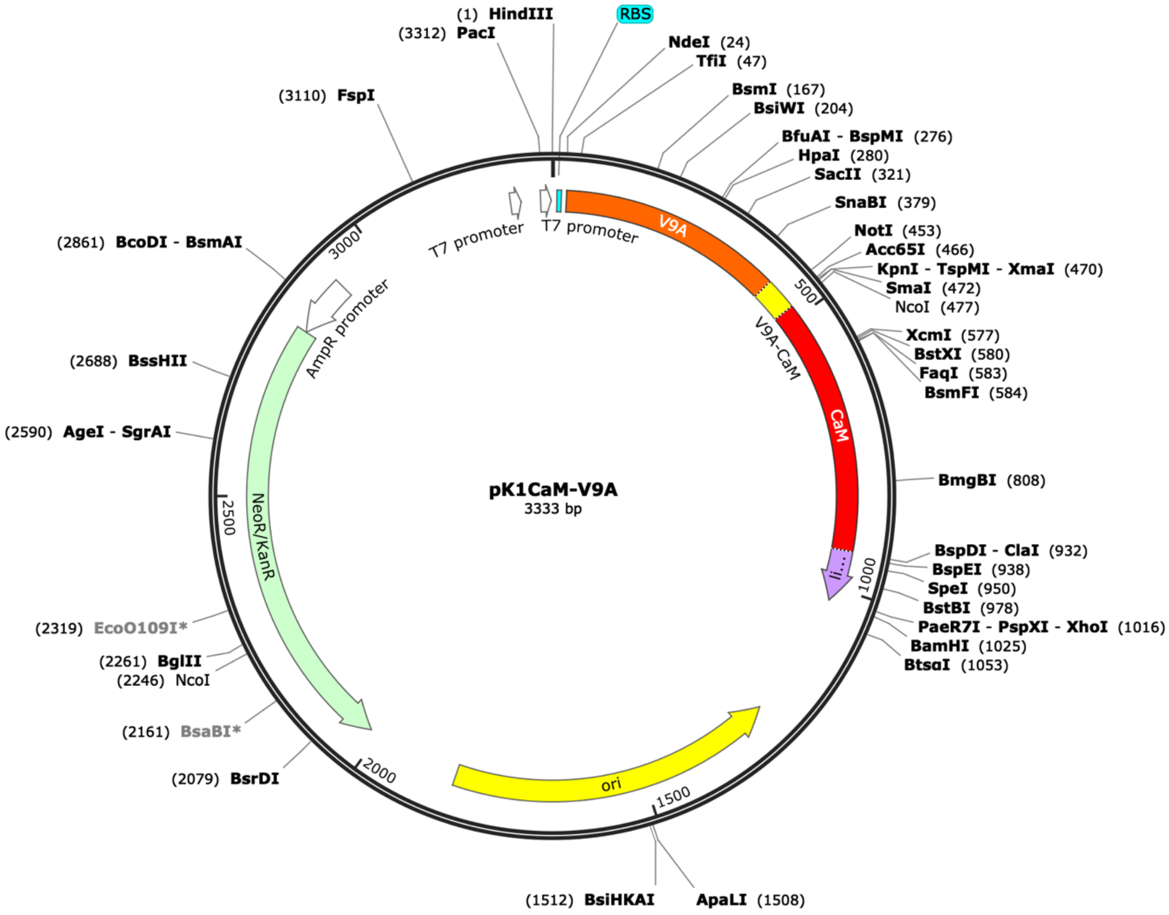


#### Full DNA sequence

**aagctt**ataagaaggagatatacatATGGCTGACGTTCAGCTGCAGGAATCTGGTGGTGGTTCTGTTCAGGCGGG
TGGTTCTCTGCGTCTGTCTTGCGCGGCTAGCGGTGACACCTTCTCTTCTTACTCTATGGCGTGGTTCCGTCAGGC
GCCGGGTAAAGAATGCGAACTGGTTTCTAACATCCTGCGTGACGGTACTACCACGTACGCCGGCTCTGTTAAAGG
TCGTTTCACCATCTCTCGTGACGACGCGAAAAACACCGTTTACCTGCAGATGGTTAACCTGAAATCTGAAGACAC
CGCGCGTTACTACTGCGCCGCGGACTCTGGTACTCAGCTGGGTTACGTTGGTGCGGTTGGTCTGTCTTGCCTGGA
CTACGTAATGGACTACTGGGGTAAAGGTACTCAGGTTACCGTTTCTTCTGAACCGAAAACCCCGAAACCGCAGCC
AGCGGCCGCTGAAAAGGTACCCGGGTCCATGGCTAGCGCCGACCAGCTGACCGAGGAGCAGATCGCCGAGTTCAA
GGAGGCCTTCTCCCTGTTCGACAAGGACGGCGACGGCACCATCACCACCAAGGAGCTGGGCACCGTCATGCGGTC
CCTGGGCCAGAACCCCACCGAGGCCGAGCTTCAGGACATGATCAACGAGGTCGACGCCGACGGCAACGGCACCAT
CGACTTCCCCGAGTTCCTGACCATGATGGCCCGGAAGATGAAGGACACCGACTCCGAGGAGGAGATCCGGGAGGC
CTTCCGGGTCTTCGACAAGGACGGCAACGGCTATATCTCCGCCGCCGAGCTGCGGCACGTCATGACCAACCTGGG
CGAGAAGCTGACCGACGAGGAGGTCGACGAGATGATCCGGGAGGCCGACATCGACGGCGACGGCCAGGTCAACTA
TGAGGAGTTCGTCCAGATGATGACCGCCAAATCGATGTCCGGAGGTGGCACTAGTGCTTCAGGTCTGAACGACAT
CTTCGAAGCTCAGAAAATCGAATGGCACGAAGGCGGCACCCTCGAGTAA**ggatcc**ctgggcctcatgggccttcc
tttcactgcccgctttccagtcgggaaacctgtcgtgccagctgcattaacatggtcatagctgtttccttgcgt
attgggcgctctccgcttcctcgctcactgactcgctgcgctcggtcgttcgggtaaagcctggggtgcctaatg
agcaaaaggccagcaaaaggccaggaaccgtaaaaaggccgcgttgctggcgtttttccataggctccgcccccc
tgacgagcatcacaaaaatcgacgctcaagtcagaggtggcgaaacccgacaggactataaagataccaggcgtt
tccccctggaagctccctcgtgcgctctcctgttccgaccctgccgcttaccggatacctgtccgcctttctccc
ttcgggaagcgtggcgctttctcatagctcacgctgtaggtatctcagttcggtgtaggtcgttcgctccaagct
gggctgtgtgcacgaaccccccgttcagcccgaccgctgcgccttatccggtaactatcgtcttgagtccaaccc
ggtaagacacgacttatcgccactggcagcagccactggtaacaggattagcagagcgaggtatgtaggcggtgc
tacagagttcttgaagtggtggcctaactacggctacactagaagaacagtatttggtatctgcgctctgctgaa

gccagttaccttcggaaaaagagttggtagctcttgatccggcaaacaaaccaccgctggtagcggtggttttt
tgtttgcaagcagcagattacgcgcagaaaaaaaggatctcaagaagatcctttgatcttttctacggggtctga
cgctcagtggaacgaaaactcacgttaagggattttggtcatgagattatcaaaaaggatcttcacctagatcct
tttaaattaaaaatgaagttttaaatcaatctaaagtatatatgagtaaacttggtctgacagttattagaaaaa
ttcatccagcagacgataaaacgcaatacgctggctatccggtgccgcaatgccatacagcaccagaaaacgatc
cgcccattcgccgcccagttcttccgcaatatcacgggtggccagcgcaatatcctgataacgatccgccacgcc
cagacggccgcaatcaataaagccgctaaaacggccattttccaccataatgttcggcaggcacgcatcaccatg
ggtcaccaccagatcttcgccatccggcatgctcgctttcagacgcgcaaacagctctgccggtgccaggccctg
atgttcttcatccagatcatcctgatccaccaggcccgcttccatacgggtacgcgcacgttcaatacgatgttt
cgcctgatgatcaaacggacaggtcgccgggtccagggtatgcagacgacgcatggcatccgccataatgctcac
tttttctgccggcgccagatggctagacagcagatcctgacccggcacttcgcccagcagcagccaatcacggcc
cgcttcggtcaccacatccagcaccgccgcacacggaacaccggtggtggccagccagctcagacgcgccgcttc
atcctgcagctcgttcagcgcaccgctcagatcggttttcacaaacagcaccggacgaccctgcgcgctcagacg
aaacaccgccgcatcagagcagccaatggtctgctgcgcccaatcatagccaaacagacgttccacccacgctgc
cgggctacccgcatgcaggccatcctgttcaatcatactcttcctttttcaatattattgaagcatttatcaggg
ttattgtctcatgagcggatacatatttgaatgtatttagaaaaataaacaaataggggttccgcgcacatttcc
ccgaaaagtgccacctaaattgtaagcgttaatattttgttaaaattcgcgttaaatttttgttaaatcagctca
ttttttaaccaataggccgaaatcggcaaaatcccttataaatcaaaagaatagaccgagatagggttgagtggc
cgctacagggcgctcccattcgccattcaggctgcgcaactgttgggaagggcgtttcggtgcgggcctcttcgc
tattacgccagctggcgaaagggggatgtgctgcaaggcgattaagttgggtaacgccagggttttcccagtcac
gacgttgtaaaacgacggccagtgagcgcgacgtaatacgactcactatagggcgaattgaaggaaggccgtcaa
ggccgcattaattaatacgactcactatagggg

#### pK2Cam-V$_{9A}$( 3318 bp)

#### Map

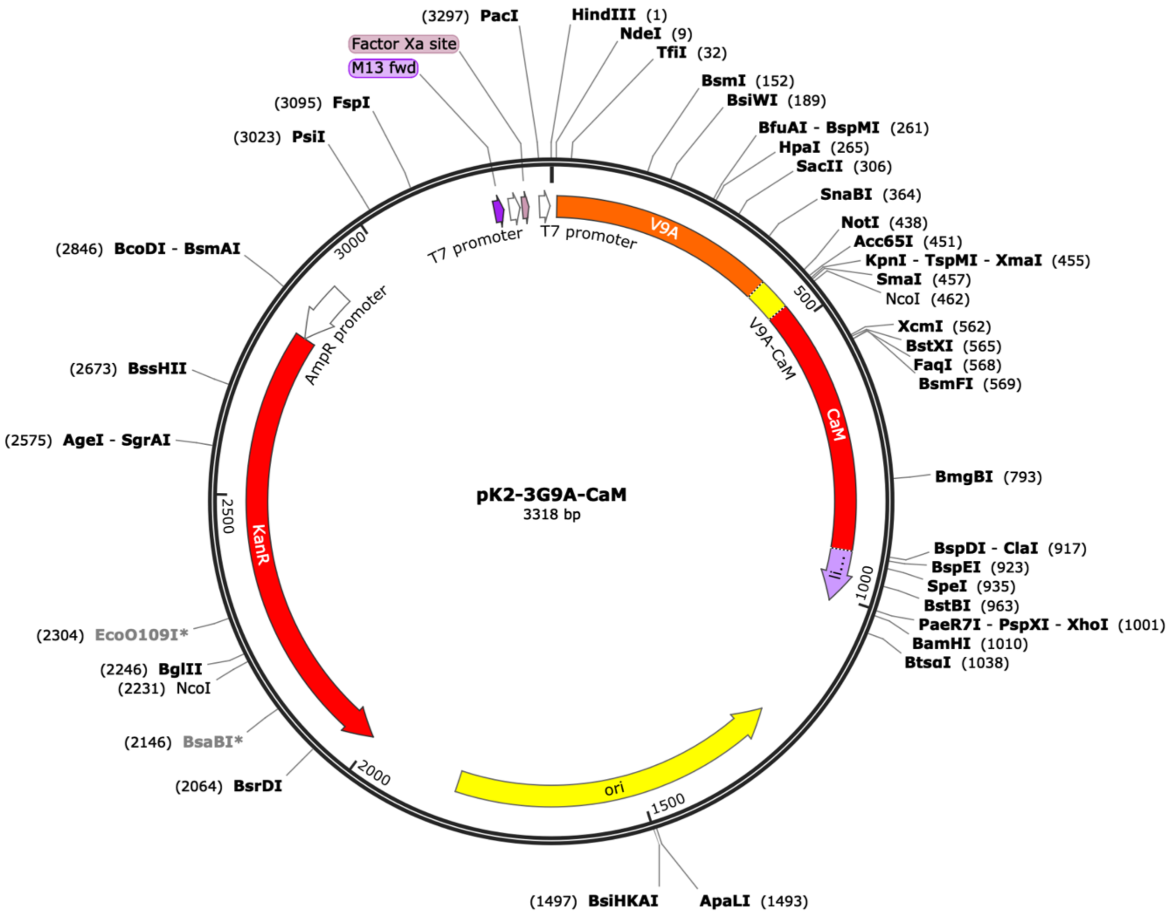


#### DNA sequence (*Hind*III-B*amH*I fragment)

**aagctt**gcatATGGCTGACGTTCAGCTGCAGGAATCTGGTGGTGGTTCTGTTCAGGCGGGTGGTTCTCTGCGTCT
GTCTTGCGCGGCTAGCGGTGACACCTTCTCTTCTTACTCTATGGCGTGGTTCCGTCAGGCGCCGGGTAAAGAATG
CGAACTGGTTTCTAACATCCTGCGTGACGGTACTACCACGTACGCCGGCTCTGTTAAAGGTCGTTTCACCATCTC
TCGTGACGACGCGAAAAACACCGTTTACCTGCAGATGGTTAACCTGAAATCTGAAGACACCGCGCGTTACTACTG
CGCCGCGGACTCTGGTACTCAGCTGGGTTACGTTGGTGCGGTTGGTCTGTCTTGCCTGGACTACGTAATGGACTA
CTGGGGTAAAGGTACTCAGGTTACCGTTTCTTCTGAACCGAAAACCCCGAAACCGCAGCCAGCGGCCGCTGAAAA
GGTACCCGGGTCCATGGCTAGCGCCGACCAGCTGACCGAGGAGCAGATCGCCGAGTTCAAGGAGGCCTTCTCCCT
GTTCGACAAGGACGGCGACGGCACCATCACCACCAAGGAGCTGGGCACCGTCATGCGGTCCCTGGGCCAGAACCC
CACCGAGGCCGAGCTTCAGGACATGATCAACGAGGTCGACGCCGACGGCAACGGCACCATCGACTTCCCCGAGTT
CCTGACCATGATGGCCCGGAAGATGAAGGACACCGACTCCGAGGAGGAGATCCGGGAGGCCTTCCGGGTCTTCGA
CAAGGACGGCAACGGCTATATCTCCGCCGCCGAGCTGCGGCACGTCATGACCAACCTGGGCGAGAAGCTGACCGA
CGAGGAGGTCGACGAGATGATCCGGGAGGCCGACATCGACGGCGACGGCCAGGTCAACTATGAGGAGTTCGTCCA
GATGATGACCGCCAAATCGATGTCCGGAGGTGGCACTAGTGCTTCAGGTCTGAACGACATCTTCGAAGCTCAGAA
AATCGAATGGCACGAAGGCGGCACCCTCGAGTAA**ggatcc**

#### pK1Cam-V$_{9A}$-Zip (3522 bp)

#### Map

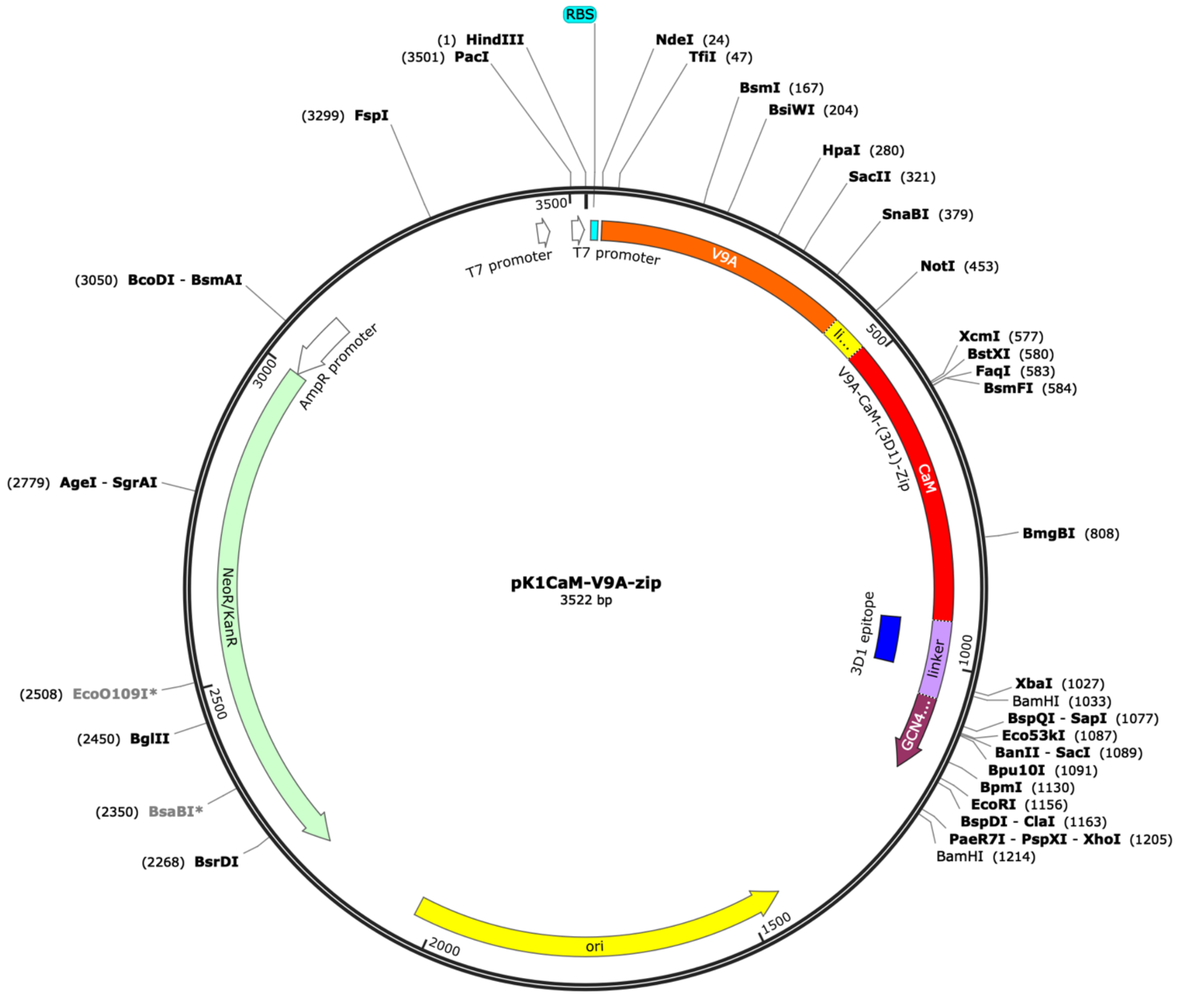


#### DNA sequence (*Hind*III-B*amH*I fragment)

**aagctt**ataagaaggagatatacatATGGCTGACGTTCAGCTGCAGGAATCTGGTGGTGGTTCTGTTCAGGCGGG
TGGTTCTCTGCGTCTGTCTTGCGCGGCTAGCGGTGACACCTTCTCTTCTTACTCTATGGCGTGGTTCCGTCAGGC
GCCGGGTAAAGAATGCGAACTGGTTTCTAACATCCTGCGTGACGGTACTACCACGTACGCCGGCTCTGTTAAAGG
TCGTTTCACCATCTCTCGTGACGACGCGAAAAACACCGTTTACCTGCAGATGGTTAACCTGAAATCTGAAGACAC
CGCGCGTTACTACTGCGCCGCGGACTCTGGTACTCAGCTGGGTTACGTTGGTGCGGTTGGTCTGTCTTGCCTGGA
CTACGTAATGGACTACTGGGGTAAAGGTACTCAGGTTACCGTTTCTTCTGAACCGAAAACCCCGAAACCGCAGCC
AGCGGCCGCTGAAAAGGTACCCGGGTCCATGGCTAGCGCCGACCAGCTGACCGAGGAGCAGATCGCCGAGTTCAA
GGAGGCCTTCTCCCTGTTCGACAAGGACGGCGACGGCACCATCACCACCAAGGAGCTGGGCACCGTCATGCGGTC
CCTGGGCCAGAACCCCACCGAGGCCGAGCTTCAGGACATGATCAACGAGGTCGACGCCGACGGCAACGGCACCAT
CGACTTCCCCGAGTTCCTGACCATGATGGCCCGGAAGATGAAGGACACCGACTCCGAGGAGGAGATCCGGGAGGC
CTTCCGGGTCTTCGACAAGGACGGCAACGGCTATATCTCCGCCGCCGAGCTGCGGCACGTCATGACCAACCTGGG
CGAGAAGCTGACCGACGAGGAGGTCGACGAGATGATCCGGGAGGCCGACATCGACGGCGACGGCCAGGTCAACTA
TGAGGAGTTCGTCCAGATGATGACCGCCAAATCGAAGTTCTCGCCGGATGTACTGGAAACGGTGCCGGCGTCACC
CGGATTGCGGCGGCCGTCGCTGGGCGCAGTGGAACGCCACTGCAGGTCGACTCTAGAGGATCCCCGGGTACCTAT
CCAGCGTATGAAACAGCTGGAAGACAAAGTTGAAGAGCTCCTGAGCAAAAACTACCACCTGGAGAACGAAGTTGC
GCGCCTGAAAAAACTGGTGGGTGAACGTGGGAATTCATCGATATAActaagtaatatggtgcactctcagtacaa
tctgctcgagtaa**ggatcc**

#### pK1Cam-V$_{9A}$-TM-Zip ( 3612 bp)

#### Map

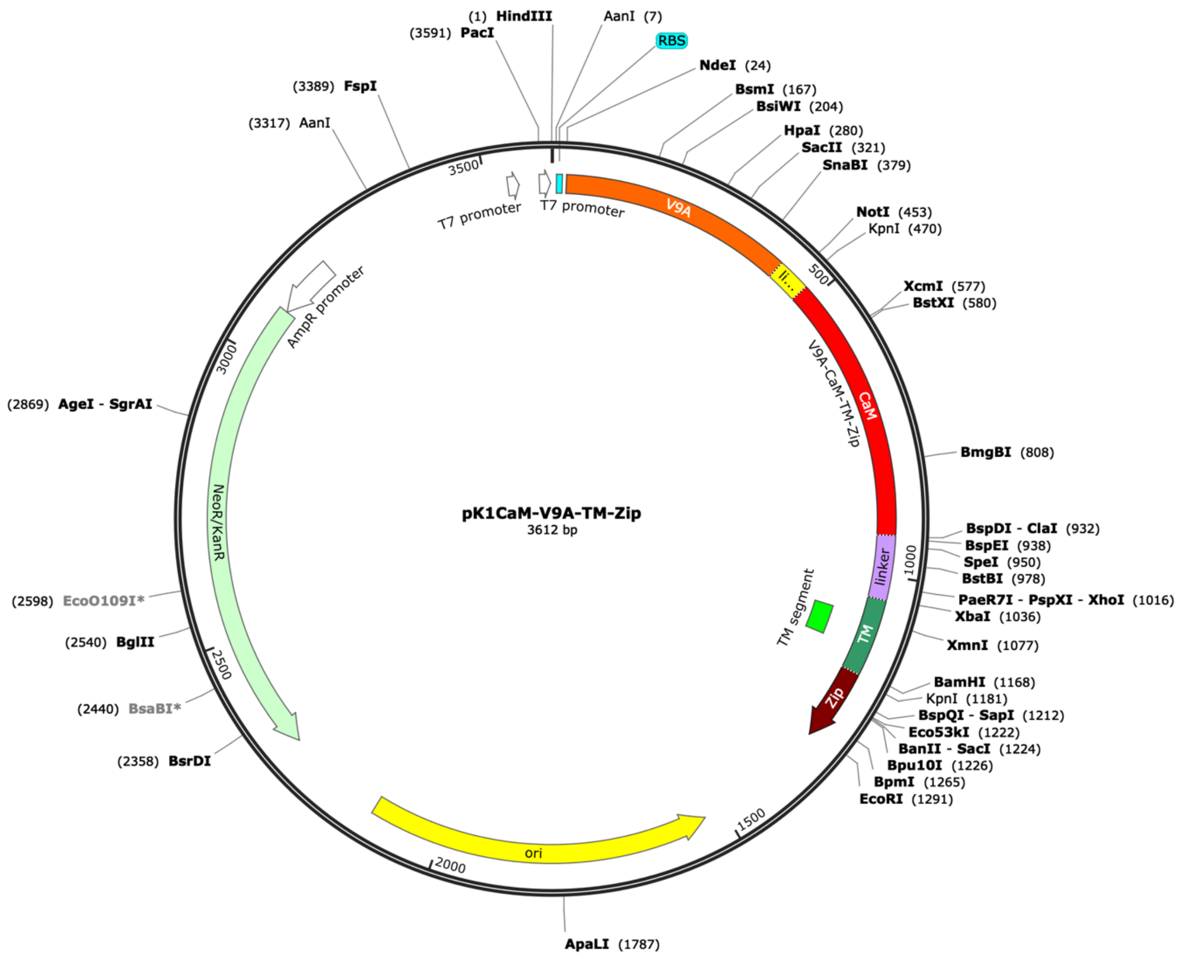


#### Full DNA sequence

**<u>aagctt</u>**ataagaaggagatatacatATGGCTGACGTTCAGCTGCAGGAATCTGGTGGTGGTTCTGTTCAGGCGGG
TGGTTCTCTGCGTCTGTCTTGCGCGGCTAGCGGTGACACCTTCTCTTCTTACTCTATGGCGTGGTTCCGTCAGGC
GCCGGGTAAAGAATGCGAACTGGTTTCTAACATCCTGCGTGACGGTACTACCACGTACGCCGGCTCTGTTAAAGG
TCGTTTCACCATCTCTCGTGACGACGCGAAAAACACCGTTTACCTGCAGATGGTTAACCTGAAATCTGAAGACAC
CGCGCGTTACTACTGCGCCGCGGACTCTGGTACTCAGCTGGGTTACGTTGGTGCGGTTGGTCTGTCTTGCCTGGA
CTACGTAATGGACTACTGGGGTAAAGGTACTCAGGTTACCGTTTCTTCTGAACCGAAAACCCCGAAACCGCAGCC
AGCGGCCGCTGAAAAGGTACCCGGGTCCATGGCTAGCGCCGACCAGCTGACCGAGGAGCAGATCGCCGAGTTCAA
GGAGGCCTTCTCCCTGTTCGACAAGGACGGCGACGGCACCATCACCACCAAGGAGCTGGGCACCGTCATGCGGTC
CCTGGGCCAGAACCCCACCGAGGCCGAGCTTCAGGACATGATCAACGAGGTCGACGCCGACGGCAACGGCACCAT
CGACTTCCCCGAGTTCCTGACCATGATGGCCCGGAAGATGAAGGACACCGACTCCGAGGAGGAGATCCGGGAGGC
CTTCCGGGTCTTCGACAAGGACGGCAACGGCTATATCTCCGCCGCCGAGCTGCGGCACGTCATGACCAACCTGGG
CGAGAAGCTGACCGACGAGGAGGTCGACGAGATGATCCGGGAGGCCGACATCGACGGCGACGGCCAGGTCAACTA
TGAGGAGTTCGTCCAGATGATGACCGCCAAATCGATGTCCGGAGGTGGCACTAGTGCTTCAGGTCTGAACGACAT
CTTCGAAGCTCAGAAAATCGAATGGCACGAAGGCGGCACCCTCGAGCACTGCAGGTCGACTCTAGAGGATCTAAA
ATTTATTCTACGTCGCTGTCTGGAAGCGATTCCGACGCTATTTATTCTTATTACTATTTCGTTCTTTATGATGCG
CCTCGCGCCGGGAAGCCCTTTTACCGGCGAACGTACTTTAGCGGATCCCCGGGTACCTATCCAGCGTATGAAACA
GCTGGAAGACAAAGTTGAAGAGCTCCTGAGCAAAAACTACCACCTGGAGAACGAAGTTGCGCGCCTGAAAAAACT
GGTGGGTGAACGTGGGAATTCATCGTAAagatccctgggcctcatgggccttcctttcactgcccgctttccagt
cgggaaacctgtcgtgccagctgcattaacatggtcatagctgtttccttgcgtattgggcgctctccgcttcct
cgctcactgactcgctgcgctcggtcgttcgggtaaagcctggggtgcctaatgagcaaaaggccagcaaaaggc
caggaaccgtaaaaaggccgcgttgctggcgtttttccataggctccgcccccctgacgagcatcacaaaaatcg
acgctcaagtcagaggtggcgaaacccgacaggactataaagataccaggcgtttccccctggaagctccctcgt

gcgctctcctgttccgaccctgccgcttaccggatacctgtccgcctttctcccttcgggaagcgtggcgctttc
tcatagctcacgctgtaggtatctcagttcggtgtaggtcgttcgctccaagctgggctgtgtgcacgaaccccc
cgttcagcccgaccgctgcgccttatccggtaactatcgtcttgagtccaacccggtaagacacgacttatcgcc
actggcagcagccactggtaacaggattagcagagcgaggtatgtaggcggtgctacagagttcttgaagtggtg
gcctaactacggctacactagaagaacagtatttggtatctgcgctctgctgaagccagttaccttcggaaaaag
agttggtagctcttgatccggcaaacaaaccaccgctggtagcggtggtttttttgtttgcaagcagcagattac
gcgcagaaaaaaaggatctcaagaagatcctttgatcttttctacggggtctgacgctcagtggaacgaaaactc
acgttaagggattttggtcatgagattatcaaaaaggatcttcacctagatccttttaaattaaaaatgaagttt
taaatcaatctaaagtatatatgagtaaacttggtctgacagttattagaaaaattcatccagcagacgataaaa
cgcaatacgctggctatccggtgccgcaatgccatacagcaccagaaaacgatccgcccattcgccgcccagttc
ttccgcaatatcacgggtggccagcgcaatatcctgataacgatccgccacgcccagacggccgcaatcaataaa
gccgctaaaacggccatttccaccataatgttcggcaggcacgcatcaccatgggtcaccaccagatcttcgcc
atccggcatgctcgctttcagacgcgcaaacagctctgccggtgccaggccctgatgttcttcatccagatcatc
ctgatccaccaggcccgcttccatacgggtacgcgcacgttcaatacgatgtttcgcctgatgatcaaacggaca
ggtcgccgggtccagggtatgcagacgacgcatggcatccgccataatgctcactttttctgccggcgccagatg
gctagacagcagatcctgacccggcacttcgcccagcagcagccaatcacggcccgcttcggtcaccacatccag
caccgccgcacacggaacaccggtggtggccagccagctcagacgcgccgcttcatcctgcagctcgttcagcgc
accgctcagatcggttttcacaaacagcaccggacgaccctgcgcgctcagacgaaacaccgccgcatcagagca
gccaatggtctgctgcgcccaatcatagccaaacagacgttccacccacgctgccgggctacccgcatgcaggcc
atcctgttcaatcatactcttcctttttcaatattattgaagcatttatcagggttattgtctcatgagcggata
catatttgaatgtatttagaaaaataaacaaataggggttccgcgcacatttccccgaaaagtgccacctaaatt
gtaagcgttaatattttgttaaaattcgcgttaaatttttgttaaatcagctcattttttaaccaataggccgaa
atcggcaaaatcccttataaatcaaaagaatagaccgagatagggttgagtggccgctacagggcgctcccattc
gccattcaggctgcgcaactgttgggaagggcgtttcggtgcgggcctcttcgctattacgccagctggcgaaag
ggggatgtgctgcaaggcgattaagttgggtaacgccagggttttcccagtcacgacgttgtaaaacgacggcca
gtgagcgcgacgtaatacgactcactatagggcgaattgaaggaaggccgtcaaggccgcattaattaatacgac
tcactatagggg

### pK1Cam-Frb

#### Map

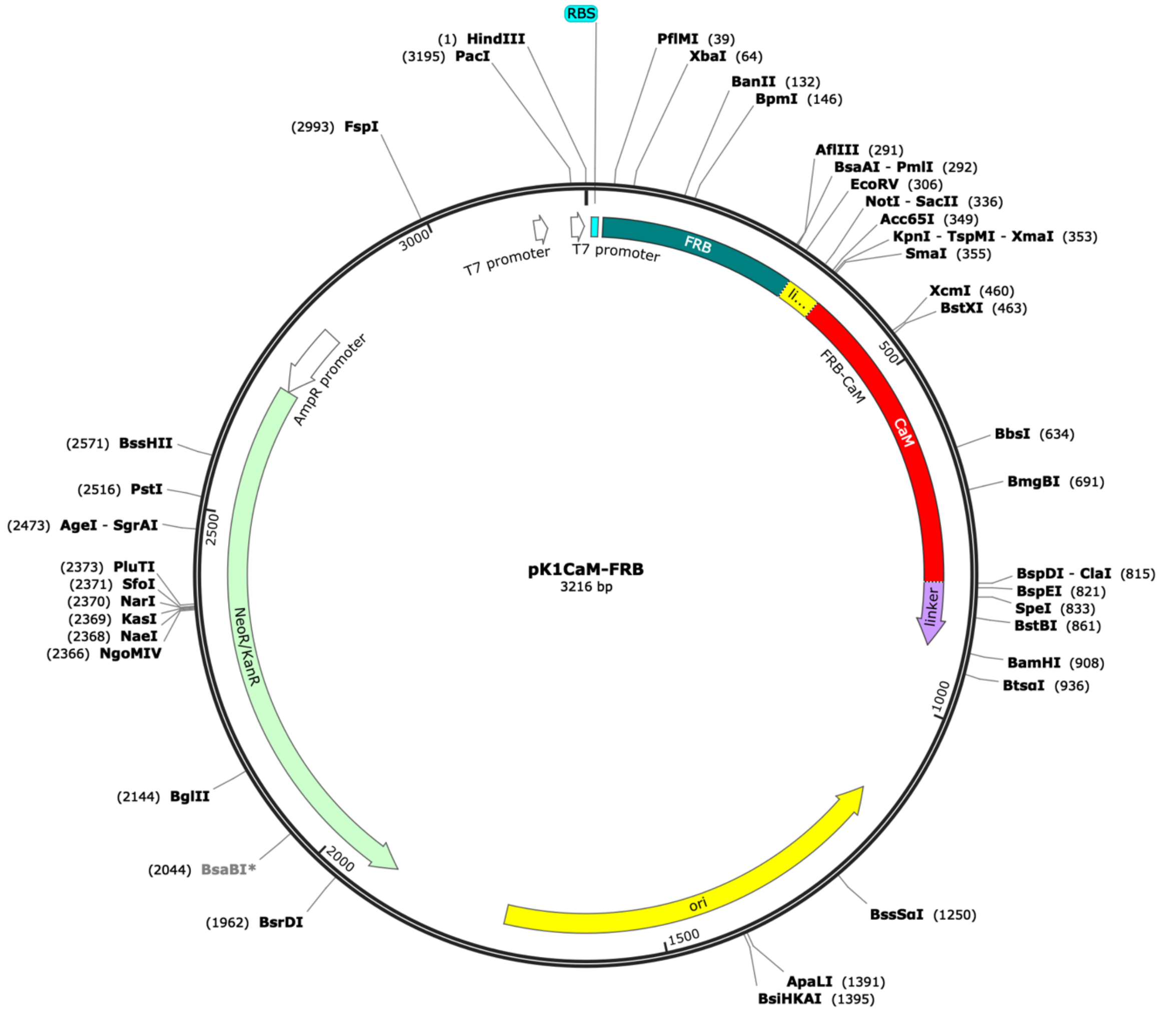


#### DNA sequence (*Hind*III-B*amH*I fragment)

**<u>aagctt</u>**ataagaaggagatatacatATGGTAGCCATCCTCTGGCATGAGATGTGGCATGAAGGTCTAGAAGAGGC
CTCTCGCTTGTACTTTGGGGAGAGGAACGTCAAAGGCATGTTTGAGGTGCTGGAGCCCCTGCATGCTATGATGGA
ACGCGGTCCCCAGACCCTGAAGGAAACGTCCTTTAATCAGGCATATGGTCGAGATTTAATGGAGGCACAAGAATG
GTGCCGAAAGTACATGAAATCAGGGAACGTCAAGGACCTCACCCAAGCCTGGGACCTCTACTATCACGTGTTCAG
ACGGATATCAGGCTCGAGTGGCCCGGCTAGCCCCGCGGCCGCTGAAAAGGTACCCGGGTCCATGGCTAGCGCCGA
CCAGCTGACCGAGGAGCAGATCGCCGAGTTCAAGGAGGCCTTCTCCCTGTTCGACAAGGACGGCGACGGCACCAT
CACCACCAAGGAGCTGGGCACCGTCATGCGGTCCCTGGGCCAGAACCCCACCGAGGCCGAGCTTCAGGACATGAT
CAACGAGGTCGACGCCGACGGCAACGGCACCATCGACTTCCCCGAGTTCCTGACCATGATGGCCCGGAAGATGAA
GGACACCGACTCCGAGGAGGAGATCCGGGAGGCCTTCCGGGTCTTCGACAAGGACGGCAACGGCTATATCTCCGC
CGCCGAGCTGCGGCACGTCATGACCAACCTGGGCGAGAAGCTGACCGACGAGGAGGTCGACGAGATGATCCGGGA
GGCCGACATCGACGGCGACGGCCAGGTCAACTATGAGGAGTTCGTCCAGATGATGACCGCCAAATCGATGTCCGG
AGGTGGCACTAGTGCTTCAGGTCTGAACGACATCTTCGAAGCTCAGAAAATCGAATGGCACGAAGGCGGCACCCT
CGAGTAA**<u>ggatcc</u>**

#### pK1Cam-Frb-Zs (4465 bp)

#### Map

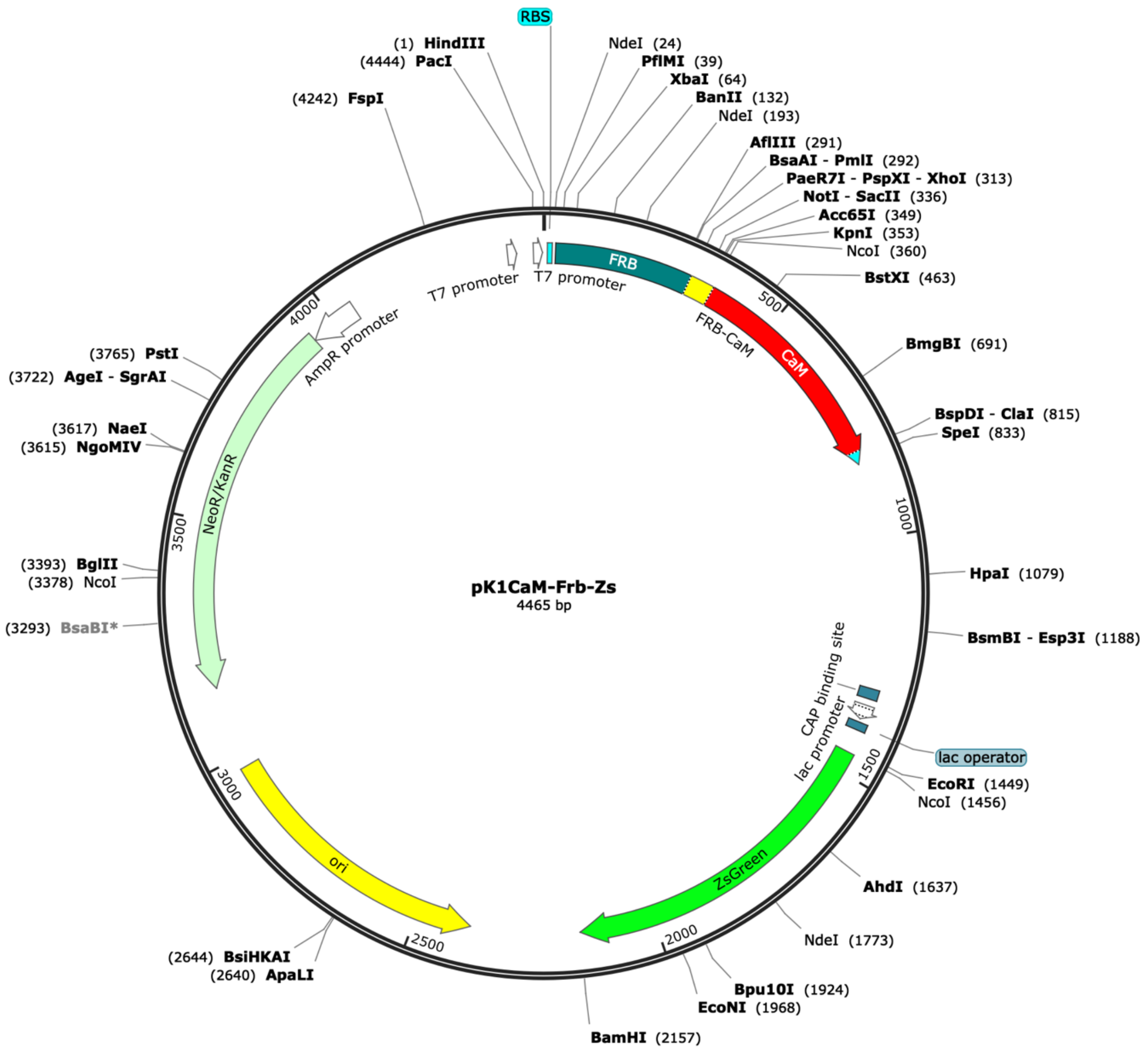


#### DNA sequence (*Hind*III-B*amH*I fragment)

**aagctt**atataagaaggagatatacatATGGTAGCCATCCTCTGGCATGAGATGTGGCATGAAGGTCTAGAAGAGGC
CTCTCGCTTGTACTTTGGGGAGAGGAACGTCAAAGGCATGTTTGAGGTGCTGGAGCCCCTGCATGCTATGATGGA
ACGCGGTCCCCAGACCCTGAAGGAAACGTCCTTTAATCAGGCATATGGTCGAGATTTAATGGAGGCACAAGAATG
GTGCCGAAAGTACATGAAATCAGGGAACGTCAAGGACCTCACCCAAGCCTGGGACCTCTACTATCACGTGTTCAG
ACGGATATCAGGCTCGAGTGGCCCGGCTAGCCCCGCGGCCGCTGAAAAGGTACCCGGGTCCATGGCTAGCGCCGA
CCAGCTGACCGAGGAGCAGATCGCCGAGTTCAAGGAGGCCTTCTCCCTGTTCGACAAGGACGGCGACGGCACCAT
CACCACCAAGGAGCTGGGCACCGTCATGCGGTCCCTGGGCCAGAACCCCACCGAGGCCGAGCTTCAGGACATGAT
CAACGAGGTCGACGCCGACGGCAACGGCACCATCGACTTCCCCGAGTTCCTGACCATGATGGCCCGGAAGATGAA
GGACACCGACTCCGAGGAGGAGATCCGGGAGGCCTTCCGGGTCTTCGACAAGGACGGCAACGGCTATATCTCCGC
CGCCGAGCTGCGGCACGTCATGACCAACCTGGGCGAGAAGCTGACCGACGAGGAGGTCGACGAGATGATCCGGGA
GGCCGACATCGACGGCGACGGCCAGGTCAACTATGAGGAGTTCGTCCAGATGATGACCGCCAAATCGATGTCCGG
AGGTGGCACTAGTTAAtaaatcaaattcagccgatagcggaacgggaaggcgactggagtgccatgtccggtttt
caacaaaccatgcaaatgctgaatgagggcatcgttcccactgcgatgctggttgccaacgatcagatggcgctg
ggcgcaatgcgcgccattaccgagtccgggctgcgcgttggtgcggatatctcggtagtgggatacgacgatacc
gaagacagctcatgttatatcccgccgttaaccaccatcaaacaggattttcgcctgctggggcaaaccagcgtg
gaccgcttgctgcaactctctcagggccaggcggtgaagggcaatcagctgttgcccgtctcactggtgaaaaga
aaaaccaccctggcgcccaatacgcaaaccgcctctccccgcgcgttggccgattcattaatgcagctggcacga

caggtttcccgactggaaagcgggcagtgagcgcaacgcaattaatgtgagttagctcactcattaggcacccca
ggctttacactttatgcttccggctcgtatgttgtgtggaattgtgagcggataacaatttcacacaggaaacag
ctatgaccatgattacggattcagaattctccATGGGCCAGTCAAAGCACGGTCTAACAAAAGAAATGACAATGA
AATACCGTATGGAAGGGTGCGTCGATGGACATAAATTTGTGATCACGGGAGAGGGCATTGGATATCCGTTCAAAG
GGAAACAGGCTATTAATCTGTGTGTGGTCGAAGGTGGACCATTGCCATTTGCCGAAGACATATTGTCAGCTGCCT
TTATGTACGGAAACAGGGTTTTCACTGAATATCCTCAAGACATAGCTGACTATTTCAAGAACTCGTGTCCTGCTG
GTTATACATGGGACAGGTCTTTTCTCTTTGAGGATGGAGCAGTTTGCATATGTAATGCAGATATAACAGTGAGTG
TTGAAGAAAACTGCATGTATCATGAGTCCAAATTTTATGGAGTGAATTTTCCTGCTGATGGACCTGTGATGAAAA
AGATGACAGATAACTGGGAGCCATCCTGCGAGAAGATCATACCAGTACCTAAGCAGGGGATATTGAAAGGGGATG
TCTCCATGTACCTCCTTCTGAAGGATGGTGGGCGTTTACGGTGCCAATTCGACACAGTTTACAAAGCAAAGTCTG
TGCCAAGAAAGATGCCGGACTGGCACTTCATCCAGCATAAGCTCACCCGTGAAGACCGCAGCGATGCTAAGAATC
AGAAATGGCATCTGACAGAACATGCTATTGCATCCGGATCTGCATTGCCCGGGTAA**ggatcc**

#### pK1Cam-$V_{1K}$

#### Map

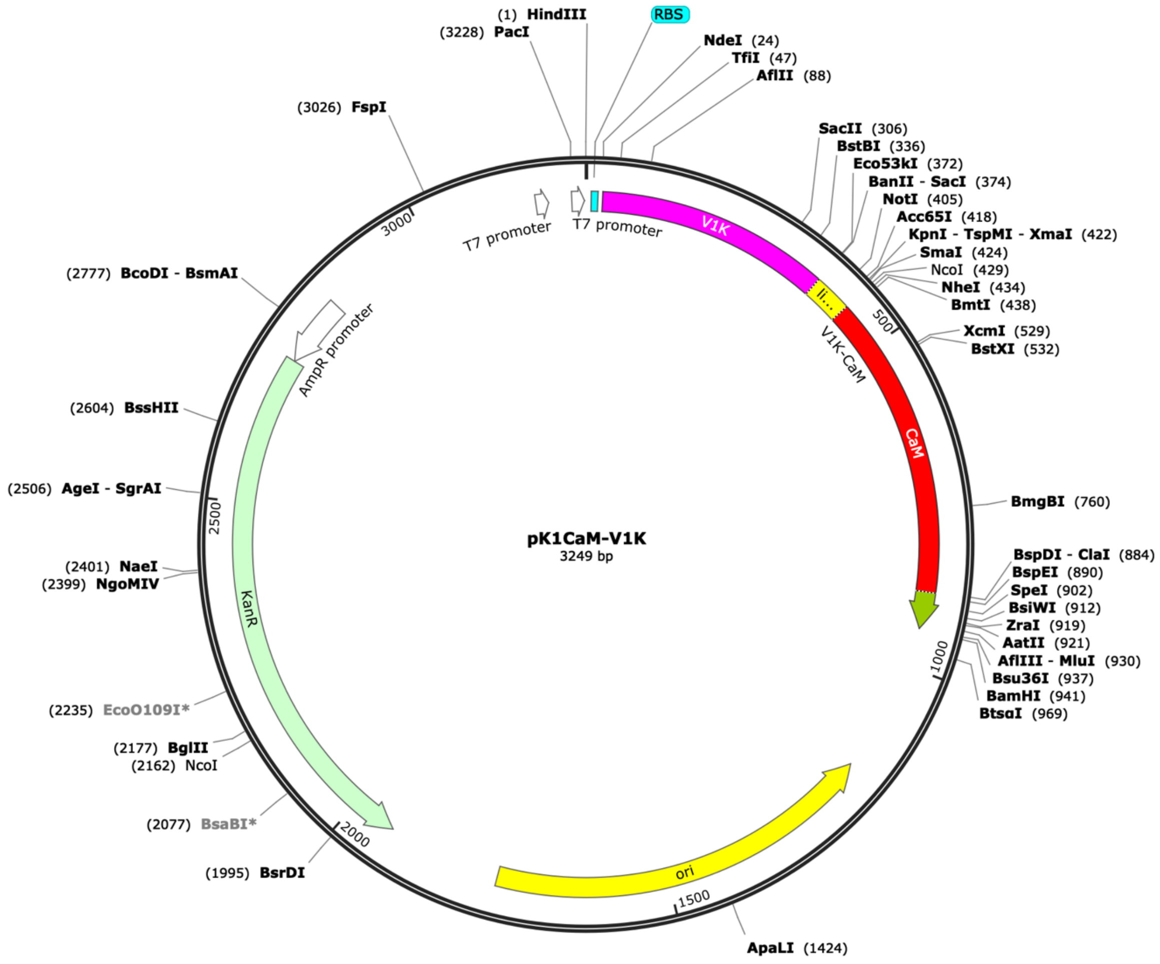


#### DNA sequence (*Hind*III-B*amH*I fragment)

**aagctt**ataagaaggagatatacatATGGCGCAGGTTCAGCTTGTTGAATCTGGTGGTGCGCTGGTTCAGCCGGG
TGGTTCTCTGCGCTTAAGCTGCGCGGCTTCCGGTTTCCCGGTTAACCGTTACTCTATGCGTTGGTATCGTCAGGC
GCCGGGTAAAGAACGTGAATGGGTTGCGGGTATGTCTTCTGCGGGTGACCGTTCTTCTTACGAAGACTCTGTTAA
AGGTCGTTTCACCATCTCTCGTGACGACGCGCGTAACACCGTTTACCTGCAGATGAACTCTCTGAAACCGGAAGA
CACCGCGGTTTACTACTGCAACGTTAACGTTGGTTTCGAATACTGGGGTCAGGGCACCCAGGTTACCGTGAGCTC
TGAACCGAAAACCCCGAAACCGCAGCCAGCGGCCGCTGAAAAGGTACCCGGGTCCATGGCTAGCGCCGACCAGCT
GACCGAGGAGCAGATCGCCGAGTTCAAGGAGGCCTTCTCCCTGTTCGACAAGGACGGCGACGGCACCATCACCAC
CAAGGAGCTGGGCACCGTCATGCGGTCCCTGGGCCAGAACCCCACCGAGGCCGAGCTTCAGGACATGATCAACGA
GGTCGACGCCGACGGCAACGGCACCATCGACTTCCCCGAGTTCCTGACCATGATGGCCCGGAAGATGAAGGACAC
CGACTCCGAGGAGGAGATCCGGGAGGCCTTCCGGGTCTTCGACAAGGACGGCAACGGCTATATCTCCGCCGCCGA
GCTGCGGCACGTCATGACCAACCTGGGCGAGAAGCTGACCGACGAGGAGGTCGACGAGATGATCCGGGAGGCCGA
CATCGACGGCGACGGCCAGGTCAACTATGAGGAGTTCGTCCAGATGATGACCGCCAAATCGATGTCCGGAGGTGG
CACTAGTTACCCGTACGACGTCCCAGATTACGCGTCCTAA**ggatcc**

#### pK1Cam-Barstar

#### Map

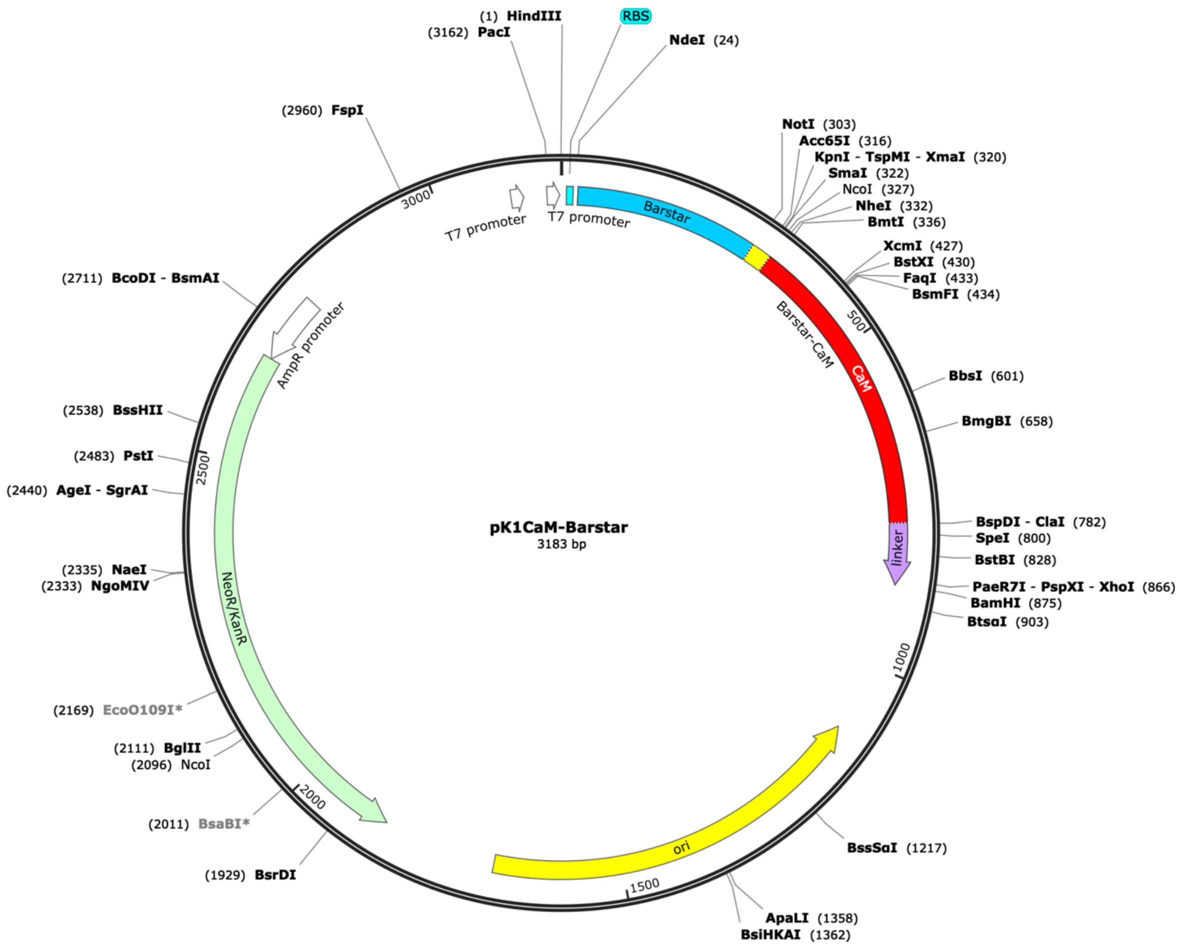


#### DNA sequence (*Hind*III-B*amH*I fragment)

**aagctt**atGaagaaggagatatacatATGGGTAAAAAAGCAGTCATTAACGGGGAACAAATCAGAAGTATCAGCGA
CCTCCACCAGACATTGAAAAAGGAGCTTGCCCTTCCGGAATACTACGGTGAAAACCTGGACGCTTTATGGGATTG
TCTGACCGGATGGGTGGAGTACCCGCTCGTTTTGGAATGGAGGCAGTTTGAACAAAGCAAGCAGCTGACTGAAAA
TGGCGCCGAGAGTGTGCTTCAGGTTTTCCGTGAAGCGAAAGCGGAAGGCTGCGACATCACCATCATACTTTCTGG
TGCGGCCGCTGAAAAGGTACCCGGGTCCATGGCTAGCGCCGACCAGCTGACCGAGGAGCAGATCGCCGAGTTCAA
GGAGGCCTTCTCCCTGTTCGACAAGGACGGCGACGGCACCATCACCACCAAGGAGCTGGGCACCGTCATGCGGTC
CCTGGGCCAGAACCCCACCGAGGCCGAGCTTCAGGACATGATCAACGAGGTCGACGCCGACGGCAACGGCACCAT
CGACTTCCCCGAGTTCCTGACCATGATGGCCCGGAAGATGAAGGACACCGACTCCGAGGAGGAGATCCGGGAGGC
CTTCCGGGTCTTCGACAAGGACGGCAACGGCTATATCTCCGCCGCCGAGCTGCGGCACGTCATGACCAACCTGGG
CGAGAAGCTGACCGACGAGGAGGTCGACGAGATGATCCGGGAGGCCGACATCGACGGCGACGGCCAGGTCAACTA
TGAGGAGTTCGTCCAGATGATGACCGCCAAATCGATGTCCGGAGGTGGCACTAGTGCTTCAGGTCTGAACGACAT
CTTCGAAGCTCAGAAAATCGAATGGCACGAAGGCGGCACCCTCGAGTAA**ggatcc**

#### pCam$_{VU8}$ (3006 bp)

#### Map

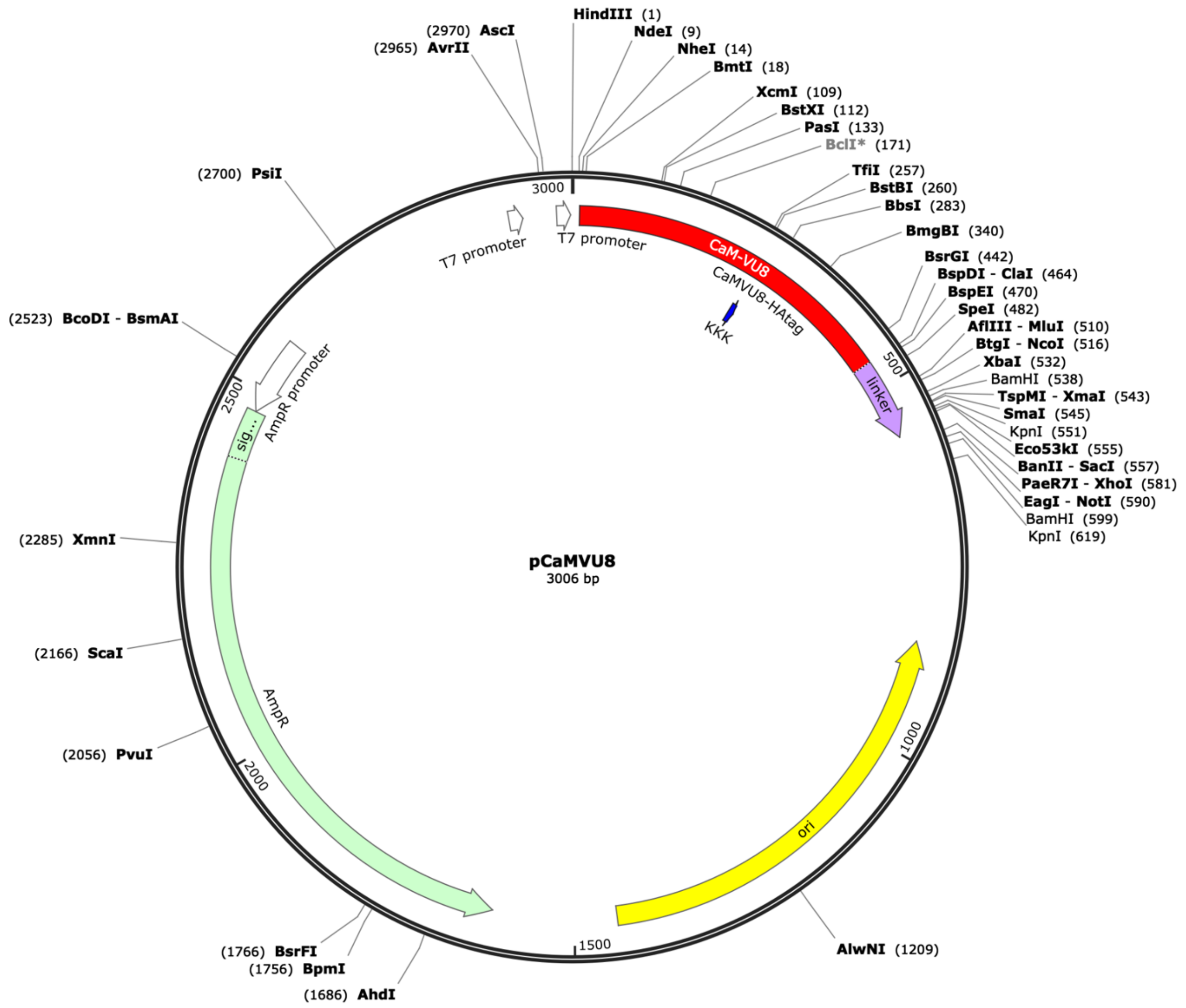


#### Full DNA sequence

**aagctt**gcatATGGCTAGCGCCGACCAGCTGACCGAGGAGCAGATCGCCGAGTTCAAGGAGGCGTTCTCCCTGTT
CGACAAGGACGGCGACGGCACCATCACCACCAAGGAGCTGGGCACCGTCATGCGGTCCCTGGGCCAGAACCCCAC
CGAGGCCGAGCTTCAGGACATGATCAACGAGGTCGACGCCGACGGCAACGGCACCATCGACTTCCCCGAATTCCT
GACCATGATGGCGCGCAAGATGAAGGACACCGATTCGAAGAAGAAGATCCGGGAGGCCTTCCGGGTCTTCGACAA
GGACGGCAACGGCTATATCTCCGCCGCCGAGCTGCGGCACGTCATGACCAACCTGGGCGAGAAGCTGACCGACGA
GGAGGTCGACGAGATGATCCGGGAGGCCGACATCGACGGCGACGGCCAGGTCAACTATGAGGAGTTTGTACAGAT
GATGACCGCCAAATCGATGTCCGGAGGTGGCACTAGTTACCCATACGATGTCCCAGATTACGCGTCCATGGCGCA
CTCGACTCTAGAGGATCCCCGGGTACCGAGCTCGAATTCATCGTAActaagtaatctcgagtagcggccgctt**gg**
**atcc**cagaattctaggtacctcttaattaactggcctcatgggccttccgctcactgcccgctttccagtcggga
aacctgtcgtgccagctgcattaacatggtcatagctgtttccttgcgtattgggcgctctccgcttcctcgctc
actgactcgctgcgctcggtcgttcgggtaaagcctggggtgcctaatgagcaaaaggccagcaaaaggccagga
accgtaaaaaggccgcgttgctggcgtttttccataggctccgcccccctgacgagcatcacaaaaatcgacgct
caagtcagaggtggcgaaacccgacaggactataaagataccaggcgtttccccctggaagctccctcgtgcgct
ctcctgttccgaccctgccgcttaccggatacctgtccgcctttctcccttcgggaagcgtggcgctttctcata
gctcacgctgtaggtatctcagttcggtgtaggtcgttcgctccaagctgggctgtgtgcacgaaccccccgttc
agcccgaccgctgcgccttatccggtaactatcgtcttgagtccaacccggtaagacacgacttatcgccactgg
cagcagccactggtaacaggattagcagagcgaggtatgtaggcggtgctacagagttcttgaagtggtggccta
actacggctacactagaagaacagtatttggtatctgcgctctgctgaagccagttaccttcggaaaaagagttg
gtagctcttgatccggcaaacaaaccaccgctggtagcggtggtttttttgtttgcaagcagcagattacgcgca
gaaaaaaaggatctcaagaagatcctttgatcttttctacggggtctgacgctcagtggaacgaaaactcacgtt
aagggattttggtcatgagattatcaaaaaggatcttcacctagatccttttaaattaaaaatgaagttttaaat

caatctaaagtatatatgagtaaacttggtctgacagttaccaatgcttaatcagtgaggcacctatctcagcga
tctgtctatttcgttcatccatagttgcctgactccccgtcgtgtagataactacgatacgggagggcttaccat
ctggccccagtgctgcaatgataccgcgagaaccacgctcaccggctccagatttatcagcaataaaccagccag
ccggaagggccgagcgcagaagtggtcctgcaactttatccgcctccatccagtctattaattgttgccgggaag
ctagagtaagtagttcgccagttaatagtttgcgcaacgttgttgccattgctacaggcatcgtggtgtcacgct
cgtcgtttggtatggcttcattcagctccggttcccaacgatcaaggcgagttacatgatcccccatgttgtgca
aaaaagcggttagctccttcggtcctccgatcgttgtcagaagtaagttggccgcagtgttatcactcatggtta
tggcagcactgcataattctcttactgtcatgccatccgtaagatgcttttctgtgactggtgagtactcaacca
agtcattctgagaatagtgtatgcggcgaccgagttgctcttgcccggcgtcaatacgggataataccgcgccac
atagcagaactttaaaagtgctcatcattggaaaacgttcttcggggcgaaaactctcaaggatcttaccgctgt
tgagatccagttcgatgtaacccactcgtgcacccaactgatcttcagcatcttttactttcaccagcgtttctg
ggtgagcaaaaacaggaaggcaaaatgccgcaaaaaagggaataagggcgacacggaaatgttgaatactcatac
tcttcctttttcaatattattgaagcatttatcagggttattgtctcatgagcggatacatatttgaatgtattt
agaaaaataaacaaataggggttccgcgcacatttccccgaaaagtgccacctaaattgtaagcgttaatatttt
gttaaaattcgcgttaaatttttgttaaatcagctcattttttaaccaataggccgaaatcggcaaaatccctta
taaatcaaaagaatagaccgagatagggttgagtggccgctacagggcgctcccattcgccattcaggctgcgca
actgttgggaagggcgtttcggtgcgggcctcttcgctattacgccagctggcgaaaggggatgtgctgcaagg
cgattaagttgggtaacgccagggttttcccagtcacgacgttgtaaaacgacggccagtgagcgcgacgtaata
cgactcactatagggcgaattggcggaaggccgtcaaggcctaggcgcgccagcattaattaatacgactcacta
tagggg

#### pCam$_{VU8}$-V$_{1K}$ (3315 bp)

#### Map

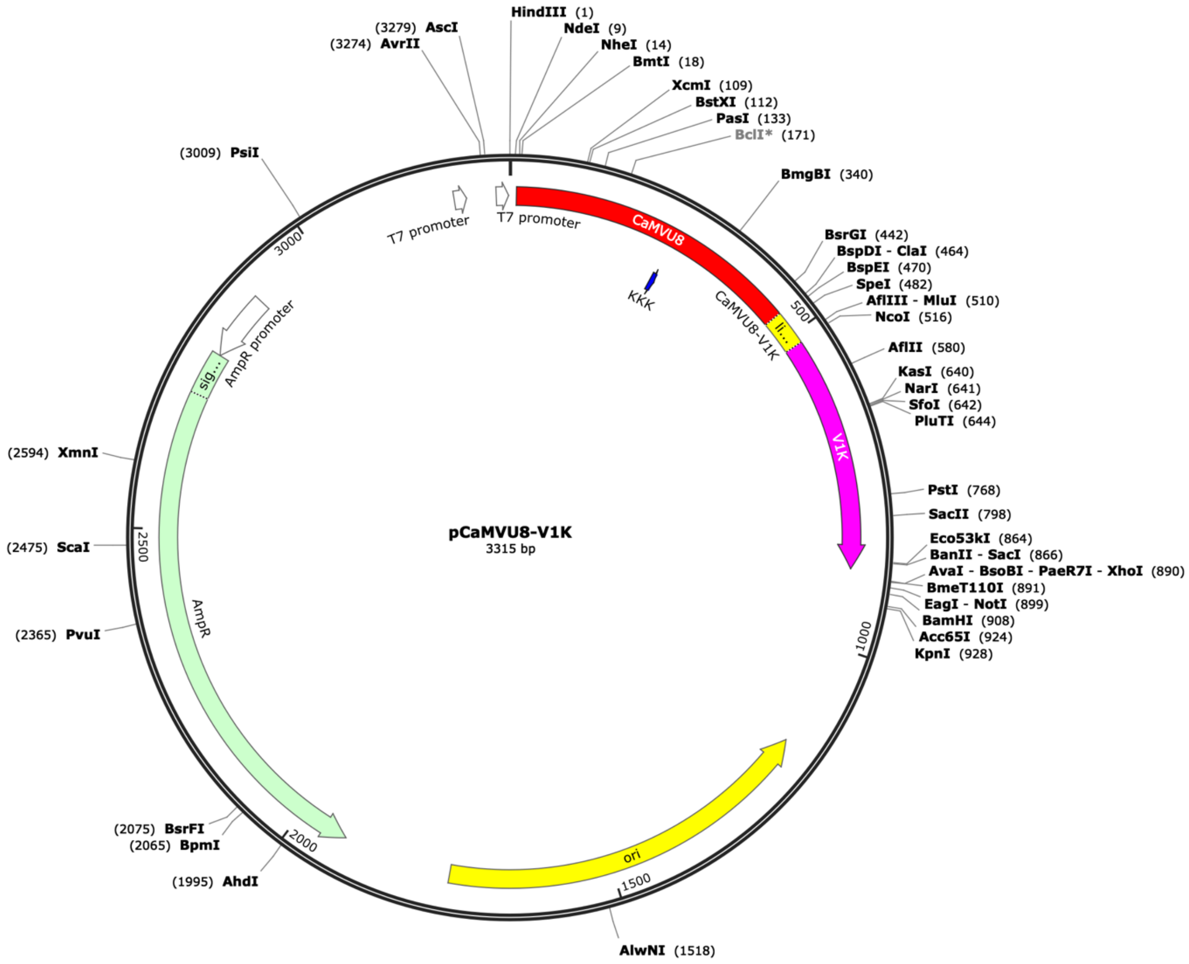


#### Full DNA sequence

aagcttgcATATGGCTAGCGCCGACCAGCTGACCGAGGAGCAGATCGCCGAGTTCAAGGAGGCGTTCTCCCTGTT
CGACAAGGACGGCGACGGCACCATCACCACCAAGGAGCTGGGCACCGTCATGCGGTCCCTGGGCCAGAACCCCAC
CGAGGCCGAGCTTCAGGACATGATCAACGAGGTCGACGCCGACGGCAACGGCACCATCGACTTCCCCGAATTCCT
GACCATGATGGCGCGCAAGATGAAGGACACCGATTCGAAGAAGAAGATCCGGGAGGCCTTCCGGGTCTTCGACAA
GGACGGCAACGGCTATATCTCCGCCGCCGAGCTGCGGCACGTCATGACCAACCTGGGCGAGAAGCTGACCGACGA
GGAGGTCGACGAGATGATCCGGGAGGCCGACATCGACGGCGACGGCCAGGTCAACTATGAGGAGTTTGTACAGAT
GATGACCGCCAAATCGATGTCCGGAGGTGGCACTAGTTACCCATACGATGTCCCAGATTACGCGTCCATGGCGCA
GGTTCAGCTTGTTGAATCTGGTGGTGCGCTGGTTCAGCCGGGTGGTTCTCTGCGCTTAAGCTGCGCGGCTTCCGG
TTTCCCGGTTAACCGTTACTCTATGCGTTGGTATCGTCAGGCGCCGGGTAAAGAACGTGAATGGGTTGCGGGTAT
GTCTTCTGCGGGTGACCGTTCTTCTTACGAAGACTCTGTTAAAGGTCGTTTCACCATCTCTCGTGACGACGCGCG
TAACACCGTTTACCTGCAGATGAACTCTCTGAAACCGGAAGACACCGCGGTTTACTACTGCAACGTTAACGTTGG
TTTCGAATACTGGGGTCAGGGCACCCAGGTTACCGTGAGCTCGAATTCATCGTAActaagtaatctcgagtagcg
gccgcttggatcccagaattcta**ggtacc**tcttaattaactggcctcatgggccttccgctcactgcccgctttc
cagtcgggaaacctgtcgtgccagctgcattaacatggtcatagctgtttccttgcgtattgggcgctctccgct
tcctcgctcactgactcgctgcgctcggtcgttcggtaaagcctggggtgcctaatgagcaaaaggccagcaaa
aggccaggaaccgtaaaaaggccgcgttgctggcgtttttccataggctccgcccccctgacgagcatcacaaaa
atcgacgctcaagtcagaggtggcgaaacccgacaggactataaagataccaggcgtttccccctggaagctccc
tcgtgcgctctcctgttccgaccctgccgcttaccggatacctgtccgcctttctcccttcgggaagcgtggcgc
tttctcatagctcacgctgtaggtatctcagttcggtgtaggtcgttcgctccaagctgggctgtgtgcacgaac
cccccgttcagcccgaccgctgcgccttatccggtaactatcgtcttgagtccaacccggtaagacacgacttat
cgccactggcagcagccactggtaacaggattagcagagcgaggtatgtaggcggtgctacagagttcttgaagt
ggtggcctaactacggctacactagaagaacagtatttggtatctgcgctctgctgaagccagttaccttcggaa
aaagagttggtagctcttgatccggcaaacaaaccaccgctggtagcggtggtttttttgtttgcaagcagcaga

ttacgcgcagaaaaaaaggatctcaagaagatcctttgatcttttctacggggtctgacgctcagtggaacgaaa
actcacgttaagggattttggtcatgagattatcaaaaaggatcttcacctagatccttttaaattaaaaatgaa
gttttaaatcaatctaaagtatatatgagtaaacttggtctgacagttaccaatgcttaatcagtgaggcaccta
tctcagcgatctgtctatttcgttcatccatagttgcctgactccccgtcgtgtagataactacgatacgggagg
gcttaccatctggccccagtgctgcaatgataccgcgagaaccacgctcaccggctccagatttatcagcaataa
accagccagccggaagggccgagcgcagaagtggtcctgcaactttatccgcctccatccagtctattaattgtt
gccgggaagctagagtaagtagttcgccagttaatagtttgcgcaacgttgttgccattgctacaggcatcgtgg
tgtcacgctcgtcgtttggtatggcttcattcagctccggttcccaacgatcaaggcgagttacatgatccccca
tgttgtgcaaaaaagcggttagctccttcggtcctccgatcgttgtcagaagtaagttggccgcagtgttatcac
tcatggttatggcagcactgcataattctcttactgtcatgccatccgtaagatgcttttctgtgactggtgagt
actcaaccaagtcattctgagaatagtgtatgcggcgaccgagttgctcttgcccggcgtcaatacgggataata
ccgcgccacatagcagaactttaaaagtgctcatcattggaaaacgttcttcggggcgaaaactctcaaggatct
taccgctgttgagatccagttcgatgtaacccactcgtgcacccaactgatcttcagcatcttttactttcacca
gcgtttctgggtgagcaaaaacaggaaggcaaaatgccgcaaaaaagggaataagggcgacacggaaatgttgaa
tactcatactcttcctttttcaatattattgaagcatttatcagggttattgtctcatgagcggatacatatttg
aatgtatttagaaaaataaacaaataggggttccgcgcacatttccccgaaaagtgccacctaaattgtaagcgt
taatattttgttaaaattcgcgttaaatttttgttaaatcagctcattttttaaccaataggccgaaatcggcaa
aatcccttataaatcaaaagaatagaccgagatagggttgagtggccgctacagggcgctcccattcgccattca
ggctgcgcaactgttgggaagggcgtttcggtgcgggcctcttcgctattacgccagctggcgaaagggggatgt
gctgcaaggcgattaagttgggtaacgccagggttttcccagtcacgacgttgtaaaacgacggccagtgagcgc
gacgtaatacgactcactatagggcgaattggcggaaggccgtcaaggcctaggcgcgccagcattaattaatac
gactcactataggggg

#### pCam$_{Cter}$-V$_{1K}$ (3087 bp)

#### Map

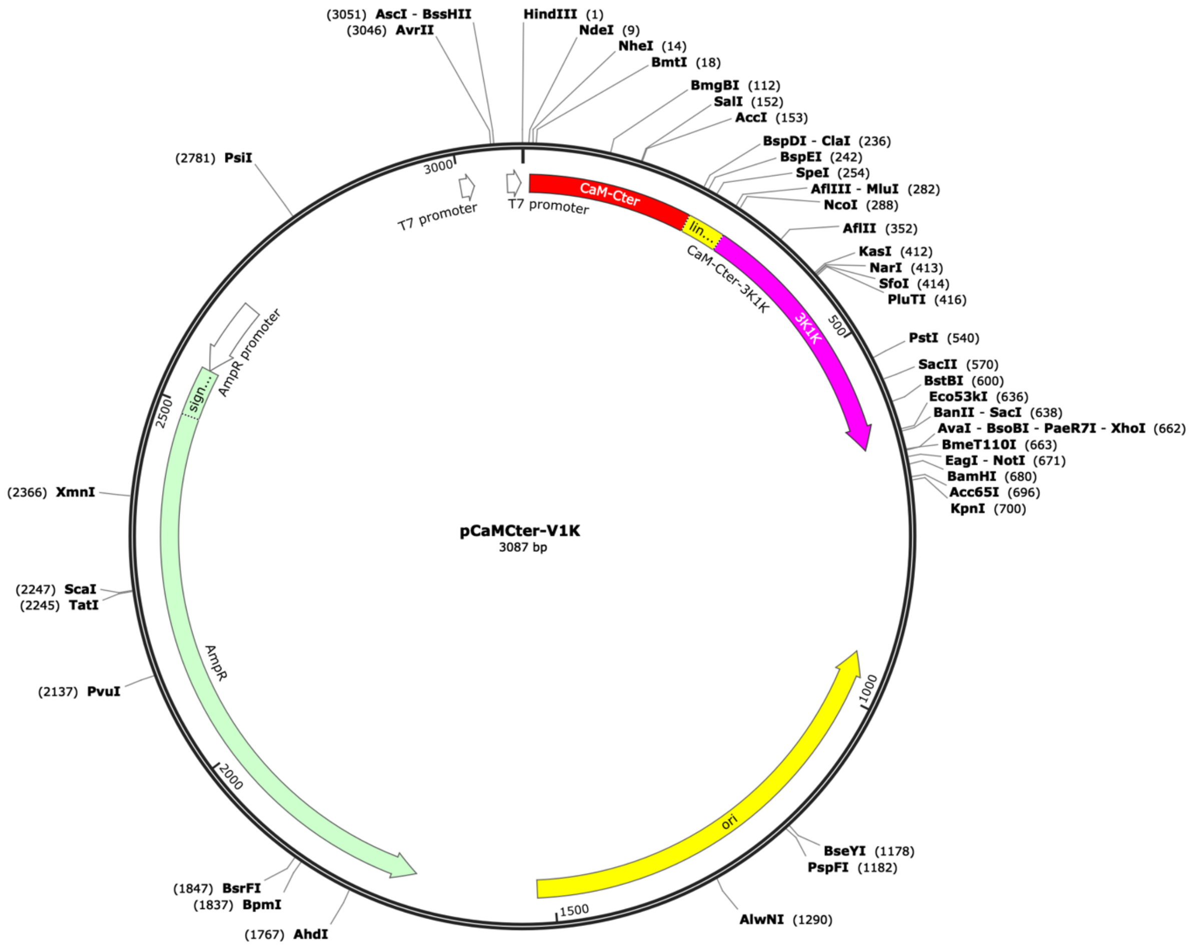


#### Full DNA sequence

**aagctt**gcatATGGCTAGCAAGGACACCGACTCCGAGGAGGAGATCCGGGAGGCCTTCCGGGTCTTCGACAAGGA
CGGCAACGGCTATATCTCCGCCGCCGAGCTGCGGCACGTCATGACCAACCTGGGCGAGAAGCTGACCGACGAGGA
GGTCGACGAGATGATCCGGGAGGCCGACATCGACGGCGACGGCCAGGTCAACTATGAGGAGTTCGTCCAGATGAT
GACCGCCAAATCGATGTCCGGAGGTGGCACTAGTTACCCATACGATGTCCCAGATTACGCGTCCATGGCGCAGGT
TCAGCTTGTTGAATCTGGTGGTGCGCTGGTTCAGCCGGGTGGTTCTCTGCGCTTAAGCTGCGCGGCTTCCGGTTT
CCCGGTTAACCGTTACTCTATGCGTTGGTATCGTCAGGCGCCGGGTAAAGAACGTGAATGGGTTGCGGGTATGTC
TTCTGCGGGTGACCGTTCTTCTTACGAAGACTCTGTTAAAGGTCGTTTCACCATCTCTCGTGACGACGCGCGTAA
CACCGTTTACCTGCAGATGAACTCTCTGAAACCGGAAGACACCGCGGTTTACTACTGCAACGTTAACGTTGGTTT
CGAATACTGGGGTCAGGGCACCCAGGTTACCGTGAGCTCGAATTCATCGTAActaagtaatctcgagtagcggcc
gcttggatcccagaattcta**ggtacc**tcttaattaactggcctcatgggccttccgctcactgcccgctttccag
tcgggaaacctgtcgtgccagctgcattaacatggtcatagctgtttccttgcgtattgggcgctctccgcttcc
tcgctcactgactcgctgcgctcggtcgttcgggtaaagcctggggtgcctaatgagcaaaaggccagcaaaagg
ccaggaaccgtaaaaaggccgcgttgctggcgtttttccataggctccgcccccctgacgagcatcacaaaaatc
gacgctcaagtcagaggtggcgaaacccgacaggactataaagataccaggcgtttccccctggaagctccctcg
tgcgctctcctgttccgaccctgccgcttaccggatacctgtccgcctttctcccttcgggaagcgtggcgcttt
ctcatagctcacgctgtaggtatctcagttcggtgtaggtcgttcgctccaagctgggctgtgtgcacgaacccc
ccgttcagcccgaccgctgcgccttatccggtaactatcgtcttgagtccaacccggtaagacacgacttatcgc
cactggcagcagccactggtaacaggattagcagagcgaggtatgtaggcggtgctacagagttcttgaagtggt
ggcctaactacggctacactagaagaacagtatttggtatctgcgctctgctgaagccagttaccttcggaaaaa
gagttggtagctcttgatccggcaaacaaaccaccgctggtagcggtggtttttttgtttgcaagcagcagatta
cgcgcagaaaaaaaggatctcaagaagatcctttgatcttttctacggggtctgacgctcagtggaacgaaaact

cacgttaagggattttggtcatgagattatcaaaaaggatcttcacctagatccttttaaattaaaaatgaagtt
ttaaatcaatctaaagtatatatgagtaaacttggtctgacagttaccaatgcttaatcagtgaggcacctatct
cagcgatctgtctatttcgttcatccatagttgcctgactccccgtcgtgtagataactacgatacgggagggct
taccatctggccccagtgctgcaatgataccgcgagaaccacgctcaccggctccagatttatcagcaataaacc
agccagccggaagggccgagcgcagaagtggtcctgcaactttatccgcctccatccagtctattaattgttgcc
gggaagctagagtaagtagttcgccagttaatagtttgcgcaacgttgttgccattgctacaggcatcgtggtgt
cacgctcgtcgtttggtatggcttcattcagctccggttcccaacgatcaaggcgagttacatgatcccccatgt
tgtgcaaaaaagcggttagctccttcggtcctccgatcgttgtcagaagtaagttggccgcagtgttatcactca
tggttatggcagcactgcataattctcttactgtcatgccatccgtaagatgcttttctgtgactggtgagtact
caaccaagtcattctgagaatagtgtatgcggcgaccgagttgctcttgcccggcgtcaatacgggataataccg
cgccacatagcagaactttaaaagtgctcatcattggaaaacgttcttcggggcgaaaactctcaaggatcttac
cgctgttgagatccagttcgatgtaacccactcgtgcacccaactgatcttcagcatcttttactttcaccagcg
tttctgggtgagcaaaaacaggaaggcaaaatgccgcaaaaaagggaataagggcgacacggaaatgttgaatac
tcatactcttcctttttcaatattattgaagcatttatcagggttattgtctcatgagcggatacatatttgaat
gtatttagaaaaataaacaaataggggttccgcgcacatttccccgaaaagtgccacctaaattgtaagcgttaa
tattttgttaaaattcgcgttaaatttttgttaaatcagctcattttttaaccaataggccgaaatcggcaaaat
cccttataaatcaaaagaatagaccgagatagggttgagtggccgctacagggcgctcccattcgccattcaggc
tgcgcaactgttgggaagggcgtttcggtgcgggcctcttcgctattacgccagctggcgaaagggggatgtgct
gcaaggcgattaagttgggtaacgccagggttttcccagtcacgacgttgtaaaacgacggccagtgagcgcgac
gtaatacgactcactatagggcgaattggcggaaggccgtcaaggcctaggcgcgccagcattaattaatacgac
tcactatagggg
